# Spatial overlap and phenotypic similarity predict hybridisation in birds

**DOI:** 10.64898/2026.10.21.759569

**Authors:** D. A. Nesbit, M. C. Cowen, G. F. Grether, R. Ligon, E. T. Miller, M. Brambilla, T. Chodkiewicz, B. Fontaine, A. Gamero, F. Jiguet, I.J. Øien, Å. Lindström, D. Massimino, D. Palm, B.G. Stokke, S. Trautmann, J. P. Drury

**Affiliations:** Department of Biosciences, Durham University, Stockton Road, Durham, DH1 3LE, United Kingdom; Department of Ecology C Evolutionary Biology, University of California, Los Angeles, United States of America; Cornell Lab of Ornithology, 159 Sapsucker Woods Rd., Ithaca, New York, 14850, USA; Department of Environmental Science and Policy, University of Milan, via Celoria 26, I-20133, Milan, Italy; Museum and Institute of Zoology PAS, OTOP BirdLife Poland, ul. Odrowąża 24, 05-270, Marki k., Warszawy, Poland; Muséum National d’Histoire Naturelle, 57 rue Cuvier, 75005, Paris, France; Czech Society for Ornithology, Na Bělidle 34, 150 00 Praha 5, Czechia; BirdLife Norway, Sandgata 30 B, NO-7012, Trondheim, Norway; Department of Biology, Division of Biodiversity and Evolution, Lund University, Ecology Building, S-223 62, Lund, Sweden; British Trust for Ornithology, The Nunnery, Thetford, Norfolk, IP24 2PU, United Kingdom; Danish Ornithological Society, Vesterbrogade 138-140, 1620, København V, Denmark; Norwegian Institute for Nature Research, P.O. Box 5685, Torgarden, 7485, Trondheim, Norway; DDA-Geschäftsstelle, An den Speichern 2, D-48157 Münster, Germany

**Keywords:** hybridisation, Aves, species recognition, speciation, species interactions

## Abstract

Hybridisation is widespread in birds and has important eco-evolutionary implications. Historically, much research has focused on the consequences of hybridisation. To date, however, far less attention has been paid to the causes of hybridisation and, in particular, why particular pairs of species form hybrids. Using large-scale comparative analyses, we find that similarity in plumage, vocalisations, and morphometric traits, as well as evolutionary relatedness, drive hybridisation in North American and European birds. In passerines, the avian order including songbirds, similarity in vocalisations and morphology are the key phenotypic predictors of hybridisation, whereas in non-passerines, species pairs with similar plumage and morphology are more likely to hybridise. We also find that the likelihood of hybrid formation increases with the magnitude of range and habitat overlap, demonstrating that hybridisation is not just a product of occasional contact between closely related species. Our findings have important conservation implications in this age of rapid anthropogenic change and species reassembly.

## Introduction

Growing molecular evidence of introgression between species has placed hybridisation at the centre of much research in ecology and evolutionary biology (Edelman and Mallet, 2021; Irwin and Schluter, 2022; Porretta and Canestrelli, 2023; Peñalba *et al*., 2024). For instance, though the study of hybrid zones has long been important to speciation research, widespread introgression and the discovery of hybrid speciation are making investigators re-evaluate how reticulate evolution can be accommodated in existing speciation and diversification research frameworks (Mallet, Besansky and Hahn, 2016; Barley *et al*., 2022). Similarly, conservation biologists are grappling with how to conserve species in the face of both naturally occurring and invasion-driven hybridisation (Fitzpatrick *et al*., 2015; Wayne and Shaffer, 2016), even considering hybridisation itself as a potential conservation tool (Chan, Hoffmann and van Oppen, 2019).

Historically, much research has focused on the consequences of hybridisation (e.g., lower hybrid fitness as a selective mechanism driving the evolution of reproductive isolation). To date, however, much less attention has been paid to the causes of hybridisation. At the species level, the propensity to hybridise can be predicted by particular traits (e.g., perenniality in plants (Mitchell *et al*., 2019), social bond length and migratory behaviour in birds (Leighton *et al*., 2021)). Yet, a critical open question is: what are the drivers for particular pairs of species to form hybrid offspring in the first place (Willis, Symula and Lovette, 2014; Crossman, Taylor and Barrett-Lennard, 2016; Brown *et al*., 2023)? Although genetic divergence is clearly an important factor (e.g., Price and Bouvier, 2002; Brown *et al*., 2023), other factors must also be important, as not all coexisting congeners hybridise. Mate recognition, or the behavioural response to the stimuli of potential mates (Mendelson, 2015), plays a primary role in interspecific mating and, therefore, is likely central to explaining which species pairs are likely to hybridise (Wallace, 1889). In animals, this hypothesis posits that hybridisation is more likely to occur between species that are behaviourally, visually, and vocally similar, which, due to phylogenetic conservatism, are often closely related (Price 2008).

Similarly, ecological factors such as niche overlap may influence hybridisation owing to increased encounter rates or shared selection pressures that influence hybrid viability (Edelman and Mallet, 2021; Porretta and Canestrelli, 2023). Currently, however, we lack a clear understanding of which of these factors are most likely to influence the probability of hybridisation, hampering our ability to predict which species are likely to hybridise in novel assemblages that are forming as a result of ongoing anthropogenic climate change and introduction to non-native areas (Parmesan, 2006; Elith, Kearney and Phillips, 2010; Chen *et al*., 2011; HilleRisLambers *et al*., 2013; Elmhagen *et al*., 2015; Freeman *et al*., 2018; Avidad, Clavero and Viana, 2025). Indeed, the fate of such range shifts may, in part, be determined by reproductive interactions with heterospecifics (Pfennig, Kelly and Pierce, 2016; Patterson and Drury, 2023).

Hybridisation is widespread in birds (Grant and Grant, 1992; McCarthy, 2006; Ottenburghs, 2023), with an estimated 9-19% of species thought to hybridise with at least one other species (Aliabadian and Nijman, 2007). Here, we combine phylogenetic and phenotypic data across 812 species of birds with fine-scale occurrence data to test the hypothesis that traits likely to be involved in mate recognition predict variation in the propensity to hybridise. We find that increasing similarity in traits typically associated with mate choice—vocalisations and plumage traits—corresponds to an increased probability of hybridisation. Surprisingly, we also find strong evidence that similarity in morphology is a key predictor for hybridisation. Moreover, we consistently find that spatial overlap at both range-wide and habitat scales is associated with an increased likelihood of hybridisation. Our findings have important implications for both speciation research and conservation in the face of ongoing range shifts.

## Results

In our dataset of North American and European birds, 8.8% of co-occurring species pairs from the same family have been reported to hybridise (Fig. 1, Supplementary Table 13). To identify factors that influence the formation of hybrids, we modelled hybridisation using phylogenetic logistic mixed models. In addition to a full model with all species pairs, we fit separate models for passerines and non-passerines due to potential differences in the relative importance of mate recognition traits in these groups, vocalisations being more important in mate recognition in passerines (the order including songbirds), for instance (Arato and Fitch, 2021).

**Figure 1.**
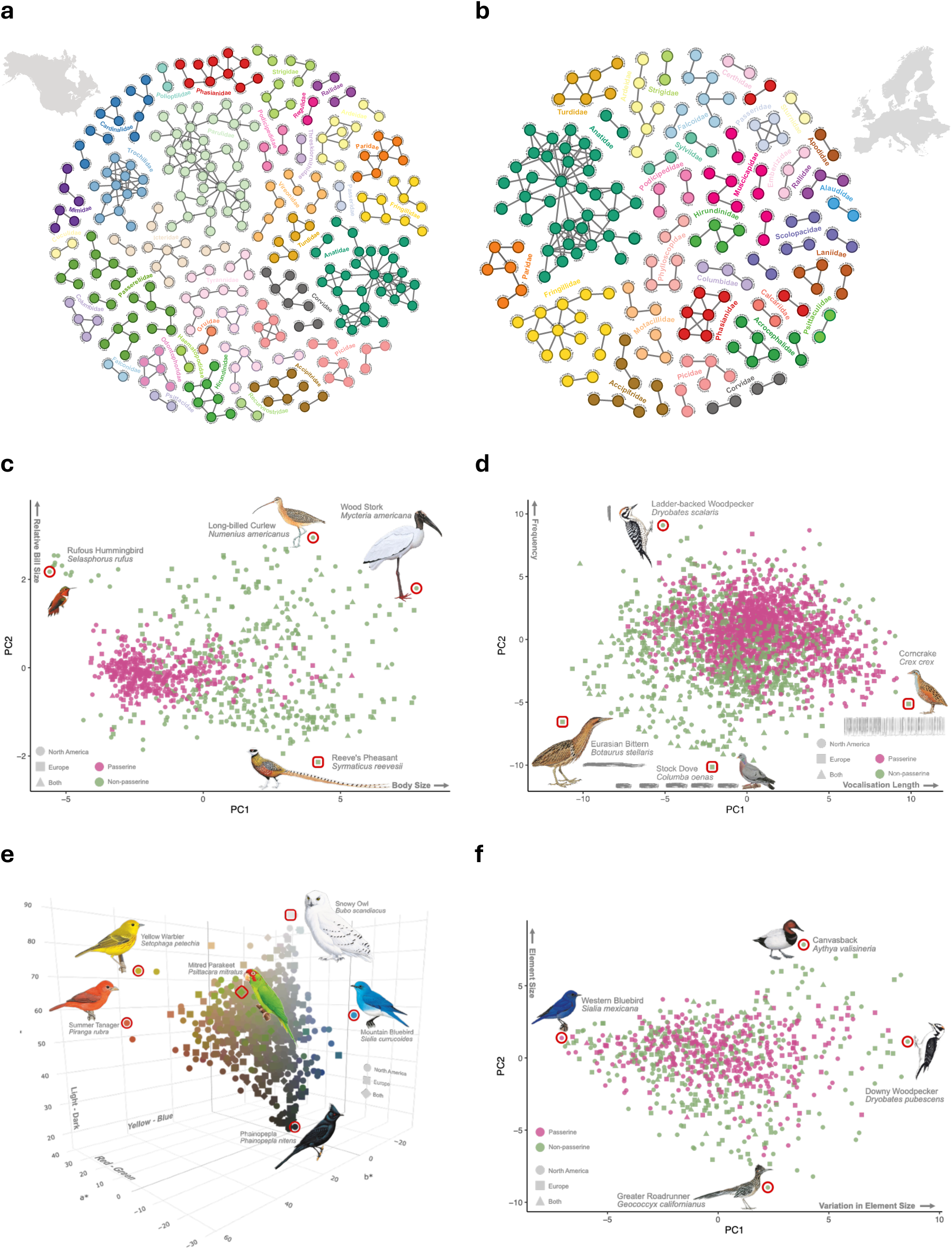
**a.** Network of North American hybridisation. Nodes represent species (n=285) (coloured by family), and lines indicate recorded hybridisation in the wild (n=309). Non-hybridising pairs are not included**. b.** Network of European hybridisation. Nodes represent species (n=188) (coloured by family), and lines indicate recorded hybridisation in the wild (n=208). Non-hybridising pairs are not included. **c.** The first two axes of the principal component analysis (PCA) representing morphospace for all birds in the dataset. **d.** The first two axes of the PCA representing vocalisations. Sonogram illustrations are not to scale. **e.** CIELAB colour space representing the average colour of birds in the dataset. **f.** The first two axes of the PCA representing plumage pattern variation. Illustrations: Cornell Lab of Ornithology | Birds of the World (https://doi.org/10.2173/bow) [Reeves Pheasant: Antoni Lombarte, Wood Stork: Francesc Jutglar, Rufous Hummingbird: Antoni Llobet, Long-billed Curlew: Francesc Jutglar, Ladder-backed Woodpecker: Ian Willis, Corncrake: Hilary Burn, Stock Dove: Jan Wilczur, Eurasian Bittern: Francesc Jutglar, Canvasback: Àngels Jutglar, Western Bluebird: Ian Willis, Downy Woodpecker: Ian Willis, Greater Roadrunner: Tim Worfolk, Mountain Bluebird: Ian Willis, Phainopepla: Doug Pratt, Summer Tanager: Hilary Burn, Yellow Warbler: David Quinn, Snowy Owl: Tim Worfolk, Mitred Parakeet: Àngels Jutglar].

### Divergence time between hybridising species

Consistent with a key role of genomic divergence in hybrid formation, more distantly related species are less likely to hybridise (Figure 2a-b, Supplementary Tables 2-4). Nevertheless, the modelled probability of hybrid formation was high (>0.5) until ∼10My of separation between species, and the median time to the most recent common ancestor of hybridising pairs was 5.66 My. The oldest hybridising pair in our dataset was Snowy Egret (*Egretta thula*) and Western Cattle Egret (*Bubulcus ibis*), which shared a common ancestor 28.8 Ma.

**Figure 2.**
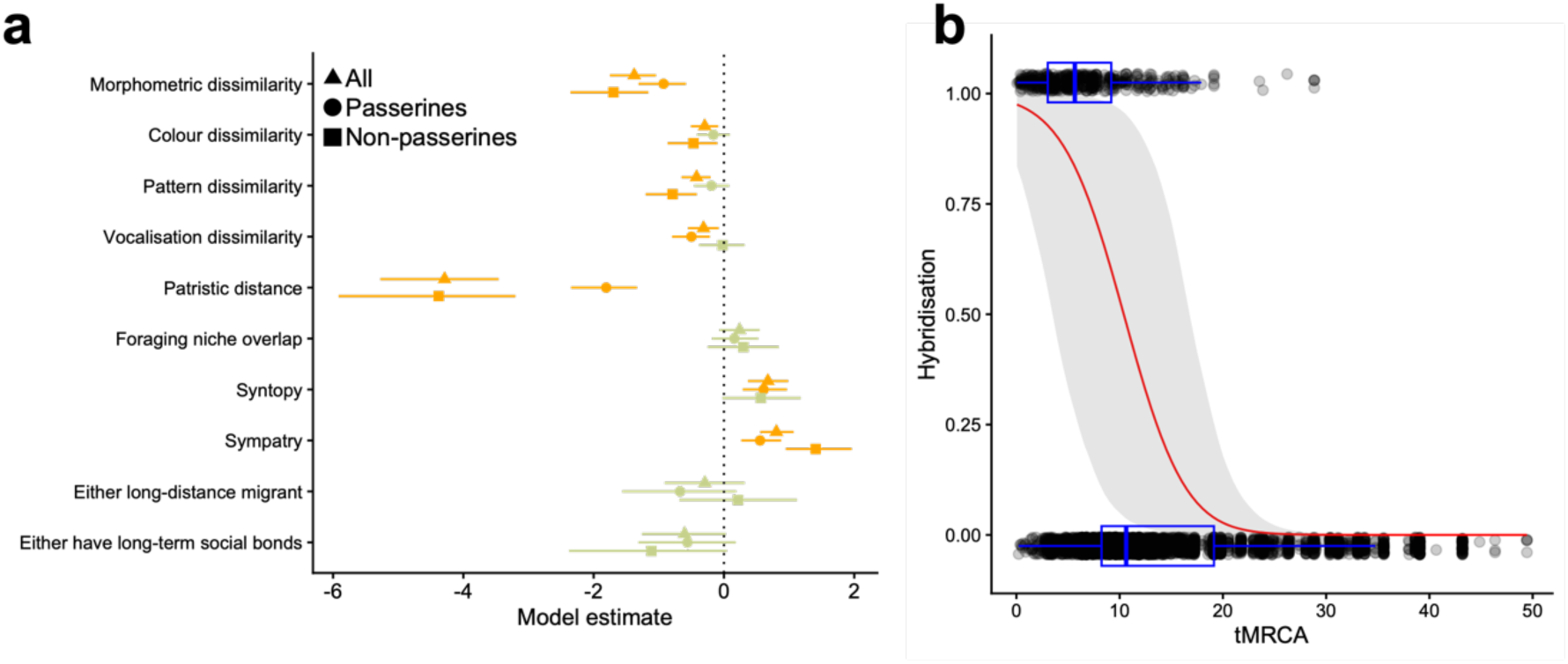
**a.** Predictors of hybridisation. Plotted values are coefficient estimates from phylogenetic generalised linear mixed models with hybridisation as the response variable. Points correspond to the median and error bars represent the 95% credibility interval from four combined MCMC chains. Orange points indicate fixed effects with estimates with 95% credibility intervals that exclude 0. **b.** Relationship between hybrid formation and time since the most recent common ancestor (tMRCA). Each point (jittered vertically for clarity) represents a single species pair. The red line depicts the posterior mean predicted probability of hybridisation, with 95% Bayesian credible intervals (in grey) derived from the posterior distribution.

### Phenotypically similar species are more likely to form hybrids

Controlling for the effect of divergence time, we found that phenotypic similarity between species increases the likelihood of hybrid formation. Firstly, we found that morphometric dissimilarity strongly reduced the likelihood of hybridisation across all analyses (Figure 2a, Supplementary Tables 2-4). In the full model, both plumage and vocalisation similarity increased the likelihood of hybridisation. However, these effects were largely driven by different factors in non-passerines and passerines. Specifically, plumage colour and pattern dissimilarity—but not vocalisation dissimilarity—reduced the likelihood of hybridisation in non-passerines (Figure 2a, Supplementary Tables 2-4). Contrastingly, vocalisation dissimilarity—but not plumage colour or pattern dissimilarity—reduced the likelihood of hybridisation in passerines (Figure 2a, Supplementary Tables 2-4).

In the full model and the passerine model, the relationship between phenotypic similarity and hybridisation did not depend on patristic distance (i.e., the evolutionary divergence time separating the two species in a pair). In the non-passerine model, however, the relationship between vocalisation dissimilarity and hybridisation depended on patristic distance (Supplementary Tables 5-6, Supplementary Figure 1). Specifically, at small patristic distances, hybridisation was less likely between species with similar vocalisations.

### Habitat, range, and foraging niche overlap inffuence the likelihood of hybridisation

Although our analyses were restricted to co-occurring species, we found that the extent of fine-scale habitat overlap (syntopy) and range overlap (sympatry) also influenced the probability of hybrid formation, with increasing values of both indices corresponding to an increased likelihood of hybridisation (Figure 2a, Supplementary Tables 2-6). Foraging niche overlap did not influence the likelihood of hybridisation in most models. However, an interaction between foraging niche overlap and sympatry in non-passerines demonstrates that hybrid formation is more likely between species pairs with high syntopy and low foraging niche overlap (Supplementary Tables 5-6, Supplementary Figure 2).

In the full model and the passerine model, but not the non-passerine model, the relationship between spatial overlap and hybridisation depended on patristic distance. In particular, an interaction between sympatry and patristic distance in the full model indicates that, while recently diverged taxa are predicted to form hybrids across a wide range of sympatry values, more distantly related pairs are only predicted to hybridise when their ranges overlap substantially (Supplementary Table 6, Supplementary Figure 3). Similarly, in the passerine model, whereas syntopy does not influence hybridisation between recently diverged species pairs, more distantly related pairs only hybridise at high values of fine-scale habitat overlap (Supplementary Table 7, Supplementary Figure 4).

### Species-specific traits

To control for species-specific traits that are known to influence hybridisation (Leighton *et al*., 2021), we included variables associated with migratory strategy and social bond length in models. Neither the presence of a migratory species within a pair, nor social bond length influenced the likelihood of hybridisation (Figure 2a, Supplementary Tables 2-4).

### Effect of evolutionary timescales

In addition to the models with all species pairs, we fit models restricted to only those species pairs which share a common ancestor <5 or <10 Ma (Supplementary Table 9). The factors predicting hybridisation in these models are largely similar to those of the full model, though among the most recently diverged species, morphometric dissimilarity and pattern dissimilarity no longer emerge as predictors of hybridisation (Supplementary Table 9). Similarly, in the analyses of species pairs separated by <10Ma, vocalisation dissimilarity was no longer predictive of hybridisation.

### Hybrid frequency

To determine how the frequency of hybridisation is influenced by the factors identified above, we fit separate models to determine which factors predict frequent hybridisation (i.e., ‘extensive’ and ‘ongoing’ hybridisation, sensu McCarthy (2006), Supplementary Table 10). We found that, as with our main analyses, morphology, colour dissimilarity, patristic distance, and syntopy were key predictors of frequent hybridisation (Supplementary Table 11), though plumage pattern, vocalisation dissimilarity, and sympatry no longer emerged as significant predictors of hybridisation. When we restricted analyses to species pairs which form hybrids, we found that frequently hybridising species pairs were even more similar in morphology and colour and more recently diverged than other hybridising pairs, but less sympatric than other hybridising pairs (Supplementary Table 12).

## Discussion

We found that, in addition to the well-known relationship between hybrid formation and divergence time, spatial overlap and phenotypic similarity are key predictors of hybridisation in birds. Our analyses showed that hybridisation is more likely between species with greater range (sympatry) and fine-scale habitat (syntopy) overlap. Indeed, we find that fine-scale habitat overlap predicted hybridisation even among more recently diverged species pairs and in analyses of frequently hybridising pairs. Such a pattern may reflect the opportunity for species to encounter one another and form hybrid offspring. Yet, much theoretical literature suggests that hybridisation can generate Allee effects that constrain range dynamics—a prediction which is supported in many empirical cases (Patterson and Drury, 2023). In line with this prediction, we found that frequently hybridising species pairs, while not differing in range overlap from non-hybridising pairs, tend to exhibit lower range-wide overlap than less extensively hybridising pairs. Similarly, in some analyses, the influence of range overlap and fine-scale habitat overlap on hybridisation increases with divergence time, such that more distantly related species pairs are only likely to hybridise at high levels of spatial overlap. Nevertheless, our results underscore the need for further research on the interactions between hybridisation and range dynamics to clarify the circumstances under which hybridisation may facilitate, rather than constrain, range expansion (Pfennig, Kelly and Pierce, 2016).

Consistent with the hypothesis that similarity in traits used in mate recognition influences the likelihood of interspecific mating, we found that similarity in plumage and vocalisations both influence the likelihood of hybrid formation. The true direction of causality might be the reverse in some species pairs (Supplementary Figure 5), either because of convergent agonistic character displacement (ACD) (Grether et al. 2009), convergent selection related to adaptation for signal transmission in similar habitats (Morton 1975), or as a result of introgression. In birds, comparative analyses often find exaggerated signal and morphological divergence between sympatric pairs (Martin, Montgomerie and Lougheed, 2010; Anderson and Weir, 2021), suggesting convergence in general is rare relative to rates of hybridisation. Moreover, our analyses control for niche and fine-scale habitat overlap, which should eliminate any confounding owing to phenotypic convergence in species that share similar environments (e.g., adaptation to similar signal transmission environments (Morton, 1975) or convergent ACD driven by resource overlap (Grether *et al*., 2009; Grether and Okamoto, 2022)). Convergent ACD could also result from interspecific mate competition (Payne, 1980; Drury, Cowen and Grether, 2020). However, convergent ACD requires higher rates of interspecific mating than is thought to occur between most hybridising species, and thus we consider it very unlikely that the overall effects of plumage and vocalisation similarity in our analyses are products of that evolutionary process. An additional, non-mutually exclusive hypothesis for the relationship between phenotypic similarity and hybridisation is that trait convergence is also a consequence of hybridisation, owing to introgression of alleles that code for signal and morphological traits. Analyses of hybrid zones commonly report that clines are much steeper for plumage or song traits than for background genomic variation (Toews *et al*., 2016; Kenyon *et al*., 2017; Schield *et al*., 2024), indicating that loci coding for signal traits may be less likely to be introgressed than other loci. Further genomic analyses and functional mapping of loci that are exchanged between hybridising pairs—beyond the sister taxa that are commonly targeted for such cline analyses—combined with modelling of ancient and contemporary migration, are key to determining the relative role of phenotypic similarity driving hybridisation versus the converse, as well as any positive feedbacks between these two.

The relationship between plumage and vocalisation similarity and hybridisation may indicate that ‘errors’ in mate recognition drive hybridisation (a common assumption of reproductive character displacement hypotheses (Qvarnström *et al*., 2006; Rosenthal, 2013; Willis, 2013)). With respect to vocalisations, most passerines in our dataset are oscines (92%), which learn their songs, generating the additional possibility of mistakes in learning that could explain the relationship between vocalisations and hybridisation in this group. Given the limited number of suboscines— which have innate songs (Beecher and Brenowitz, 2005)—in our dataset, we could not conduct a robust test of this hypothesis, but further analyses expanding beyond northern temperate regions, where suboscines are scarce, would shed further light on the role of song learning on patterns of hybridisation.

An alternative hypothesis is that the relationship between signal similarity and hybridisation reflects a role of preference for novelty or sensory bias in sexual signals (Ryan, 2018; Broder *et al*., 2021). Under this hypothesis, rather than being unable to distinguish between conspecific and heterospecific signals, choosing individuals can perceive salient differences between stimuli and assess them favourably against an underlying neurological template, or mate recognition function (Mendelson, 2015).

Vocalisations and plumage often evolve at much faster rates than morphological traits (Matysioková *et al*., 2017; Drury *et al*., 2018); in oscine passerines, moreover, song learning generates the possibility of rapid cultural evolution (Williams *et al*., 2022). This means that, on average, divergence in reproductive traits should outpace divergence in other traits, and indeed, even small differences are detectable to birds (Brambilla *et al*., 2008; Seddon and Tobias, 2010). Under this alternative hypothesis, hybridisation is the outcome of choosing individuals preferring heterospecific signals over conspecific signals owing to their novelty, as long as those traits do not fall outside of the range of their mate recognition function (hence, in a large comparative context, appearing as similarity in mate recognition traits). Although low hybrid fitness may eventually drive reinforcement, it is also possible that hybridisation is adaptive in some instances, driving interspecific sexual selection (Chen and Pfennig, 2020). Distinguishing between these hypotheses across different hybridising species pairs is beyond the scope of our analyses, and further research characterising the mate recognition functions of choosing individuals would shed further light on the relationship between signal similarity and hybrid formation.

While our analyses point to a key role for traits involved in mate recognition in hybridisation, we found, surprisingly, that morphometric similarity was the strongest predictor of hybridisation. However, in analyses of species pairs separated by <5 Ma, morphometric similarity was no longer predictive of hybridisation. On the one hand, this is what we would expect if rare, distantly related hybrids were driving this result. Yet, on the other hand, we find that morphometric similarity is an important predictor in our analyses of frequently hybridising species pairs. Similarly, we find that there is no support for an interaction between evolutionary divergence (patristic distance) and morphological divergence.

It is also possible that the relationship between morphometric similarity and hybridisation reflects an underlying correlation between morphology and some other, unmeasured ecological or behavioural trait. For instance, the relationship between morphology and hybridisation could reflect convergent evolution of species that share similar niches (though this relationship is still detected in our models after controlling for niche and fine-scale habitat overlap). Nevertheless, the emergence of morphometric similarity across analyses could indicate a role for mechanical isolation and/or an underappreciated role for body size and shape and locomotory behaviour in mate recognition (Ratciliffe and Grant, 1983).

Our analyses focus on the formation of viable hybrids. Characterising the relationship between our set of predictor variables and other forms of reproductive interference (e.g., interspecific courtship, mixed-species mating that does not result in viable hybrids), hybrid fertility, and the frequency of hybridisation are important avenues for future research.

In the context of conservation, hybridisation can be problematic, particularly when one or both hybridising species are rare (Rhymer and Simberloff, 1996; Simberloff, 1996). Hybridisation can lead to ‘erosion’ of the genome, genetic swamping and hybrid swarms (Rhymer and Simberloff, 1996; Todesco *et al*., 2016; Wells *et al*., 2019), and when species are on the brink of extinction, could prevent recovery and drive them towards extinction (Rhymer and Simberloff, 1996; Wallis, 1999; Wolf, Takebayashi and Rieseberg, 2001; Muñoz-Fuentes *et al*., 2007). Indeed, hybridisation has been identified as a conservation concern for several species in our dataset (e.g., American black ducks [*Anas rubripes*] (Mank, Carlson and Brittingham, 2004), White-headed Ducks [*Oxyura leucocephala*] (Muñoz-Fuentes *et al*., 2007), Golden-Winged Warblers [*Vermivora chrysoptera*] (Toews *et al*., 2016)). Our results show that it should be possible, in principle, to make predictions about the likelihood that species which do not currently encounter one another will form hybrids. Given the ongoing reshuffling of Earth’s biota, predicting the risk of hybridisation in novel avian assemblages arising as a result of climate change and human-mediated changes has the potential to support conservation interventions, identify areas and species groups where investigating the magnitude and consequences of hybridisation is more urgently needed, and shape new research avenues.

## Methods

### Creating species lists

We focused on North American and European species pairs owing to the research bias in ornithology towards northern temperate regions (Reddy, 2014; Titley, Snaddon and Turner, 2017). In other words, by focusing on these regions, we assume that, for species in our dataset, a lack of reported hybridisation is not simply a reflection of a lack of observation.

For North American species pairs, a list of species breeding within the contiguous United States was created using the American Birding Association (ABA) checklist codes (Pyle *et al*., 2021) and the eBird ‘Lower 48’ regional checklist (Sullivan *et al*., 2009). Our species classifications followed the 2021 eBird taxonomy (Clements *et al*., 2021). Only ABA-designated Code-1 and Code-2 species (those which regularly occur and breed within the ABA area) were included to exclude rare vagrants and non-breeders. Range maps from Birds of the World were consulted to ensure species bred within the contiguous United States; 64 non-breeders were removed at this stage (Billerman *et al*., 2022). Non-native species were included if they occurred on the ABA list or the United States Register of Introduced and Invasive Species (US-RIIS) (Simpson, Blake and Dorado, 2021). While inclusion on these lists depends on different criteria, it requires species to be established and breeding, which is decided by expert panels. 8 non-natives were included only on the ABA list, one was on the US-RIIS list only, and 22 were shared by both lists. As a result of their pelagic habits and underreporting using traditional, terrestrial survey methods, pelagic seabirds (larids, alcids, procellariiforms) were excluded from the species list. This left 524 species that breed within the contiguous United States.

Our European species list was based on those present in the Pan-European Common Bird Monitoring Scheme (PECBMS) dataset (Brlík *et al*., 2021). The countries/schemes with available survey data were Austria, Brussels, Catalonia, Denmark, France, Germany, Ireland, Italy, Netherlands, Norway, Poland, Sweden, United Kingdom and Wallonia. Non-native species present in the PECBMS data were included, and Birds of the World accounts (Billerman *et al*., 2022) and eBird range maps (Sullivan *et al*., 2009) were consulted to confirm that their established non-native ranges fell within the study area. Again, pelagic seabirds were removed from the species list. A total of 386 European species remained.

### Species-pair dataset

Three separate sources were consulted to identify hybridising pairs in our species list. The first source was the Handbook of Avian Hybrids of the World (McCarthy, 2006). The second was the 2021 eBird Taxonomy (Clements *et al*., 2021), which includes hybrid taxa that are distinct enough to be recognised in the field by birders. Lastly, a literature search was carried out using Web of Science (webofscience.com), utilising species current and historical vernacular and scientific names and Boolean search terms to encompass the terms ‘hybrid’, ‘introgression’, ‘reproductive interference’, ‘mixed species pair’, ‘cross-species mating’ and their associated derivations. Only articles published post-2000 were searched to avoid duplication of effort with McCarthy (2006). Species pairs, therefore, had a maximum of three classifications of hybridisation, one from each source. Only pairs with convincing evidence of natural hybridisation were included in this pair list. Thus, only species pairs that produced hybrid offspring (i.e., not just cross-species mating) were scored as hybridising, though this definition also includes species pairs that produce infertile hybrids. McCarthy (2006) classified records as no doubt, some doubt and marked doubt; only records with no doubt were included in our analyses. Similarly, from the literature search, only records corroborated in peer-reviewed literature were included; anecdotal or unconfirmed hybrids were ignored. While we have less certainty of hybrids from the eBird taxonomy, their inclusion is the result of expert review (Justyn, Callaghan and Hill, 2020). There were 309 North American (Figure 1a) and 208 European (Figure 1b) species pairs with convincing evidence of natural hybridisation from at least one source (Supplementary Table 13).

The hybrid pairs included in the above analyses include all species pairs that produce hybrid offspring, even if such hybridisation is infrequent. To examine whether frequent hybridisation is influenced by distinct factors, we first made a list of species pairs with ‘extensive’ (n = 61) and ‘ongoing’ (n = 53) hybrid records using classifications from McCarthy (2006) (Supplementary Table 10). We then fit phylogenetic logistic mixed models, as above, with ‘extensive + ongoing’ = 1 and all other pairs = 0. Finally, we fit an additional set of models on hybrid pairs only to determine whether frequently hybridising pairs differ in the extent to which predictor variables influence the likelihood of hybridisation.

To create a comparison group comprised of species pairs that have not been reported to hybridise, every species on the species list was paired with every other member of its own family with which it co-occurs (interfamilial hybridisation is rare (McCarthy, 2006))—that is, species pairs that encounter one another and could theoretically hybridise but have not been reported to. BirdLife International range maps (BirdLife International and Handbook of the Birds of the World, 2021) were used to calculate range overlap. Similarly, the number of North American Breeding Bird Survey (BBS) (Pardieck *et al*., 2020) routes and stops shared by a pair in any year between 1976 and 2019, or the number of PECBMS sites at which species co-occurred between 1976 and 2019, was determined for North American and European species pairs, respectively. If a pair had any overlap in either method and did not hybridise, they were considered to be potentially but non-hybridising (*n* = 5333 pairs in our comparison group).

Our final species-pair dataset comprised 5850 pairs, of which 517 hybridise (Supplementary Table 13). When creating species pairs, species that were the sole members of their families in our species lists were removed as they lacked confamilials to pair them with (n=49). Our dataset therefore contained 503 North American species, 358 European species and 49 found on both continents.

### Predictors of Hybridisation

#### Morphological dissimilarity

Nine morphometric traits (beak length: culmen-to-tip, beak length: nares-to-tip, beak width, beak depth, tarsus length, wing length, secondary length, tail length and mass) were extracted from the AVONET dataset for each species (Tobias *et al*., 2022). These measurements were averaged from several individuals of both sexes (and individuals of unknown sex). For the species included in our analyses (*n* = 812), the mean number of individuals was 11.8 (minimum 1, maximum 123). The mean number of male, female and unknown sex specimens per species was 6.1, 4.5 and 1.2 individuals, respectively. These variables were log-transformed, centred and scaled, and analysed in a standard principal components analysis (PCA) as in Pigot *et al*. (2020)(Figure 1c). The Euclidean distance between each species pair in the resulting PC space was calculated to quantify morphometric dissimilarity.

#### Colour and pattern dissimilarity

Using methods developed by Miller *et al*. (2019), we utilised illustrations from Birds of the World (Billerman *et al*., 2022) to quantify dissimilarity in breeding plumage colour and pattern within pairs. Full body plates of males, not in flight (except swifts, which do not have perched illustrations, but were still included in analyses), of the subspecies that occurs in the contiguous United States or Western Europe were used. If multiple subspecies occur in the study area, a random number generator was used to select one. If the subspecies that occurs within the study area did not have an associated illustration, we chose the most closely related subspecies with an illustration. In sexually dichromatic species, we quantified plumage colouration in males, as females are generally considered to use male plumage colouration in mating decisions (Darwin, 1871; Hill, 2006). We acknowledge, however, that research into female sexual selection has long been neglected and that examples of male mate choice and mutual mate choice abound (Ah-King, 2022) and that, in some clades, male mate recognition can drive the occurrence of interspecific mating (Rohwer *et al*., 2022). Using the program GIMP (The GIMP Development Team, 2024), the background and legs were removed from the illustrations. The illustrations were then scaled so that they were approximately the same size. While in some species, legs may be an important feature in mate recognition (Iverson and Karubian, 2017), for these analyses, they were removed to enable illustrations to be scaled more accurately.

To quantify plumage colour dissimilarity between species pairs in our dataset, we used ImageJ (Schindelin *et al*., 2012) and macros developed by Miller *et al*. (2019). 1000 random points from across each illustration were sampled, and the Red (R), Green (G), and Blue (B) colour coordinates were extracted from each point and converted to perceptually uniform CIELAB colour coordinates (Figure 1e). For each pair, the 1000 colour coordinates of each species were plotted in CIELAB space, and the average Mahalanobis distance between all the points of a pair was calculated to provide a measure of colour dissimilarity.

To quantify plumage pattern similarity between species pairs in our dataset, we split illustrations into their corresponding RGB channels (following Miller *et al*. 2019). Pattern analysis was conducted on the G channel only, as it corresponds more closely to avian luminance channels associated with pattern processing (Stevens and Cuthill, 2006). Granularity-based pattern analysis was carried out using ImageJ (Schindelin *et al*., 2012) with macros from Miller *et al*. (2019). Illustrations were band-pass filtered using the Fast Fourier Transform into 17 granularity bands corresponding to spatial frequencies ranging from 2 to 512 pixels, increasing by multiples of √2. The ‘energy’ at each scale was quantified as the standard deviation of filtered pixel values and corresponds to the contribution to overall appearance from pattern elements of that size. For example, high energy at a high granularity band (e.g., 2 pixels) corresponds to many small pattern elements, and low energy at that band corresponds to few small pattern elements. Alternatively, high energy at a low granularity band (e.g., 512 pixels) corresponds to many large pattern elements, and low energy would correspond to few large elements. Each species, therefore, has 17 values of pattern energy, one for each granularity band. These 17 energy values were analysed with a PCA (Figure 1f), and the Euclidean distance between species within each pair was calculated to provide a measure of pattern dissimilarity.

We acknowledge that this approach is conservative, as differences between species in ultraviolet plumage colouration will not be detected, nor will clade-wide variation in salient, patch-specific differences (e.g., differences in crown colouration). Nevertheless, we argue that our approach is useful for comparative analyses such as those we present here.

#### Vocalisation Dissimilarity

Following sensitivity analyses carried out by Cowen, Drury and Grether (2020), Birds of the World (Billerman *et al*., 2022) species accounts were consulted to determine species vocal repertoire size. If species had fewer than four courtship/territorial vocalisation types, they were considered to have a small repertoire, and two sound files were downloaded from xeno-canto (xeno-canto.org) (Supplementary Table 14). Four sound files were downloaded for those with large repertoires (four or more vocalisation types). We aimed to download the highest quality sound files recorded in the United States and Europe. However, this was not always possible, in which case recordings were requested from the Macaulay Library at the Cornell Lab of Ornithology (www.macaulaylibrary.org). In cases where the sex of the vocalising individual was reported, we used recordings of individual males. Given the difficulty in disentangling the vocalisations of individuals of lekking species, e.g. *Tympanuchus* grouse, we removed pairs containing lekking species from analyses (n = 60). Further, to compare like-for-like, non-vocal sounds such as the drumming of woodpeckers, winnowing of snipe and dive displays of hummingbirds were not included in analyses despite their role in species recognition and mate selection, with vocal sounds used instead (Hurly, Scott and Healy, 2001; Miles *et al*., 2018). Despite documented hybridisation, Black Storks (*Ciconia nigra*) and White Storks (*C. ciconia*) were removed from European analyses as the only available sounds for White Storks were non-vocal bill clattering. The hissing exhalations of otherwise mute Cathartid vultures and Muscovy Ducks (*Cairina moschata*) were also removed from analyses. In total, we removed 103 species pairs due to a lack of vocalisation data.

Sound files were cleaned using noise reduction in Audacity (version 3.7.1, audacityteam.org) (starting values of Noise Reduction (dB) = 12, Sensitivity = 10, Frequency Smoothing = 0, but were altered depending on the specific file) and were trimmed to encompass a single vocalisation (songs in passerines and calls in non-passerines). We defined a single vocalisation as vocal elements separated from one another by less than one second. Once trimmed, sound files were saved as 44100Hz, 16-bit, mono .wav files. They were then all normalized to standardize their amplitude. Raven Lite (version 2.0.5, Charif, Ponirakis and Krein, 2006) was then used to create selection tables for each sound file that included the start and end time of each element within a vocalisation.

To quantify vocalisation dissimilarity within pairs, we began by using the function *specan* from the R Package warbleR (Araya-Salas and Smith-Vidaurre, 2017) to extract 28 acoustic parameters; six additional parameters were calculated manually. The final 34 acoustic parameters were transformed (see Supplementary Table 15 for list of acoustic parameters and transformations), centred and scaled, and analysed in a PCA (Figure 1d). Each species had either 2 or 4 sets of associated PC coordinates, one for each sound file. The average Euclidean distance between the vocalisations of each species within a pair was calculated to provide a measure of vocalisation dissimilarity.

#### Patristic Distance

To control for phylogenetic relatedness, we calculated patristic distance (the pairwise distance between the species tips of a phylogenetic tree) using the *cophenetic.phylo* function from the R package ape (Paradis and Schliep, 2019) and a Cornell Lab of Ornithology Open Tree of Life (CLOOTL) tree (McTavish *et al*., 2025; Miller, Sanchez-Reyes and McTavish, 2025). The CLOOTL tree is time-scaled and derived from multiple published molecular phylogenies (see Supplementary Table 16 for tree citations).

#### Syntopy and Sympatry

Using the R package Auk (Strimas-Mackey *et al*., 2023), we extracted eBird occurrence data (Cornell Lab of Ornithology, 2022) for each species. We filtered eBird observations to those from complete checklists, 0 to 400m miles in length, between 1st January 2011 and 31st December 2020, in the United States and Canada, for North American pairs, and Andorra, Austria, Belgium, Czech Republic, Denmark, Estonia, Finland, France, Germany, Ireland, Italy, Latvia, Liechtenstein, Lithuania, Luxembourg, Monaco, Netherlands, Norway, Poland, Portugal, Spain, Sweden, Switzerland, United Kingdom for European analyses. In cases where species pairs occurred on both continents (n = 131), we calculated syntopy and sympatry separately for each continent. Following Drury, Cowen and Grether (2020), these observations were used to calculate measures of range (sympatry) and breeding habitat (syntopy) overlap. For each species, Birds of the World (Billerman *et al*., 2022) species accounts were consulted to determine their breeding months (i.e., the months eggs are laid). For some species, breeding month information was not available from Birds of the World, so a literature search was conducted, and if that failed, breeding months were inferred from closely related species with available data. For each pair, eBird observations were filtered to include only those from the overlapping breeding months. For each pair, we extracted the total number of records matching the aforementioned criteria. Then, for each species in a pair, the number of records that were within 39.4km (the length of a North American BBS route) of a record of the other species was calculated. Mean sympatry (i.e., broad-scale range overlap) for each year was then calculated as the number of records of each species within 39.4km, divided by the total number of records of each species, summed and then divided by 2.

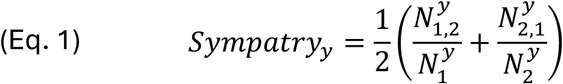

Where N^y^_1_ is the number of records of species 1 in year y, N^y^_2_ is the number of records of species 2 in year y, N^y^_1,2_ is the number of eBird records of species 1 within 39.4km of species 2 within year y and N^y^_2,1_ is the number of eBird records of species 2 within39.4km of species 1 within year y. This value was then averaged across the ten-year period to give an overall measure of range overlap:

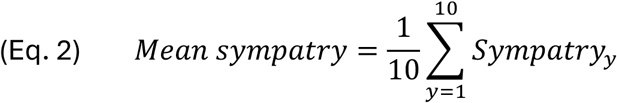

Syntopy (i.e., fine-scale range overlap) was calculated in the same way, but instead of records within 39.4km of each other, they were within 400m (the radius of a point count at a North American BBS stop) of one another (n^y^_1,2_, n^y^_2,1_).

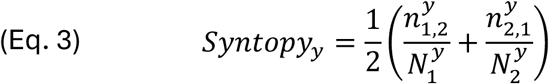

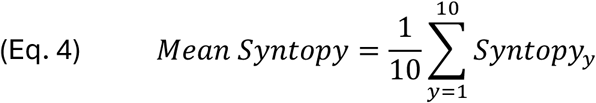

A species pair cannot be syntopic if they are not sympatric. If the breeding months of a species pair do not overlap, or either had no filtered eBird records within the ten-year period, they had no values for sympatry and syntopy and were omitted from analyses. Similarly, in an effort to filter out inaccurate records, there needed to be at least ten records in each year to calculate syntopy. As such, pairs with fewer than ten sympatric records in every year may have had a sympatry value but no value for syntopy and were subsequently removed from analyses. From this filtering, 747 pairs were removed from the analysis.

#### Foraging niche overlap

We used data on foraging niche from AVONICHE (Sayol *et al*., 2026), which classifies the diet and foraging strategy for each species into 32 distinct foraging niche categories. To calculate overlap in foraging niche, for each species pair, we first calculated the minimum percentage across both species for each foraging niche category and then summed the values across the 32 classes of foraging niche. We then divided these summed values by 100 to yield a proportion.

#### Life history traits

Data on migratory strategy and social bond length—factors shown to influence the species-level propensity to hybridise, likely due to their impacts on time available for mate searching (Leighton *et al*., 2021)—were taken from Tobias *et al.,* (2016) for each species. For migratory strategy, species were classified as sedentary, partially migratory and migratory. For social bond length, species were classified as solitary, having short-term pair/group bonds, and long-term pair/group bonds. If either species in a pair were migratory/had long-term pair bonds, the pair were assigned the binary classifier “1” for that specific variable; if neither were they were assigned “0”. 2602 North American pairs contained at least one migratory species, 1874 contained at least one species with long-term social bonds. In Europe, 1585 pairs contained at least one migratory species, 1551 contained at least one species with long-term social bonds.

### Analyses

#### Phylogenetic linear mixed models

All statistical analyses were carried out in R using the package MCMCglmm (Hadfield, 2010). Phylogenetic logistic mixed models (PLMMs) were run with hybridisation as a binary response variable. Pairs containing species without vocalisation data and those without eBird syntopy and sympatry measures were removed from analyses (n=864), leaving 4986 pairs. To facilitate MCMC chain mixing and make model estimates comparable, we transformed and scaled each continuous variable (Supplementary Table 1). Models were run using the full dataset for all pairs, passerines alone and non-passerines alone. Although it would have been interesting to differentiate between song learning passerines (oscines) and other passerines with innate songs (suboscines) (Beecher and Brenowitz, 2005), there were too few suboscine taxa (*n* = 34 species) in our dataset to conduct a robust analysis of this differentiation. We included continent, species identity and a maximum clade credibility CLOOTL phylogeny (Miller, Sanchez-Reyes and McTavish, 2025), specifying nodes representing a pair’s most recent common ancestor as random effects. Uninformative, inverse-Wishart distribution priors were used (Hadfield, 2014), and four chains for each model were run for two million iterations with a burn-in of 20,000 and a thinning interval of 1000, though in a few cases, six million iterations were needed for models to converge. Model convergence was assessed using Gelman-Rubin diagnostics (Gelman and Rubin, 1992) and visual inspection of trace plots. We assessed the potential for multicollinearity by fitting the fixed effect only models (Supplementary Tables 2-4) as generalised linear models and calculating the variance inflation factor (VIF) for each predictor. In all cases, VIF was < 2, indicating low levels of multicollinearity.

We tested whether the relationship between phenotypic similarity and hybrid formation depended on patristic distance by fitting additional models with interaction terms between these variables. We also tested whether the relationship between hybridisation and range, habitat, and niche overlap depended on the patristic distance between taxa. Similarly, we tested for an interactive effect of syntopy and foraging niche overlap. For each data subset, we used backward model selection, eliminating interaction terms with the highest p-values until remaining interaction terms had a pMCMC value < 0.05.

## Acknowledgements

This work was funded by a NERC IAPETUS studentship to DN and NSF/DEB-NERC-2040883 to JPD and GFG. We are grateful to the following PECBMs coordinators for the provision of their data: Norbert Teufelbauer, Antoine Derouaux, Daniel Palm, Benoît Fontaine, Frédéric Jiguet, Lesley Lewis, Mattia Brambilla, Laura Silva, Gianpiero Calvi, Michiel Boom, Bård Stokke, Ingar J. Øien, Tomasz Chodkiewicz, Sergi Herrando, Marc Anton, Åke Lindström, David Noble, Dario Massimino, Sven Trautmann, Johannes Kamp and Ana Gamero. We thank Sally Street and Joe Tobias for helpful comments on a previous draft. We also thank Leonard Davis, Maya Solly and Ted Poulett for their assistance in cleaning and preparing vocalisation data.

