## Supplementary Information for "Spatial overlap and phenotypic similarity predict hybridisation in birds"

Supplementary Tables 1-16
Supplementary Figures 1-6

Supplementary Table 1. Fixed effects included in all models and the corresponding transformations used. All predictor variables were scaled to a mean of 0 and standard deviation of 1.

| Variable | Transformation |
|---|---|
| Morphometric dissimilarity | ln(x+0.4) |
| Colour dissimilarity | ln(x) |
| Pattern dissimilarity | ln(x) |
| Vocalisation dissimilarity | ln(x) |
| Patristic distance | x |
| Syntopy | asin(sqrt(x)) |
| Sympatry | asin(sqrt(x)) |
| Foraging niche overlap | asin(sqrt(x)) |
| Either long-distance migrant | N/A |
| Either have long-term social bonds | N/A |

Supplementary Table 2. Predictors of hybridisation in all pairs (passerines and non-passerines combined) from phylogenetic logistic mixed models (*n* = 4986). The median coefficient estimates from the posterior distribution, as well as 95% credibility intervals and Markov Chain Monte Carlo (MCMC) derived *p*-values are shown. Bold rows indicate fixed effects with 95% credibility intervals that do not overlap 0. pMCMC values are from one chain (results are similar across all chains). The mean phylogenetic signal (λ) for this model is 0.853 (95% CI = 0.673, 0.922). Gelman convergence diagnostics equal 1 or 1.01 for all terms. Significance codes: < 0.05*, < 0.01**, < 0.001***

| Model | Variables | Median | 2.5% | 97.5% | pMCMC | |
|---|---|---|---|---|---|---|
| All | Intercept | -1.495 | -4.519 | 1.784 | 0.346 | |
| | **Morphometric dissimilarity** | **-1.377** | **-1.751** | **-1.043** | **< 5e-04** | *** |
| | **Colour dissimilarity** | **-0.296** | **-0.508** | **-0.088** | **0.004** | ** |
| | **Pattern dissimilarity** | **-0.421** | **-0.652** | **-0.205** | **< 5e-04** | *** |
| | **Vocalisation dissimilarity** | **-0.314** | **-0.551** | **-0.082** | **0.007** | ** |
| | **Patristic distance** | **-4.289** | **-5.275** | **-3.459** | **< 5e-04** | *** |
| | Foraging niche overlap | 0.240 | -0.070 | 0.549 | 0.140 | |
| | **Syntopy** | **0.674** | **0.368** | **0.987** | **< 5e-04** | *** |
| | **Sympatry** | **0.805** | **0.558** | **1.070** | **< 5e-04** | *** |
| | Either long-distance migrant | -0.293 | -0.909 | 0.319 | 0.346 | |
| | Either have long-term social bonds | -0.605 | -1.259 | 0.029 | 0.064 | |

Supplementary Table 3. Predictors of hybridisation in passerine pairs from phylogenetic logistic mixed models ($n$ = 2711). The median coefficient estimates from the posterior distribution, as well as 95% credibility intervals and MCMC derived $p$-values are shown. Bold rows indicate fixed effects with 95% credibility intervals that do not overlap 0. pMCMC values are from one chain (results are similar across all chains). mean phylogenetic signal ($\lambda$) for this model is 0.906 (95% CI = 0.764, 0.961). Gelman convergence diagnostics equal 1 for all terms. Significance codes: < 0.05*, < 0.01**, < 0.001***

| Model | Variables | Median | 2.5% | 97.5% | pMCMC | |
|---|---|---|---|---|---|---|
| Passerines | Intercept | -0.405 | -4.215 | 3.323 | 0.822 | |
| | **Morphometric dissimilarity** | **-0.927** | **-1.312** | **-0.582** | **<5e-04** | *** |
| | Colour dissimilarity | -0.162 | -0.415 | 0.092 | 0.206 | |
| | Pattern dissimilarity | -0.192 | -0.457 | 0.081 | 0.157 | |
| | **Vocalisation dissimilarity** | **-0.499** | **-0.797** | **-0.216** | **<5e-04** | *** |
| | **Patristic distance** | **-1.808** | **-2.348** | **-1.334** | **<5e-04** | *** |
| | Foraging niche overlap | 0.160 | -0.183 | 0.525 | 0.369 | |
| | **Syntopy** | **0.610** | **0.288** | **0.967** | **<5e-04** | *** |
| | **Sympatry** | **0.555** | **0.266** | **0.879** | **<5e-04** | *** |
| | Either long-distance migrant | -0.675 | -1.562 | 0.190 | 0.103 | |
| | Either have long-term social bonds | -0.562 | -1.314 | 0.183 | 0.165 | |

Supplementary Table 4. Predictors of hybridisation in non-passerine pairs from phylogenetic logistic mixed models ($n$ = 2275). The median coefficient estimates from the posterior distribution, as well as 95% credibility intervals and MCMC derived *p*-values are shown. Bold rows indicate fixed effects with 95% credibility intervals that do not overlap 0. pMCMC values are from one chain (results are similar across all chains). The mean phylogenetic signal (λ) for this model was 0.700 (95% CI = 0.081, 0.913). Gelman convergence diagnostics equal 1 or 1.01 for all terms. Significance codes: < 0.05*, < 0.01**, < 0.001***

| Model | Variables | Median | 2.5% | 97.5% | pMCMC | |
|---|---|---|---|---|---|---|
| Non-passerines | Intercept | -2.632 | -6.037 | 1.926 | 0.247 | |
| | **Morphometric dissimilarity** | -1.696 | -2.358 | -1.159 | <3e-04 | *** |
| | **Colour dissimilarity** | -0.467 | -0.866 | -0.101 | 0.019 | * |
| | **Pattern dissimilarity** | -0.789 | -1.202 | -0.416 | <3e-04 | *** |
| | Vocalisation dissimilarity | -0.022 | -0.382 | 0.317 | 0.888 | |
| | **Patristic distance** | -4.380 | -5.915 | -3.20 | <3e-04 | *** |
| | Foraging niche overlap | 0.297 | -0.249 | 0.843 | 0.286 | |
| | Syntopy | 0.567 | -0.024 | 1.179 | 0.057 | |
| | **Sympatry** | 1.408 | 0.949 | 1.961 | <3e-04 | *** |
| | Either long-distance migrant | 0.216 | -0.685 | 1.122 | 0.604 | |
| | Either have long-term social bonds | -1.113 | -2.378 | 0.047 | 0.053 | |

Supplementary Table 5. Results of backward model selection for interaction models. We then eliminated interaction terms that did not predict hybridisation (i.e., terms were removed using backward model selection, one at a time, based on pMCMC values). If no interaction terms were significant ($p < 0.05$), the fixed-effect only model was chosen.

| model terms | DIC mean (range) |
|---|---|
| *Full model* | |
| fixed effects + patristic distance * morphometric dissimilarity + patristic distance * colour dissimilarity + patristic distance * pattern dissimilarity + patristic distance * vocalisation dissimilarity + syntopy * foraging niche overlap + patristic distance*foraging niche overlap + patristic distance * syntopy + patristic distance * sympatry | 1477.94 (1476.38 - 1479.78) |
| fixed effects + patristic distance * morphometric dissimilarity + patristic distance * colour dissimilarity + patristic distance * pattern dissimilarity + syntopy * foraging niche overlap + patristic distance*foraging niche overlap + patristic distance * syntopy + patristic distance * sympatry | 1478.68 (1477.80 - 1479.60) |
| fixed effects + patristic distance * morphometric dissimilarity + patristic distance * colour dissimilarity + patristic distance * pattern dissimilarity + syntopy * foraging niche overlap + patristic distance*foraging niche overlap + patristic distance * sympatry | 1480.55 (1479.04 - 1482.69) |
| fixed effects + patristic distance * morphometric dissimilarity + patristic distance * pattern dissimilarity + syntopy * foraging niche overlap + patristic distance*foraging niche overlap + patristic distance * sympatry | 1481.97 (1480.35 - 1482.76) |
| fixed effects + patristic distance * morphometric dissimilarity + syntopy * foraging niche overlap + patristic distance*foraging niche overlap + patristic distance * sympatry | 1482.04 (1480.02 - 1483.82) |
| fixed effects + syntopy * foraging niche overlap + patristic distance*foraging niche overlap + patristic distance * sympatry | 1483.82 (1481.75 - 1485.66) |
| fixed effects + syntopy * foraging niche overlap + patristic distance * sympatry | 1485.86 (1484.97 - 1486.43) |
| **fixed effects + patristic distance * sympatry** | **1484.78 (1483.19 - 1487.75)** |
| fixed effects (no interactions) | 1474.37 (1472.68 - 1476.46) |
| *Passerines* | |
| fixed effects + patristic distance * morphometric dissimilarity + patristic distance * colour dissimilarity + patristic distance * pattern dissimilarity + patristic distance * vocalisation dissimilarity + syntopy * foraging niche overlap + patristic distance*foraging niche overlap + patristic distance * syntopy + patristic distance * sympatry | 923.77 (922.70 - 924.48) |
| fixed effects + patristic distance * morphometric dissimilarity + patristic distance * pattern dissimilarity + patristic distance * vocalisation dissimilarity + syntopy * foraging niche overlap + patristic distance*foraging niche overlap + patristic distance * syntopy + patristic distance * sympatry | 922.76 (922.38 - 923.30) |
| fixed effects + patristic distance * morphometric dissimilarity + patristic distance * vocalisation dissimilarity + syntopy * foraging niche overlap + patristic distance*foraging niche overlap + patristic distance * syntopy + patristic distance * sympatry | 921.59 (921.123 - 922.36) |
| fixed effects + patristic distance * morphometric dissimilarity + patristic distance * vocalisation dissimilarity + patristic distance*foraging niche overlap + patristic distance * syntopy + patristic distance * sympatry | 920.60 (919.80 - 921.42) |

| | |
|---|---|
| fixed effects + patristic distance * morphometric dissimilarity + patristic distance*foraging niche overlap + patristic distance * syntopy + patristic distance * sympatry | 919.52 (918.93 - 920.50) |
| fixed effects + patristic distance * morphometric dissimilarity + patristic distance * syntopy + patristic distance * sympatry | 918.06 (917.61 - 918.57) |
| fixed effects + patristic distance * morphometric dissimilarity + patristic distance * syntopy | 914.41 (913.80 - 915.77) |
| **fixed effects + patristic distance * syntopy** | **917.11 (916.72 - 917.41)** |
| fixed effects (no interactions) | 911.04 (909.61 - 911.67) |
| *Non-passerines* | |
| fixed effects + patristic distance * morphometric dissimilarity + patristic distance * colour dissimilarity + patristic distance * pattern dissimilarity + patristic distance * vocalisation dissimilarity + syntopy * foraging niche overlap + patristic distance*foraging niche overlap + patristic distance * syntopy + patristic distance * sympatry | 535.10 (534.62 - 535.76) |
| fixed effects + patristic distance * morphometric dissimilarity + patristic distance * colour dissimilarity + patristic distance * vocalisation dissimilarity + syntopy * foraging niche overlap + patristic distance*foraging niche overlap + patristic distance * syntopy + patristic distance * sympatry | 537.69 (535.87 - 539.55) |
| fixed effects + patristic distance * morphometric dissimilarity + patristic distance * colour dissimilarity + patristic distance * vocalisation dissimilarity + syntopy * foraging niche overlap + patristic distance * syntopy + patristic distance * sympatry | 539.73 (537.64 - 541.65) |
| fixed effects + patristic distance * morphometric dissimilarity + patristic distance * colour dissimilarity + patristic distance * vocalisation dissimilarity + syntopy * foraging niche overlap + patristic distance * syntopy | 536.85 (535.29 - 539.02) |
| fixed effects + patristic distance * morphometric dissimilarity + patristic distance * colour dissimilarity + patristic distance * vocalisation dissimilarity + syntopy * foraging niche overlap | 538.83 (537.02 - 542.22) |
| fixed effects + patristic distance * colour dissimilarity + patristic distance * vocalisation dissimilarity + syntopy * foraging niche overlap | 541.92 (540.45 - 543.31) |
| **fixed effects + patristic distance * vocalisation dissimilarity + syntopy * foraging niche overlap** | **545.42 (544.25 - 546.33)** |
| fixed effects (no interactions) | 554.02 (553.01 - 555.66) |

Supplementary Table 6. Predictors of hybridisation in all species pairs from the best-fitting phylogenetic logistic mixed model including interaction terms (*n* = 4986). The median coefficient estimates from the posterior distribution, as well as 95% credibility intervals and MCMC derived *p*-values are shown. Bold rows indicate fixed effects with 95% credibility intervals that do not overlap 0. pMCMC values are from one chain (results are similar across all chains). The mean phylogenetic signal (λ) for this model was 0.831 (95% CI = 0.600, 0.912). Gelman convergence diagnostics equal 1 or 1.01 for all terms. Significance codes: < 0.05*, < 0.01**, < 0.001***

| Model | Variables | Median | 2.5% | 97.5% | pMCMC | |
|---|---|---|---|---|---|---|
| All | Intercept | -1.714 | -2.770 | 1.461 | 0.261 | |
| (+ interactions) | **Morphometric dissimilarity** | **-1.354** | **-1.475** | **-1.028** | **< 5e-04** | *** |
| | **Colour dissimilarity** | **-0.289** | **-0.358** | **-0.088** | **0.005** | ** |
| | **Pattern dissimilarity** | **-0.410** | **-0.485** | **-0.191** | **< 5e-04** | *** |
| | **Vocalisation dissimilarity** | **-0.291** | **-0.375** | **-0.063** | **0.01** | * |
| | Foraging niche overlap | 0.238 | 0.139 | 0.554 | 0.117 | |
| | **Syntopy** | **0.639** | **0.529** | **0.939** | **< 5e-04** | *** |
| | **Sympatry** | **1.166** | **1.041** | **1.538** | **< 5e-04** | *** |
| | **Patristic distance** | **-4.402** | **-4.727** | **-3.576** | **< 5e-04** | *** |
| | Either long-distance migrant | -0.237 | -0.446 | 0.355 | 0.456 | |
| | Either have long-term social bonds | -0.525 | -0.750 | 0.094 | 0.104 | |
| | **Patristic distance*Sympatry** | **0.580** | **0.443** | **0.966** | **0.002** | ** |

Supplementary Table 7. Predictors of hybridisation in passerine pairs from the best-fitting phylogenetic logistic mixed model including interaction terms (*n* = 2711). The median coefficient estimates from the posterior distribution, as well as 95% credibility intervals and MCMC derived *p*-values are shown. Bold rows indicate fixed effects with 95% credibility intervals that do not overlap 0. pMCMC values are from one chain (results are similar across all chains). The mean phylogenetic signal (λ) for this model was 0.893 (95% CI = 0.705, 0.956). Gelman convergence diagnostics equal 1 or 1.01 for all terms. Significance codes: < 0.05*, < 0.01**, < 0.001***

| Model | Variables | Median | 2.5% | 97.5% | pMCMC | |
|---|---|---|---|---|---|---|
| Passerines | Intercept | -0.462 | -1.784 | 3.400 | 0.799 | |
| (+ interactions) | **Morphometric dissimilarity** | **-0.935** | **-1.056** | **-0.585** | **<5e-04** | *** |
| | Colour dissimilarity | -0.154 | -0.240 | 0.094 | 0.194 | |
| | Pattern dissimilarity | -0.193 | -0.286 | 0.063 | 0.165 | |
| | **Vocalisation dissimilarity** | **-0.470** | **-0.567** | **-0.189** | **0.003** | ** |
| | Foraging niche overlap | 0.159 | 0.041 | 0.500 | 0.330 | |
| | **Syntopy** | **0.758** | **0.634** | **1.125** | **<5e-04** | *** |
| | **Patristic distance** | **-1.957** | **-2.139** | **-1.468** | **<5e-04** | *** |
| | **Sympatry** | **0.546** | **0.443** | **0.864** | **0.001** | ** |
| | Either long-distance migrant | -0.688 | -0.979 | 0.170 | 0.117 | |
| | Either have long-term social bonds | -0.526 | -0.785 | 0.227 | 0.176 | |
| | **Patristic distance*Syntopy** | **0.293** | **0.193** | **0.587** | **0.034** | * |

Supplementary Table 8. Predictors of hybridisation in non-passerine pairs from phylogenetic logistic mixed models (*n* = 2275). The median coefficient estimates from the posterior distribution, as well as 95% credibility intervals and MCMC derived *p*-values are shown. Bold rows indicate fixed effects with 95% credibility intervals that do not overlap 0. pMCMC values are from one chain (results are similar across all chains). The mean phylogenetic signal (λ) for this model was 0.679 (95% CI = 0.061, 0.911). Gelman convergence diagnostics equal 1 or 1.01 for all terms. Significance codes: < 0.05*, < 0.01**, < 0.001***

| Model | Variables | Median | 2.5% | 97.5% | pMCMC | |
|---|---|---|---|---|---|---|
| Non-passerines (+ interactions) | Intercept | -2.630 | -6.181 | 1.949 | 0.244 | |
| | **Morphometric dissimilarity** | **-1.675** | **-2.336** | **-1.127** | **<3e-04** | *** |
| | **Colour dissimilarity** | **-0.449** | **-0.855** | **-0.084** | **0.014** | * |
| | **Pattern dissimilarity** | **-0.747** | **-1.157** | **-0.363** | **<3e-04** | *** |
| | Vocalisation dissimilarity | -0.611 | -1.225 | -0.058 | 0.026 | * |
| | **Patristic distance** | **-4.667** | **-6.236** | **-3.443** | **<3e-04** | *** |
| | **Foraging niche overlap** | **0.547** | **-0.048** | **1.163** | **0.075** | |
| | **Syntopy** | **0.908** | **0.243** | **1.603** | **0.003** | ** |
| | **Sympatry** | **1.374** | **0.907** | **1.918** | **<3e-04** | *** |
| | Either long-distance migrant | 0.217 | -0.711 | 1.144 | 0.653 | |
| | **Either have long-term social bonds** | **-1.220** | **-2.505** | **-0.048** | **0.047** | * |
| | **Vocalisation dissimilarity*Patristic distance** | **-0.780** | **-1.38** | **-0.246** | **0.003** | ** |
| | **Foraging niche overlap*Syntopy** | **-0.568** | **-1.106** | **-0.058** | **0.028** | * |

Supplementary Table 9. Predictors of hybridisation in all species pairs in phylogenetic logistic mixed models fit to species pairs sharing a common ancestor (a) ≤ 5 Ma (*n* = 482) and (b) ≤10 Ma (*n* = 2176). The median coefficient estimates from the posterior distribution, as well as 95% credibility intervals and MCMC derived *p*-values are shown. Bold rows indicate fixed effects with 95% credibility intervals that do not overlap 0. pMCMC values are from one chain (results are similar across all chains). Gelman convergence diagnostics equal 1 or 1.01 for all terms. Significance codes: < 0.05*, < 0.01**, < 0.001***

| Model | Variables | Median | 2.5% | 97.5% | pMCMC | |
|---|---|---|---|---|---|---|
| (a) tMRCA ≤ 5Ma | Intercept | 1.386 | -1.263 | 3.934 | 0.266 | |
| | Morphometric dissimilarity | -0.259 | -0.669 | 0.148 | 0.211 | |
| | **Colour dissimilarity** | **-0.398** | **-0.837** | **-0.011** | **0.040** | * |
| | Pattern dissimilarity | -0.058 | -0.486 | 0.361 | 0.727 | |
| Phylogenetic signal (λ) = 0.639 (95% CI = 0.241, 0.879) | **Vocalisation dissimilarity** | **-0.500** | **-0.936** | **-0.094** | **0.011** | * |
| | **Patristic distance** | **-0.934** | **-1.480** | **-0.520** | **<5e-04** | *** |
| | Foraging niche overlap | 0.220 | -0.226 | 0.664 | 0.351 | |
| | **Syntopy** | **0.633** | **0.153** | **1.151** | **0.016** | * |
| | **Sympatry** | **0.548** | **0.114** | **1.120** | **0.014** | * |
| | Either long-distance migrant | -0.001 | -1.118 | 1.120 | 0.995 | |
| | Either have long-term social bonds | -0.323 | -1.525 | 0.802 | 0.560 | |
| (b) tMRCA ≤ 10Ma | Intercept | -0.522 | -3.518 | 2.499 | 0.695 | |
| | **Morphometric dissimilarity** | **-0.759** | **-1.107** | **-0.443** | **<5e-04** | *** |
| | **Colour dissimilarity** | **-0.321** | **-0.603** | **-0.062** | **0.014** | * |
| Phylogenetic signal (λ) = 0.797 (95% CI = 0.522, 0.907) | **Pattern dissimilarity** | **-0.296** | **-0.570** | **-0.032** | **0.024** | * |
| | Vocalisation dissimilarity | -0.260 | -0.545 | 0.016 | 0.071 | |
| | **Patristic distance** | **-1.717** | **-2.183** | **-1.332** | **<5e-04** | *** |
| | Foraging niche overlap | 0.097 | -0.240 | 0.459 | 0.551 | |
| | **Syntopy** | **0.620** | **0.272** | **0.973** | **<5e-04** | *** |
| | **Sympatry** | **0.806** | **0.499** | **1.156** | **<5e-04** | *** |
| | Either long-distance migrant | -0.257 | -1.039 | 0.514 | 0.543 | |
| | Either have long-term social bonds | -0.119 | -0.909 | 0.623 | 0.740 | |

Supplementary Table 10. Hybrid categories from McCarthy (2006) and how these map on to analyses.

| **Acronym** | **Definition** | **Included as hybrid in full analysis** | **Included as frequently hybridising pair (see Supplementary Table 11-12)** |
|---|---|---|---|
| CAENHR | Captive and extensive natural hybridisation reported | x | x |
| ENHI | Extensive natural hybridisation inferred | x | x |
| ENHR | Extensive natural hybridisation reported | x | x |
| CAONHR | Captive and ongoing natural hybridisation reported | x | x |
| ONHR | Ongoing natural hybridisation reported | x | x |
| CANHR | Captive and natural hybridisation reported | x | |
| CHRNHI | Captive hybridisation reported natural hybridisation inferred | x | |
| NHI | Natural hybridisation inferred | x | |
| NHR | Natural hybridisation reported | x | |
| CHR | Captive hybridisation reported | | |

Supplementary Table 11. Predictors of hybridisation in all pairs (passerines and non-passerines combined) from phylogenetic logistic mixed models (*n* = 4986), limiting 'hybrid' classification to only frequently hybridising pairs (*n* = 114) (see Supplementary Table 10). The median coefficient estimates from the posterior distribution, as well as 95% credibility intervals and MCMC derived *p*-values are shown. Bold rows indicate fixed effects with 95% credibility intervals that do not overlap 0. pMCMC values are from one chain (results are similar across all chains). The mean phylogenetic signal (λ) for this model was 0.474 (95% CI = 0.017, 0.848). Gelman convergence diagnostics equal 1 or 1.01 for all terms. Significance codes: < 0.05*, < 0.01**, < 0.001***

| Model | Variables | Median | 2.5% | 97.5% | pMCMC | |
|---|---|---|---|---|---|---|
| Frequent hybridisation (vs. all other species pairs) | Intercept | -3.425 | -7.197 | 2.431 | 0.262 | |
| | **Morphometric dissimilarity** | **-1.236** | **-1.958** | **-0.641** | **<5e-04** | *** |
| | **Colour dissimilarity** | **-0.656** | **-1.083** | **-0.293** | **<5e-04** | *** |
| | Pattern dissimilarity | -0.335 | -0.764 | 0.045 | 0.069 | |
| | Vocalisation dissimilarity | 0.109 | -0.325 | 0.53 | 0.601 | |
| | **Patristic distance** | **-4.211** | **-5.804** | **-3.025** | **<5e-04** | *** |
| | Foraging niche overlap | -0.063 | -0.666 | 0.495 | 0.816 | |
| | **Syntopy** | **0.649** | **0.146** | **1.188** | **0.012** | * |
| | Sympatry | 0.032 | -0.375 | 0.449 | 0.865 | |
| | Either long-distance migrant | -0.426 | -1.435 | 0.559 | 0.372 | |
| | Either have long-term social bonds | -0.293 | -1.439 | 0.725 | 0.549 | |

Supplementary Table 12. Predictors of frequent hybridisation (*n* = 114) from phylogenetic logistic mixed models of hybridising pairs only (*n* = 463) (see Supplementary Table 10). The median coefficient estimates from the posterior distribution, as well as 95% credibility intervals and MCMC derived *p*-values are shown. Bold rows indicate fixed effects with 95% credibility intervals that do not overlap 0. pMCMC values are from one chain (results are similar across all chains). Phylogenetic signal (λ) for this model was 0.238 (95% CI = < 0.001, 0.619). Gelman convergence diagnostics equal 1 or 1.01 for all terms. Significance codes: < 0.05*, < 0.01**, < 0.001***

| Model | Variables | Median | 2.5% | 97.5% | pMCMC | |
|---|---|---|---|---|---|---|
| Frequent hybridisation (vs. other hybrid pairs) | Intercept | -1.452 | -3.431 | 0.901 | 0.178 | |
| | **Morphometric dissimilarity** | **-0.537** | **-1.267** | **-0.034** | **0.030** | * |
| | **Colour dissimilarity** | **-0.609** | **-1.288** | **-0.160** | **0.007** | ** |
| | Pattern dissimilarity | -0.146 | -0.694 | 0.291 | 0.491 | |
| | Vocalisation dissimilarity | 0.235 | -0.305 | 0.760 | 0.355 | |
| | **Patristic distance** | **-1.135** | **-2.336** | **-0.501** | **< 5e-04** | *** |
| | Foraging niche overlap | -0.021 | -0.563 | 0.550 | 0.942 | |
| | Syntopy | 0.413 | -0.110 | 1.011 | 0.110 | |
| | **Sympatry** | **-0.573** | **-1.223** | **-0.049** | **0.028** | * |
| | Either long-distance migrant | -0.601 | -1.919 | 0.471 | 0.274 | |
| | Either have long-term social bonds | -0.075 | -1.321 | 0.964 | 0.892 | |

### Supplementary Table 13. Hybridising pairs and references.

| Species 1 | Species 2 | Ref. | Species 1 | Species 2 | Ref. |
|---|---|---|---|---|---|
| *Acanthis flammea* | *Chloris chloris* | (1) | *Empidonax oberholseri* | *Empidonax wrightii* | (1) |
| *Acanthis flammea* | *Linaria cannabina* | (1) | *Eugenes fulgens* | *Lampornis clemenciae* | (1) |
| *Acanthis flammea* | *Loxia leucoptera* | (3) | *Euphagus cyanocephalus* | *Quiscalus mexicanus* | (1, 2) |
| *Acanthis flammea* | *Spinus pinus* | (1) | *Falco biarmicus* | *Falco peregrinus* | (1) |
| *Acanthis flammea* | *Spinus spinus* | (1) | *Falco cherrug* | *Falco peregrinus* | (1) |
| *Accipiter cooperii* | *Accipiter gentilis* | (4) | *Falco cherrug* | *Falco subbuteo* | (1) |
| *Acrocephalus arundinaceus* | *Acrocephalus scirpaceus* | (1, 6) | *Falco mexicanus* | *Falco peregrinus* | (1, 2) |
| *Acrocephalus dumetorum* | *Acrocephalus palustris* | (1, 7) | *Falco naumanni* | *Falco tinnunculus* | (1) |
| *Acrocephalus palustris* | *Acrocephalus schoenobaenus* | (5) | *Falco peregrinus* | *Falco rusticolus* | (1) |
| *Acrocephalus palustris* | *Acrocephalus scirpaceus* | (1, 8) | *Falco subbuteo* | *Falco vespertinus* | (1) |
| *Acrocephalus schoenobaenus* | *Acrocephalus scirpaceus* | (1) | *Ficedula albicollis* | *Ficedula hypoleuca* | (1, 2, 55) |
| *Aechmophorus clarkii* | *Aechmophorus occidentalis* | (1, 2, 9) | *Fringilla coelebs* | *Fringilla montifringilla* | (1) |
| *Agelaius phoeniceus* | *Quiscalus mexicanus* | (1) | *Fulica americana* | *Gallinula galeata* | (1, 2, 56) |
| *Agelaius phoeniceus* | *Xanthocephalus xanthocephalus* | (1) | *Fulica atra* | *Fulica cristata* | (2) |
| *Aix galericulata* | *Aix sponsa* | (2, 10) | *Fulica atra* | *Gallinula chloropus* | (1, 2, 57) |
| *Aix sponsa* | *Anas platyrhynchos* | (1, 2, 11) | *Galerida cristata* | *Galerida theklae* | (1) |
| *Aix sponsa* | *Aythya collaris* | (1) | *Gallinago gallinago* | *Gallinago media* | (2, 58) |
| *Aix sponsa* | *Bucephala clangula* | (1) | *Geothlypis formosa* | *Geothlypis philadelphia* | (2) |
| *Aix sponsa* | *Lophodytes cucullatus* | (1, 2) | *Geothlypis formosa* | *Setophaga pinus* | (1) |
| *Aix sponsa* | *Mareca americana* | (2) | *Geothlypis philadelphia* | *Geothlypis tolmiei* | (1, 2, 59) |
| *Alectoris graeca* | *Alectoris rufa* | (1, 2) | *Geothlypis philadelphia* | *Geothlypis trichas* | (1, 2) |
| *Alopochen aegyptiaca* | *Anser anser* | (1) | *Geothlypis philadelphia* | *Oporornis agilis* | (1, 2) |
| *Alopochen aegyptiaca* | *Anser cygnoides* | (1) | *Geothlypis tolmiei* | *Geothlypis trichas* | (2) |
| *Alopochen aegyptiaca* | *Cairina moschata* | (1) | *Geothlypis trichas* | *Setophaga petechia* | (2) |
| *Alopochen aegyptiaca* | *Tadorna ferruginea* | (1, 2) | *Haematopus bachmani* | *Haematopus palliatus* | (1, 2, 60) |
| *Alopochen aegyptiaca* | *Tadorna tadorna* | (1) | *Haemorhous cassinii* | *Haemorhous purpureus* | (1) |
| *Ammodramus savannarum* | *Passerculus sandwichensis* | (1, 12) | *Haemorhous mexicanus* | *Haemorhous purpureus* | (1) |
| *Ammospiza caudacuta* | *Ammospiza maritima* | (1) | *Haemorhous purpureus* | *Pinicola enucleator* | (1) |
| *Ammospiza caudacuta* | *Ammospiza nelsoni* | (1, 2, 13) | *Himantopus mexicanus* | *Recurvirostra americana* | (1, 2, 61) |
| *Anas acuta* | *Anas crecca* | (1, 2) | *Hippolais icterina* | *Hippolais polyglotta* | (1, 62) |
| *Anas acuta* | *Anas platyrhynchos* | (1, 2, 14) | *Hirundo rustica* | *Petrochelidon fulva* | (1, 2) |
| *Anas acuta* | *Anas rubripes* | (1, 2) | *Hirundo rustica* | *Petrochelidon pyrrhonota* | (1, 2, 63) |
| *Anas acuta* | *Aythya ferina* | (1) | *Hirundo rustica* | *Riparia riparia* | (1, 2, 64) |
| *Anas acuta* | *Mareca americana* | (1, 2) | *Hirundo rustica* | *Tachycineta bicolor* | (2) |
| *Anas acuta* | *Mareca penelope* | (1, 2) | *Icterus bullockii* | *Icterus galbula* | (1, 2, 65) |
| *Anas acuta* | *Mareca strepera* | (1, 2) | *Icterus graduacauda* | *Icterus gularis* | (1, 2) |
| *Anas acuta* | *Netta rufina* | (1) | *Junco hyemalis* | *Zonotrichia albicollis* | (1, 2, 66) |
| *Anas acuta* | *Somateria mollissima* | (1) | *Lagopus lagopus* | *Lagopus muta* | (67) |
| *Anas acuta* | *Spatula clypeata* | (1) | *Lagopus lagopus* | *Lyrurus tetrix* | (1) |
| *Anas acuta* | *Spatula querquedula* | (1) | *Lagopus lagopus* | *Tetrao urogallus* | (1) |
| *Anas crecca* | *Anas platyrhynchos* | (1, 2) | *Lagopus leucura* | *Tympanuchus phasianellus* | (1) |
| *Anas crecca* | *Aythya ferina* | (1) | *Lagopus muta* | *Lyrurus tetrix* | (1) |
| *Anas crecca* | *Mareca americana* | (2) | *Lanius collurio* | *Lanius minor* | (1) |
| *Anas crecca* | *Mareca penelope* | (1, 2) | *Lanius collurio* | *Lanius senator* | (1, 2, 68) |
| *Anas crecca* | *Mareca strepera* | (1, 2) | *Lanius excubitor* | *Lanius meridionalis* | (1) |
| *Anas crecca* | *Spatula clypeata* | (1) | *Lanius minor* | *Lanius senator* | (1) |
| *Anas crecca* | *Spatula cyanoptera* | (1, 2) | *Leiothlypis celata* | *Leiothlypis ruficapilla* | (2, 69) |
| *Anas crecca* | *Spatula discors* | (1, 2) | *Leiothlypis peregrina* | *Leiothlypis ruficapilla* | (1) |
| *Anas crecca* | *Spatula querquedula* | (1, 2, | *Leiothlypis ruficapilla* | *Setophaga caerulescens* | (1) |
| *Anas diazi* | *Anas fulvigula* | (2) | *Leiothlypis ruficapilla* | *Setophaga magnolia* | (2) |
| *Anas diazi* | *Anas platyrhynchos* | (2) | *Leiothlypis ruficapilla* | *Setophaga ruticilla* | (1) |
| *Anas fulvigula* | *Anas platyrhynchos* | (1, 2, 15) | *Linaria cannabina* | *Linaria flavirostris* | (1) |
| *Anas platyrhynchos* | *Anas rubripes* | (1, 2, 16) | *Linaria cannabina* | *Pyrrhula pyrrhula* | (1) |
| *Anas platyrhynchos* | *Anser cygnoides* | (1) | *Linaria cannabina* | *Serinus serinus* | (1) |
| *Anas platyrhynchos* | *Aythya americana* | (2) | *Linaria cannabina* | *Spinus spinus* | (1) |
| *Anas platyrhynchos* | *Aythya ferina* | (1) | *Lophodytes cucullatus* | *Mergus merganser* | (2) |
| *Anas platyrhynchos* | *Aythya fuligula* | (1, 2) | *Lophodytes cucullatus* | *Mergus serrator* | (1) |
| *Anas platyrhynchos* | *Branta canadensis* | (1) | *Lophophanes cristatus* | *Periparus ater* | (1) |
| *Anas platyrhynchos* | *Branta leucopsis* | (1) | *Lophophanes cristatus* | *Poecile montanus* | (1) |
| *Anas platyrhynchos* | *Bucephala clangula* | (1) | *Lophophanes cristatus* | *Poecile palustris* | (1) |
| *Anas platyrhynchos* | *Cairina moschata* | (1, 2) | *Loxia curvirostra* | *Loxia leucoptera* | (1) |
| *Anas platyrhynchos* | *Mareca americana* | (2) | *Loxia curvirostra* | *Spinus pinus* | (1, 2) |
| *Anas platyrhynchos* | *Mareca penelope* | (2) | *Loxia pytyopsittacus* | *Loxia scotica* | (70) |
| *Anas platyrhynchos* | *Mareca strepera* | (1, 2, 14) | *Luscinia luscinia* | *Luscinia megarhynchos* | (1, 71) |
| *Anas platyrhynchos* | *Netta rufina* | (1, 2, 14) | *Lyrurus tetrix* | *Phasianus colchicus* | (1) |
| *Anas platyrhynchos* | *Somateria mollissima* | (1, 2) | *Lyrurus tetrix* | *Tetrao urogallus* | (1, 2, 72) |
| *Anas platyrhynchos* | *Spatula clypeata* | (2) | *Mareca americana* | *Mareca strepera* | (2) |
| *Anas rubripes* | *Mareca americana* | (2) | *Mareca americana* | *Spatula clypeata* | (2) |
| *Anser albifrons* | *Anser anser* | (1, 17) | *Mareca americana* | *Spatula discors* | (1, 2) |
| *Anser albifrons* | *Anser canagicus* | (2) | *Mareca penelope* | *Mareca strepera* | (1, 2) |
| *Anser albifrons* | *Anser fabalis* | (1, 17) | *Mareca penelope* | *Spatula clypeata* | (1) |
| *Anser albifrons* | *Anser indicus* | (2, 17) | *Mareca penelope* | *Spatula querquedula* | (1) |
| *Anser albifrons* | *Branta canadensis* | (1, 2, 17) | *Mareca strepera* | *Spatula clypeata* | (1, 2, 73) |
| *Anser albifrons* | *Branta hutchinsii* | (2) | *Megascops asio* | *Megascops kennicottii* | (1, 2, 74) |
| *Anser albifrons* | *Branta leucopsis* | (1, 2, 17) | *Megascops kennicottii* | *Megascops trichopsis* | (1, 29) |
| *Anser anser* | *Anser brachyrhynchus* | (1, 17) | *Melanerpes aurifrons* | *Melanerpes carolinus* | (1, 2) |
| *Anser anser* | *Anser cygnoides* | (17) | *Melanerpes aurifrons* | *Melanerpes uropygialis* | (1) |
| *Anser anser* | *Anser fabalis* | (17) | *Meleagris gallopavo* | *Phasianus colchicus* | (1) |
| *Anser anser* | *Anser indicus* | (1, 17) | *Melospiza georgiana* | *Melospiza melodia* | (2) |
| *Anser anser* | *Branta canadensis* | (17) | *Melospiza lincolnii* | *Melospiza melodia* | (2, 75) |
| *Anser anser* | *Branta leucopsis* | (17) | *Melospiza melodia* | *Zonotrichia leucophrys* | (1) |
| *Anser anser* | *Cygnus olor* | (1, 2, 17) | *Melozone aberti* | *Melozone fusca* | (2, 76) |
| *Anser brachyrhynchus* | *Branta leucopsis* | (2) | *Melozone fusca* | *Pipilo maculatus* | (2, 77) |
| *Anser canagicus* | *Branta hutchinsii* | (2) | *Mergus merganser* | *Mergus serrator* | (1, 2) |
| *Anser cygnoides* | *Anser fabalis* | (1) | *Mergus merganser* | *Somateria mollissima* | (2) |
| *Anser cygnoides* | *Anser indicus* | (1) | *Milvus migrans* | *Milvus milvus* | (1, 78) |
| *Anser cygnoides* | *Branta canadensis* | (17) | *Mniotilta varia* | *Setophaga cerulea* | (1) |
| *Anser cygnoides* | *Branta leucopsis* | (1) | *Mniotilta varia* | *Setophaga fusca* | (1) |
| *Anser cygnoides* | *Cygnus olor* | (1) | *Mniotilta varia* | *Setophaga virens* | (1) |
| *Anser fabalis* | *Branta canadensis* | (1, 17) | *Monticola saxatilis* | *Monticola solitarius* | (1) |
| *Anser fabalis* | *Branta leucopsis* | (1) | *Motacilla alba* | *Motacilla cinerea* | (1) |
| *Anser indicus* | *Branta canadensis* | (1, 2, 17) | *Motacilla cinerea* | *Motacilla flava* | (1) |

| | | |
|---|---|---|
| *Anser indicus* | *Branta leucopsis* | (2, 17) |
| *Anthus pratensis* | *Anthus spinoletta* | (1, 18) |
| *Anthus pratensis* | *Anthus trivialis* | (1) |
| *Antigone canadensis* | *Grus americana* | (2) |
| *Aphelocoma californica* | *Cyanocitta stelleri* | (1, 2) |
| *Aphelocoma coerulescens* | *Cyanocitta cristata* | (1) |
| *Aphelocoma woodhouseii* | *Cyanocitta stelleri* | (2) |
| *Apus apus* | *Apus pallidus* | (19) |
| *Aquila chrysaetos* | *Aquila heliaca* | (1, 20) |
| *Aquila chrysaetos* | *Clanga pomarina* | (1) |
| *Archilochus alexandri* | *Archilochus colubris* | (1, 2, 21) |
| *Archilochus alexandri* | *Calothorax lucifer* | (1, 2, 22) |
| *Archilochus alexandri* | *Calypte anna* | (1, 2, 22) |
| *Archilochus alexandri* | *Calypte costae* | (1, 2, 22) |
| *Archilochus alexandri* | *Cynanthus latirostris* | (2, 22) |
| *Archilochus alexandri* | *Lampornis clemenciae* | (1) |
| *Archilochus alexandri* | *Selasphorus calliope* | (1, 2, 22) |
| *Archilochus alexandri* | *Selasphorus platycercus* | (1, 22) |
| *Archilochus alexandri* | *Selasphorus rufus* | (2, 22) |
| *Archilochus alexandri* | *Selasphorus sasin* | (1, 22) |
| *Ardea alba* | *Ardea cinerea* | (1, 2, 23) |
| *Ardea alba* | *Ardea herodias* | (1, 2, 24) |
| *Ardea cinerea* | *Ardea purpurea* | (1, 25) |
| *Ardea cinerea* | *Egretta garzetta* | (1) |
| *Ardeola ralloides* | *Bubulcus ibis* | (26) |
| *Asio flammeus* | *Asio otus* | (1) |
| *Aythya affinis* | *Aythya americana* | (2) |
| *Aythya affinis* | *Aythya collaris* | (1, 2) |
| *Aythya affinis* | *Aythya valisineria* | (1) |
| *Aythya americana* | *Aythya collaris* | (1, 2) |
| *Aythya americana* | *Aythya valisineria* | (1, 2) |
| *Aythya collaris* | *Aythya valisineria* | (1, 2) |
| *Aythya collaris* | *Mergus serrator* | (2) |
| *Aythya ferina* | *Aythya fuligula* | (1, 2, 14) |
| *Aythya ferina* | *Aythya marila* | (1) |
| *Aythya ferina* | *Aythya nyroca* | (1, 2, 14) |
| *Aythya ferina* | *Netta rufina* | (1, 2, 14) |
| *Aythya fuligula* | *Aythya marila* | (1, 2, 14) |
| *Aythya fuligula* | *Aythya nyroca* | (1, 2, 14) |
| *Aythya fuligula* | *Netta rufina* | (1, 2, 14) |
| *Aythya fuligula* | *Spatula querquedula* | (1) |
| *Aythya marila* | *Bucephala clangula* | (1) |
| *Aythya nyroca* | *Netta rufina* | (1, 2) |
| *Baeolophus atricristatus* | *Baeolophus bicolor* | (1, 2, 27) |
| *Baeolophus bicolor* | *Poecile atricapillus* | (2) |
| *Baeolophus bicolor* | *Poecile carolinensis* | (2) |
| *Bonasa umbellus* | *Canachites canadensis* | (1) |
| *Bonasa umbellus* | *Dendragapus obscurus* | (1) |
| *Branta canadensis* | *Branta hutchinsii* | (28) |
| *Branta canadensis* | *Branta leucopsis* | (1, 2, 17) |
| *Branta canadensis* | *Cygnus olor* | (1) |
| *Branta hutchinsii* | *Branta leucopsis* | (2, 17) |
| *Brotogeris chiriri* | *Brotogeris versicolurus* | (2) |
| *Bubo virginianus* | *Strix nebulosa* | (29) |
| *Bubulcus ibis* | *Egretta caerulea* | (1, 30) |
| *Bubulcus ibis* | *Egretta garzetta* | (31) |
| *Bubulcus ibis* | *Egretta thula* | (1) |
| *Bucephala albeola* | *Bucephala clangula* | (2, 32) |
| *Bucephala albeola* | *Lophodytes cucullatus* | (1, 2, 32) |
| *Bucephala clangula* | *Bucephala islandica* | (1, 2, 33) |
| *Bucephala clangula* | *Lophodytes cucullatus* | (1, 2) |
| *Bucephala clangula* | *Mergellus albellus* | (1, 2) |
| *Bucephala clangula* | *Mergus merganser* | (1) |
| *Bucephala clangula* | *Mergus serrator* | (1) |
| *Bucephala islandica* | *Lophodytes cucullatus* | (2, 34) |
| *Buteo buteo* | *Milvus migrans* | (1) |
| *Buteo jamaicensis* | *Buteo lineatus* | (2) |
| *Buteo jamaicensis* | *Buteo regalis* | (2, 35) |
| *Buteo jamaicensis* | *Buteo swainsoni* | (36) |
| *Buteo lineatus* | *Buteo plagiatus* | (1) |
| *Buteo lineatus* | *Buteo platypterus* | (2) |
| *Cairina moschata* | *Tadorna tadorna* | (1) |
| *Calcarius lapponicus* | *Plectrophenax nivalis* | (2, 37) |
| *Calidris alpina* | *Calidris maritima* | (1, 2) |
| *Calidris minuta* | *Calidris temminckii* | (1) |
| *Callipepla californica* | *Callipepla gambelii* | (1, 2, 38) |
| *Callipepla californica* | *Callipepla squamata* | (1, 2) |
| *Callipepla californica* | *Colinus virginianus* | (1) |
| *Callipepla californica* | *Oreortyx pictus* | (1) |
| *Callipepla gambelii* | *Callipepla squamata* | (1, 2) |
| *Callipepla squamata* | *Colinus virginianus* | (1, 2) |
| *Calothorax lucifer* | *Calypte anna* | (2) |
| *Calothorax lucifer* | *Calypte costae* | (2) |
| *Calypte anna* | *Calypte costae* | (1, 2, 22) |
| *Calypte anna* | *Cynanthus latirostris* | (2) |
| *Calypte anna* | *Lampornis clemenciae* | (1, 21) |
| *Calypte anna* | *Selasphorus calliope* | (1, 2, 22) |
| *Calypte anna* | *Selasphorus rufus* | (1, 2, 22) |
| *Calypte anna* | *Selasphorus sasin* | (1, 2, 22) |
| *Calypte costae* | *Lampornis clemenciae* | (1) |
| *Calypte costae* | *Selasphorus calliope* | (1, 22) |
| *Calypte costae* | *Selasphorus platycercus* | (1, 21) |
| *Calypte costae* | *Selasphorus sasin* | (22) |
| *Canachites canadensis* | *Dendragapus obscurus* | (1) |
| *Cardellina canadensis* | *Geothlypis philadelphia* | (1) |
| *Cardinalis cardinalis* | *Cardinalis sinuatus* | (1, 2) |
| *Carduelis carduelis* | *Chloris chloris* | (1) |
| *Nyctanassa violacea* | *Nycticorax nycticorax* | (2, 79) |
| *Oxyura jamaicensis* | *Oxyura leucocephala* | (1, 2, 80) |
| *Parus major* | *Poecile palustris* | (1) |
| *Passer domesticus* | *Passer hispaniolensis* | (1, 2, 81) |
| *Passer domesticus* | *Passer italiae* | (81) |
| *Passer domesticus* | *Passer montanus* | (1, 2, 82) |
| *Passer hispaniolensis* | *Passer italiae* | (81) |
| *Passer hispaniolensis* | *Passer montanus* | (1, 2) |
| *Passer italiae* | *Passer montanus* | (1) |
| *Passerina amoena* | *Passerina cyanea* | (1, 2, 83) |
| *Passerina caerulea* | *Passerina cyanea* | (2) |
| *Passerina ciris* | *Passerina cyanea* | (1, 2) |
| *Passerina ciris* | *Passerina versicolor* | (1, 2) |
| *Periparus ater* | *Poecile montanus* | (1) |
| *Petrochelidon fulva* | *Petrochelidon pyrrhonota* | (2) |
| *Petrochelidon pyrrhonota* | *Tachycineta bicolor* | (1) |
| *Petrochelidon pyrrhonota* | *Tachycineta thalassina* | (1, 2) |
| *Phasianus colchicus* | *Tetrao urogallus* | (1) |
| *Pheucticus ludovicianus* | *Pheucticus melanocephalus* | (2, 84) |
| *Phoenicurus ochruros* | *Phoenicurus phoenicurus* | (1, 2, 85) |
| *Phoenicurus phoenicurus* | *Saxicola rubetra* | (2, 86) |
| *Phylloscopus bonelli* | *Phylloscopus sibilatrix* | (1, 87) |
| *Phylloscopus collybita* | *Phylloscopus ibericus* | (88) |
| *Phylloscopus collybita* | *Phylloscopus trochilus* | (1) |
| *Phylloscopus trochiloides* | *Phylloscopus trochilus* | (1) |
| *Picus canus* | *Picus viridis* | (2, 89) |
| *Pipilo chlorurus* | *Pipilo maculatus* | (1, 2) |
| *Pipilo erythrophthalmus* | *Pipilo maculatus* | (1, 2) |
| *Piranga ludoviciana* | *Piranga olivacea* | (1, 2, 90) |
| *Piranga ludoviciana* | *Piranga rubra* | (1, 2, 91) |
| *Piranga olivacea* | *Piranga rubra* | (1, 92) |
| *Plegadis chihi* | *Plegadis falcinellus* | (1, 2, 93) |
| *Podiceps auritus* | *Podiceps nigricollis* | (1) |
| *Podiceps cristatus* | *Podiceps grisegena* | (1) |
| *Poecile atricapillus* | *Poecile carolinensis* | (1, 2, 94) |
| *Poecile atricapillus* | *Poecile gambeli* | (1, 2, 95) |
| *Poecile atricapillus* | *Poecile hudsonicus* | (2, 96) |
| *Poecile atricapillus* | *Poecile rufescens* | (97) |
| *Poecile cinctus* | *Poecile montanus* | (1, 2) |
| *Poecile gambeli* | *Poecile hudsonicus* | (2, 97) |
| *Poecile hudsonicus* | *Poecile rufescens* | (97) |
| *Polioptila californica* | *Polioptila melanura* | (1) |
| *Pooecetes gramineus* | *Spizella pusilla* | (1) |
| *Protonotaria citrea* | *Setophaga petechia* | (1) |
| *Psittacula eupatria* | *Psittacula krameri* | (98) |
| *Pyrrhocorax graculus* | *Pyrrhocorax pyrrhocorax* | (1) |
| *Quiscalus major* | *Quiscalus mexicanus* | (1, 2) |
| *Quiscalus mexicanus* | *Quiscalus quiscula* | (2, 99) |
| *Rallus crepitans* | *Rallus elegans* | (1, 2, 100) |
| *Saxicola rubetra* | *Saxicola rubicola* | (1) |
| *Sayornis nigricans* | *Sayornis phoebe* | (2, 101) |
| *Sayornis nigricans* | *Sayornis saya* | (2) |
| *Selasphorus calliope* | *Selasphorus platycercus* | (2, 22) |
| *Selasphorus calliope* | *Selasphorus rufus* | (1, 2, 22) |
| *Selasphorus platycercus* | *Selasphorus rufus* | (2, 22) |
| *Selasphorus rufus* | *Selasphorus sasin* | (1, 2, 22) |
| *Setophaga americana* | *Setophaga cerulea* | (1, 2, 102) |
| *Setophaga americana* | *Setophaga coronata* | (1) |
| *Setophaga americana* | *Setophaga dominica* | (1, 2, 103) |
| *Setophaga americana* | *Setophaga ruticilla* | (1, 2) |
| *Setophaga caerulescens* | *Setophaga cerulea* | (2, 104) |
| *Setophaga caerulescens* | *Setophaga magnolia* | (2) |
| *Setophaga caerulescens* | *Setophaga pensylvanica* | (2) |
| *Setophaga caerulescens* | *Setophaga petechia* | (105) |
| *Setophaga castanea* | *Setophaga coronata* | (1) |
| *Setophaga castanea* | *Setophaga fusca* | (1, 2) |
| *Setophaga castanea* | *Setophaga striata* | (1, 2) |
| *Setophaga cerulea* | *Vermivora cyanoptera* | (2, 106) |
| *Setophaga citrina* | *Setophaga petechia* | (2) |
| *Setophaga coronata* | *Setophaga dominica* | (2) |
| *Setophaga coronata* | *Setophaga graciae* | (1, 2) |
| *Setophaga coronata* | *Setophaga magnolia* | (1, 2) |
| *Setophaga coronata* | *Setophaga nigrescens* | (2) |
| *Setophaga coronata* | *Setophaga palmarum* | (1, 2) |
| *Setophaga coronata* | *Setophaga pinus* | (1) |
| *Setophaga coronata* | *Setophaga tigrina* | (1, 2) |
| *Setophaga coronata* | *Setophaga townsendi* | (1, 2, 107) |
| *Setophaga coronata* | *Setophaga virens* | (2) |
| *Setophaga discolor* | *Setophaga petechia* | (2) |
| *Setophaga discolor* | *Vermivora cyanoptera* | (2) |
| *Setophaga fusca* | *Setophaga kirtlandii* | (108) |
| *Setophaga graciae* | *Setophaga nigrescens* | (1, 2) |
| *Setophaga magnolia* | *Setophaga palmarum* | (2) |
| *Setophaga magnolia* | *Setophaga pensylvanica* | (2) |
| *Setophaga magnolia* | *Setophaga petechia* | (109) |
| *Setophaga magnolia* | *Setophaga ruticilla* | (110) |
| *Setophaga nigrescens* | *Setophaga occidentalis* | (1, 2, 111) |
| *Setophaga nigrescens* | *Setophaga townsendi* | (1, 2) |
| *Setophaga occidentalis* | *Setophaga townsendi* | (1, 2, 111) |
| *Setophaga petechia* | *Setophaga ruticilla* | (2) |
| *Setophaga pinus* | *Vermivora chrysoptera* | (1) |
| *Setophaga tigrina* | *Setophaga townsendi* | (1) |
| *Sialia currucoides* | *Sialia mexicana* | (1, 2, 112) |
| *Sialia currucoides* | *Sialia sialis* | (1, 2) |
| *Sialia mexicana* | *Sialia sialis* | (2) |
| *Somateria mollissima* | *Tadorna tadorna* | (1) |
| *Spatula clypeata* | *Spatula cyanoptera* | (1, 2) |

| | | | | | |
|---|---|---|---|---|---|
| *Carduelis carduelis* | *Linaria cannabina* | (1) | *Spatula clypeata* | *Spatula discors* | (1, 2, 113) |
| *Carduelis carduelis* | *Pyrrhula pyrrhula* | (1) | *Spatula clypeata* | *Spatula querquedula* | (1, 2) |
| *Carduelis citrinella* | *Chloris chloris* | (1) | *Spatula cyanoptera* | *Spatula discors* | (1, 2, 114) |
| *Catharus bicknelli* | *Catharus fuscescens* | (2, 39) | *Sphyrapicus nuchalis* | *Sphyrapicus ruber* | (1, 2, 115) |
| *Catharus fuscescens* | *Catharus ustulatus* | (1) | *Sphyrapicus nuchalis* | *Sphyrapicus thyroideus* | (1) |
| *Cecropis daurica* | *Hirundo rustica* | (2) | *Sphyrapicus nuchalis* | *Sphyrapicus varius* | (1, 2, 115) |
| *Centrocercus urophasianus* | *Dendragapus obscurus* | (1) | *Sphyrapicus ruber* | *Sphyrapicus varius* | (1, 2, 115) |
| *Centrocercus urophasianus* | *Tympanuchus phasianellus* | (1, 2, 40) | *Spinus pinus* | *Spinus tristis* | (2) |
| *Certhia brachydactyla* | *Certhia familiaris* | (1) | *Spinus psaltria* | *Spinus tristis* | (2) |
| *Chloris chloris* | *Linaria cannabina* | (1) | *Spizella atrogularis* | *Spizella breweri* | (2, 116) |
| *Chrysolophus amherstiae* | *Chrysolophus pictus* | (1, 41) | *Spizella breweri* | *Spizella pallida* | (2) |
| *Circus cyaneus* | *Circus macrourus* | (1, 2, 42) | *Spizella pallida* | *Spizella passerina* | (1, 2, 117) |
| *Coccyzus americanus* | *Coccyzus erythropthalmus* | (1) | *Spizella pallida* | *Spizella pusilla* | (1, 2, 118) |
| *Colaptes auratus* | *Colaptes chrysoides* | (1, 2, 43) | *Spizella passerina* | *Spizella pusilla* | (2) |
| *Columba livia* | *Streptopelia decaocto* | (1) | *Stelgidopteryx serripennis* | *Tachycineta thalassina* | (2) |
| *Columba livia* | *Zenaida macroura* | (44) | *Streptopelia decaocto* | *Streptopelia turtur* | (1, 2, 119) |
| *Contopus pertinax* | *Contopus sordidulus* | (1) | *Streptopelia decaocto* | *Zenaida macroura* | (2, 120) |
| *Contopus sordidulus* | *Contopus virens* | (1, 2, 45) | *Strix nebulosa* | *Strix varia* | (29) |
| *Contopus sordidulus* | *Empidonax traillii* | (1) | *Strix occidentalis* | *Strix varia* | (1, 2, 29) |
| *Corthylio calendula* | *Regulus satrapa* | (1) | *Sturnella magna* | *Sturnella neglecta* | (1, 2, 121) |
| *Corvus corax* | *Corvus corone* | (1) | *Sturnus unicolor* | *Sturnus vulgaris* | (1) |
| *Corvus corax* | *Corvus cryptoleucus* | (1) | *Sylvia atricapilla* | *Curruca communis* | (1) |
| *Cyanocitta cristata* | *Cyanocitta stelleri* | (1, 2) | *Sylvia borin* | *Curruca nisoria* | (1) |
| *Cyanocitta cristata* | *Cyanocorax yncas* | (2) | *Tachycineta bicolor* | *Tachycineta thalassina* | (2) |
| *Cygnus atratus* | *Cygnus olor* | (17) | *Tadorna ferruginea* | *Tadorna tadorna* | (1, 2) |
| *Cygnus buccinator* | *Cygnus olor* | (2) | *Tetrastes bonasia* | *Lagopus lagopus* | (1) |
| *Cygnus cygnus* | *Cygnus olor* | (1, 17) | *Tetrastes bonasia* | *Lagopus muta* | (1) |
| *Cynanthus latirostris* | *Eugenes fulgens* | (1) | *Tetrastes bonasia* | *Lyrurus tetrix* | (1) |
| *Cynanthus latirostris* | *Leucolia violiceps* | (2, 21) | *Toxostoma crissale* | *Toxostoma redivivum* | (2) |
| *Delichon urbicum* | *Hirundo rustica* | (1, 2, 46) | *Toxostoma curvirostre* | *Toxostoma longirostre* | (2) |
| *Delichon urbicum* | *Riparia riparia* | (1, 47) | *Toxostoma curvirostre* | *Toxostoma rufum* | (1) |
| *Dendragapus obscurus* | *Phasianus colchicus* | (1) | *Toxostoma lecontei* | *Toxostoma redivivum* | (1) |
| *Dendragapus obscurus* | *Tympanuchus phasianellus* | (1, 2) | *Tringa erythropus* | *Tringa totanus* | (1) |
| *Dendrocopos leucotos* | *Dendrocopos major* | (1) | *Turdus iliacus* | *Turdus merula* | (1) |
| *Dendrocopos major* | *Dendrocopos syriacus* | (1, 2, 48) | *Turdus iliacus* | *Turdus philomelos* | (1) |
| *Dendrocopos major* | *Dendrocoptes medius* | (1) | *Turdus iliacus* | *Turdus pilaris* | (1) |
| *Dendrocygna autumnalis* | *Dendrocygna bicolor* | (2, 49) | *Turdus merula* | *Turdus philomelos* | (1) |
| *Dryobates borealis* | *Dryobates villosus* | (1) | *Turdus merula* | *Turdus pilaris* | (1) |
| *Dryobates nuttallii* | *Dryobates pubescens* | (1, 2) | *Turdus merula* | *Turdus torquatus* | (1) |
| *Dryobates nuttallii* | *Dryobates scalaris* | (1, 2, 50) | *Turdus pilaris* | *Turdus viscivorus* | (1) |
| *Dryobates nuttallii* | *Dryobates villosus* | (2) | *Tympanuchus cupido* | *Tympanuchus pallidicinctus* | (1, 2, 122) |
| *Dryobates pubescens* | *Dryobates scalaris* | (1, 2) | *Tympanuchus cupido* | *Tympanuchus phasianellus* | (1, 2, 123) |
| *Dryobates pubescens* | *Dryobates villosus* | (2, 51) | *Tyrannus couchii* | *Tyrannus forficatus* | (1, 2, 124) |
| *Dryobates scalaris* | *Dryobates villosus* | (1, 2) | *Tyrannus couchii* | *Tyrannus melancholicus* | (1) |
| *Egretta caerulea* | *Egretta thula* | (1, 2) | *Tyrannus dominicensis* | *Tyrannus melancholicus* | (2, 125) |
| *Egretta caerulea* | *Egretta tricolor* | (2) | *Tyrannus forficatus* | *Tyrannus verticalis* | (1, 2, 126) |
| *Egretta rufescens* | *Egretta tricolor* | (2) | *Tyrannus tyrannus* | *Tyrannus verticalis* | (1, 2, 127) |
| *Egretta thula* | *Egretta tricolor* | (1, 2) | *Vermivora chrysoptera* | *Vermivora cyanoptera* | (2, 128) |
| *Egretta tricolor* | *Nycticorax nycticorax* | (1) | *Vireo bellii* | *Vireo griseus* | (129) |
| *Emberiza cirlus* | *Emberiza citrinella* | (1) | *Vireo cassinii* | *Vireo plumbeus* | (1) |
| *Emberiza citrinella* | *Emberiza schoeniclus* | (1) | *Vireo cassinii* | *Vireo solitarius* | (1) |
| *Empidonax alnorum* | *Empidonax traillii* | (1, 52) | *Vireo flavifrons* | *Vireo solitarius* | (1, 2) |
| *Empidonax difficilis* | *Empidonax occidentalis* | (1, 2, 53) | *Vireo gilvus* | *Vireo olivaceus* | (2, 130) |
| *Empidonax hammondii* | *Empidonax oberholseri* | (1) | *Vireo olivaceus* | *Vireo philadelphicus* | (2, 131) |
| *Empidonax hammondii* | *Empidonax wrightii* | (1) | *Zenaida asiatica* | *Zenaida macroura* | (132) |
| *Empidonax minimus* | *Empidonax virescens* | (54) | *Zonotrichia albicollis* | *Zonotrichia leucophrys* | (1, 2) |

1. McCarthy, E.M. (2006) Handbook of Avian Hybrids of the World. Oxford, UK: Oxford University Press.
2. Clements, J.F. et al. (2021) 'The eBird/Clements checklist of Birds of the World: v2021.' Available at: https://www.birds.cornell.edu/clementschecklist/introduction/updateindex/august-2021/2021-citation-checklist-download/.
3. Red'kin, Y.A. and Sotnikov, V.N. 2021. On hybrids between White-winged Crossbill *Loxia leucoptera* and Common Redpoll *Acanthis flammea* in nature. Russkii Ornitologicheskii Zhurnal Ekspress Vypusk. 30(2026): 287-299.
4. Haughey C.L., Neslosn A., Napier P., Rosenfield R.N., Sonsthagen S.A. & Talbot S.L. 2019.Genetic confirmation of a natural hybrid between a Northern Goshawk (*Accipiter gentilis*) and a Cooper's Hawk (*A. cooperii*). The Wilson Journal of Ornithology. 131(4): 838–844.
5. Lifjeld, J.T., Marthinsen, G., Myklebust, M. et al. A wild Marsh Warbler × Sedge Warbler hybrid (*Acrocephalus palustris* × *A. schoenobaenus*) in Norway documented with molecular markers. J Ornithol 151, 513–517 (2010). https://doi.org/10.1007/s10336-009-0469-0
6. Ion, C., Bolboaca, L., Ciorpac, M. et al. A Great Reed Warbler × Reed Warbler hybrid (*Acrocephalus arundinaceus* × *Acrocephalus scirpaceus*) in northeastern Romania. Journal of Ornithology. 153: 975–978. https://doi.org/10.1007/s10336-012-0835-1
7. Lindholm A., Bensch S., Dowsett-Lemaire F., Forsten A. & Kärkkäinen H. 2007. Hybrid Marsh x Blyth's Reed Warbler with mixed song in Finland in June 2003. Dutch Birding. 29: 223-231.
8. Reifová, R., Majerová, V., Reif, J. et al. Patterns of gene flow and selection across multiple species of Acrocephalus warblers: footprints of parallel selection on the Z chromosome. BMC Evol Biol 16, 130 (2016). https://doi.org/10.1186/s12862-016-0692-2

9. Konter A. 2011. Interbreeding of *Aechmophorus* grebes.The Wilson Journal of Ornithology. 123(1): 132–136.

10. Dubois, P.J. (2007). Les espèces d'oiseaux allochtones en France. LPO, Paris.
11. Rohwer S, Wood CS, Peters JL, Miller ET, Cagley D, Butcher BG, et al. (2022) Interspecific forced copulations generate most hybrids in broadly sympatric ducks. PLoS ONE 17(9): e0274059. https://doi.org/10.1371/journal.pone.0274059
12. Jones A.L., Shriver G.W., Bulgin N.L., Lockwood R. & Vickery P.D. 2003. A probable grasshopper x savannah sparrow hybrid singing a song sparrow song. The Wilson Bulletin. 115(3): 231-236.
13. Walsh J., Shriver G.W., Olsen B.J., O'Brien K.M. & Kovach A.I. 2015. Relationship of phenotypic variation and genetic admixture in the Saltmarsh–Nelson's sparrow hybrid zone. The Auk. 132: 704–716.

36. Hull J.M., Savage W., Smith J.P., Murphy N., Cullen L., Hutchins A.C & Ernest H.B. 2007. Hybridisation among Buteos: Swainson's hawks (*Buteo swainsoni*) x red-tailed hawks (*Buteo jamaicensis*). The Wilson Journal of Ornithology. 119(4): 579–584.
37. Macdonald, C.A., Martin, T., Ludkin, R., Hussell, D.J., Lamble, D. and Love, O.P., 2012. First report of a snow bunting× Lapland longspur hybrid. Arctic, pp.344-348.
38. Gee J.M. 2003. How a hybrid zone is maintained: Behavioural mechanisms of interbreeding, between California and Gambel's quails (*Callipepla californica* and *C.gambelii*. Evolution. 57(10): 2407–2415.
39. Martinsen E.S., McFarland K.P., Rimmer C.C. 2018. Documentation of a hybrid Bicknell's Thrush (*Catharus bicknelli*) Veery (*C. fuscescens*) using vocalization and genetic data. The Wilson Journal of Ornithology. 130(1): 70–80.
40. Aldridge C.L., Oyler-McCance S.J. & Brigham R.M. 2001. Occurrence of greater sage-grouse x sharp-taield grouse hybrids in Alberta. The Condor. 103: 657–660.
41. Li, Z., Zhou, J., Gao, M.,Liang, W., Dong, L.. 2022. Parapatric speciation with recurrent gene flow of two sexual dichromatic pheasants. Avian Research: 13. https://doi.org/10.1016/j.avrs.2022.100031.
42. Forsman, D. and Erterius, D., 2012. Pallid Harriers in northwest Europe and the identification of presumed Pallid Harrier x Hen Harrier hybrids. Birding World, 25(2), pp.68-75.
43. Manthey J.D., Geiger M. & Moyle R.G. 2016: Relationships of morphological groups in the northern flicker superspecies complex (*Colaptes auratus & C. chrysoide*s). Systematics and Biodiversity. 15(3): 183-191.
44. Hernández F., Brown J.L., Kaminski M., Harvey M.G. & Lavretsky P. 2021. Genomic evidence for rare hybridization and large demographic changes in the evolutionary histories of four north american dove species. Animals. 11: 2677.
45. Manthey J.D. & Robbins M.B. 2016. Genomic insights into hybridization in a localized region of sympatry between pewee sister species (*Contopus sordidulus* × *C. virens*) and their chromosomal patterns of differentiation. Avian Research. 7: 6.
46. Pickles, A., 2014. Hybrid: Barn Swallow x Common House Martin. Biodiversity Observations, pp.336-337.
47. Dunning, J., Hanmer, H. and Christmas, S.E., 2014. Hybridisation between House Martin *Delichon urbicum* and Sand Martin *Riparia riparia*: a new observation and review of past occurrences as a case study into hybrid reporting rates. Ringing & Migration, 29(2), pp.86-89.
48. Michalczuk, J., McDevitt, A.D., Mazgajski, T.D., Figarski, T., Ilieva, M., Bujoczek, M., Malczyk, P. and Kajtoch, Ł., 2014. Tests of multiple molecular markers for the identification of Great Spotted and Syrian Woodpeckers and their hybrids. Journal of Ornithology, 155, pp.591-600.
49. Jones D. 2016. Fulvous x Black-bellied whistling duck hybrid… A photographic record. Bulletin of the Texas Ornithological Society. 50:70-71.
50. Manthey J.D., Boissinot S. & Moyle R.J. 2019. Biodiversity genomics of North American Dryobates woodpeckers reveals little gene flow across the *D. nuttallii x D. scalaris* contact zone. The Auk. 236(2).
51. Mlodinow S.G., Leukering T., Brooks T. & Moore N. 2015. Apparent hybrid downy woodpecker x hairy woodpecker in Colorado. Western Birds. 46(1): 2-7.
52. Bemmels J.B., Bramwell A.C., Luzuriaga-Aveiga V.E., Anderson S.A.S., Mikkelsen E.K. & Weir J.T. 2021. Geographic contact drives increased reproductive isolation in two cryptic *Empidonax* flycatchers. Molecular Ecology. 30: 4833–4844.
53. Rush A.C., Cannings R.J. & Irwin D.E. 2009. Analysis of multilocus DNA reveals hybridisation in a contact zone between *Empidonax* flycatchers. Journal of Avian Biology. 40: 614-624.
54. Murray M. & Donald K. 2005. A probable acadian flycatcher x least flycatcher hybrid. Passenger Pigeon. 67(3): 337-343.
55. Vallin, N., Rice, A.M., Arntsen, H., Kulma, K. and Qvarnström, A., 2012. Combined effects of interspecific competition and hybridization impede local coexistence of *Ficedula* flycatchers. Evolutionary Ecology, 26, pp.927-942.
56. Erickson R.A. 2020. Two photographed American coot x common gallinule hybrids. Western Birds. 51: 150–156.
57. Rodrígueza, B., Curbelob, J. and Carrascoa, N., Hybrid Eurasian Coot *Fulica atra*× Common Moorhen *Gallinula chloropus* on Tenerife, Canary Islands. Bulletin of African Bird Club. 17(2): 207-209.
58. Höglund, J., Sæther, S.A., Fiske, P., Wheatcroft, D. and Kålås, J.A., 2015. A hybrid snipe *Gallinago gallinago*× *G. media* found in the wild. Journal of Ornithology, 156, pp.819-827.

59. Irwin D.E., Brelsford A., Toews D.P.L., MacDonald C. & Phinney M. 2009. Extensive hybridization in a contact zone between MacGillivray's warblers *Oporornis tolmiei* and mourning warblers *O. philadelphia* detected using molecular and morphological analyses. Journal of Avian Biology. 40: 539-552.
60. Carmona C., Ayala-Perez V., Hernandez-Alvarez A., Mendosa L.F., Marron G., Arce N., Vidal S., Danemann G.D. 2020. Breeding populations of shorebirds in northwestern Mexico wetlands. Huitzil. 21(2): e-581.
61. Morlan J., Dakin R.E. & Rosso J. 2004. Apparent hybrids between the American Avocet and Black-necked Stilt in California. Western Birds. 35(1): 57-59.
62. Secondi, J., Faivre, B. and Bensch, S., 2006. Spreading introgression in the wake of a moving contact zone. Molecular Ecology, 15(9), pp.2463-2475.
63. Brown C.R. & Page C.E. 2015. A hybrid cliff × barn swallow from Western Nebraska. Nebraska Bird Review: 83(1): 31-35.
64. Todte I., Bechert K. & Meisel F. 2006. Hybrid between barn swallow *Hirundo rustica* and sand martin *Riparia riparia*. Limicola. 20(1): 26-31.
65. Walsh J., Billerman S.M., Rohwer V.G., Butcher B.G. & Lovette I.J. 2020. Genomic and plumage variation across the controversial Baltimore and Bullock's oriole hybrid zone. The Auk. 137: 1–15.
66. Terrell M.M. 2020. Dark-eyed junco x white-throated sparrow hybrid in Chisago County. Loon. 92(3): 138.
67. Quintela, M., Thulin, C.G. and Höglund, J., 2010. Detecting hybridization between willow grouse (*Lagopus lagopus*) and rock ptarmigan (*L. muta*) in Central Sweden through Bayesian admixture analyses and mtDNA screening. Conservation Genetics, 11, pp.557-569.
68. Titeux, N. 2010. Nidification d'un hybride Pie-grièche écorcheur × Pie-grièche à tête rousse, *Lanius collurio* × *L. senator*. In Jacob, J.-P., Dehem, C., Burnel, A., Dambiermont, J.-L., Fasol, M., Kinet, T., van der Elst, D. & Paquet, J.-Y. Atlas des Oiseaux Nicheurs de Wallonie 2001–2007. Gembloux: Aves & Région wallonne.
69. Ralston J., Ermacor D.A. & Kirchman J.J. 2015. A new wood warbler hybrid (*Oreothlypis celata x O. ruficapilla*) from the Adirondack mountains, New York. The Wilson Journal of Ornithology. 127(1): 21-28.
70. Summers, R.W., Dawson, R.J. and Phillips, R.E., 2007. Assortative mating and patterns of inheritance indicate that the three crossbill taxa in Scotland are species. Journal of Avian biology, 38(2), pp.153-162.
71. Reifová, R., Kverek, P. and Reif, J., 2011. The first record of a female hybrid between the Common Nightingale (*Luscinia megarhynchos*) and the Thrush Nightingale (*Luscinia luscinia*) in nature. Journal of Ornithology, 152, pp.1063-1068.
72. Kleven, O., Brøseth, H., Jonassen, K. and Pedersen, H.C., 2020. Backcrossing of a capercaillie× black grouse hybrid male in the wild revealed with molecular markers. European Journal of Wildlife Research, 66(2), p.35.
73. Rohwer S, Wood CS, Peters JL, Miller ET, Cagley D, Butcher BG, et al. (2022) Interspecific forced copulations generate most hybrids in broadly sympatric ducks. PLoS ONE 17(9): e0274059. https://doi.org/10.1371/journal.pone.0274059
74. Gehlbach F.R. 2003. Body size variation and evolutionary ecology of eastern and western screech-owls. The Southwestern Naturalist. 48(1): 70–80.
75. Charlesworth C. 2003. Apparent hybrid Lincoln's x song sparrow at Boundary Lake. British Columbia Birds. 13: 10.
76. Johnson R.R. & Hopp S.L. 2010. The First Reported Hybridization of Abert's and Canyon Towhees (*Pipilo* spp.). The Wilson Journal of Ornithology. 122(2): 399-402.
77. Tønnessen D. 2020. Putative canyon towhee x spotted towhee: a new intergeneric hybrid. Western Birds. 51: 263–267.
78. Literák, I., Skyrpan, M. and Andreyenkova, N.G., 2020. A mixed pair of black and red kites in Ukraine, including DNA analysis of hybrid offspring. Biologia, 75, pp.115-120.
79. Platter-Rieger M.F., Shepherd T.M. & Molloy M. Hybridisation of a yellow-crowned and black-crowned night-heron in Southern California. Western Birds. 39: 233–241.
80. Sánchez, M.I., Green, A.J. and Dolz, J.C., 2000. The diets of the White-headed Duck *Oxyura leucocephala*, Ruddy Duck *O. jamaicensis* and their hybrids from Spain. Bird Study, 47(3), pp.275-284.
81. Päckert, M., Ait Belkacem, A., Wolfgramm, H., Gast, O., Canal, D., Giacalone, G., Lo Valvo, M., Vamberger, M., Wink, M., Martens, J. and Stuckas, H., 2019. Genetic admixture despite ecological segregation in a North African sparrow hybrid zone (Aves, Passeriformes, *Passer domesticus*× *Passer hispaniolensis*). Ecology and Evolution, 9(22), pp.12710-12726.

82. Solberg, E.J., Jensen, H., Ringsby, T.H. and Sæther, B.E., 2006. Fitness consequences of hybridization between house sparrows (*Passer domesticus*) and tree sparrows (*P. montanus*). Journal of Ornithology, 147, pp.504-506.
83. Carling M.D. & Zuckerberg B. 2011. Spatio-temporal changes in the genetic structure of the *Passerina* bunting hybrid zone. Molecular Ecology. 20: 1166-1175.
84. Black-headed grosbeaks (*Pheucticus melanocephalus*) and rose-breasted grosbeaks (*Pheucticus ludovicianus*) are passerine bird species known to hybridize in the Great Plains of North America.
85. Petersson, A., Bergner, A.D.A.M. and Thorin, M., 2014. A hybrid Common Redstart x Black Redstart (*Phoenicurus phoenicurus x P. ochruros*) breeding in southeastern Sweden. Ornis svecica, 24(1–2), pp.35-40.
86. Hogner, S., Riera, A.B., Wold, M., Lifjeld, J.T. and Johnsen, A., 2015. Intergeneric hybridization between Common Redstart *Phoenicurus phoenicurus* and Whinchat *Saxicola rubetra* revealed by molecular analyses. Journal of Ornithology, 156, pp.829-836.
87. Dietzen, C., Hackenberg, C., Heyne, K.H., Sauer-Gürth, H., Staudter, H. and Wink, M., 2007. Genetically confirmed interbreeding between Western Bonelli's Warbler (*Phylloscopus bonelli*) and Wood Warbler (*P. sibilatrix*). Journal of Ornithology, 148, pp.85-90.
88. Bensch, S., Helbig, A.J., Salomon, M. and Seibold, I., 2002. Amplified fragment length polymorphism analysis identifies hybrids between two subspecies of warblers. Molecular Ecology, 11(3), pp.473-481.
89. Ławicki Ł., Cofta T., Beuch S., Dmoch A., Sikora A., Aftyka S., Czechowski P., Bocheński M., Sieczak K., Mazgaj S. 2015. Identification and occurrence of Grey-headed × European Green Woodpecker hybrids in Poland. Dutch Birding, 37: 215-228.
90. Backlund D. & Bessken C. 2010. A possible hybrid scarlet/western tanager from the Slim Buttes in Harding County. South Dakota Bird Notes. 57(3): 59-60.
91. Pandolfino E.R., Risser P. & Risser L. 2010. First evidence suggesting hybridisation between the summer tanager and western tanager. Western Birds. 41: 181–183.
92. Mattsson J.P. 2007. A nesting summer tanager X scarlet tanager in Minnesota. The Loon. 79(1): 7-10.
93. Oswald J.A., Harvey M.G., Remsen R.C., Foxworth D.U., Dittmann D.L., Cardiff S.W. & Brum R.T. 2018. Evolutionary dynamics of hybridization and introgression following the recent colonization of Glossy Ibis (Aves: *Plegadis falcinellus*) into the New World. Molecular Ecology. 28: 1675–1691.
94. Bronson C.L., Grubb T.C., Sattler G.D. & Braun M.J. 2005. Reproductive success across the black-capped chickadee (*Poecile atricapillus*) and Carolina chickadee (*P. carolinensis*) hybrid zone in Ohio. The Auk. 122(3): 759–772.
95. Grava A., Otter K.A., Grava T., La Zerte S., Poesel A. & Rush A. 2013. Character displacement in dawn chorusing behaviour of sympatric mountain and black-capped chickadees. Animal Behaviour. 86: 177-187.
96. Lait L.A., Lauff R.F. & Burg T.M. 2012. Genetic evidence supports boreal chickadee (*Poecile hudsonicus*) × black-capped chickadee (P*oecile atricapillus*) hybridization in Atlantic Canada. The Canadian Field Naturalist. 126(2): 143-147.
97. Graham B.A., Gazeley I., Otter K.A. & Burg T. 2021. Do phylogeny and habitat influence admixture among four North American chickadee (family: Paridae) species? Journal of Avian Biology. E02695.
98. Viviano, A. and Mori, E., 2021. Population counts and potential impact of two successful invaders in a town of Northern Italy: the case of ring-necked parakeet and Alexandrine parakeet in Reggio Emilia. Natural History Sciences, 8(1), pp.35-40.
99. Powell A.F.L.A. & Kirkley J.S. 2021. An instance of hybridization with Common Grackle (*Quiscalus quiscula*) and male parental care by Great-tailed Grackle (*Q. mexicanus*). The Wilson Journal of Ornithology. 133(1): 73-81.
100. Coster S.S., Welsh A.B., Costanzo G., Harding S.R., Anderson J.T., McRae S.B. & Katzner T.E. Genetic analyses reveal cryptic introgression in secretive marsh bird populations. Ecology and Evolution. 8: 9870–9879.
101. Pipelow N.D., Leukering T. & Coley E. 2008. An apparent hybrid black x eastern phoebe from Colorado. Western Birds. 39(4): 209-219.
102. Rowe R.A. & Rowe L.M. 2019. Probable cerulean warbler x northern parula hybrid in Rockbridge County, Virginia in April 2019. Banisteria. 53: 72-77.
103. Anich, N.M., Swelstad, J.A. and Pierce, B., 2012. The first report of Sutton's Warbler (Northern Parula× Yellow-throated Warbler) in Wisconsin. Passenger Pigeon, 74, pp.149-155.

Supplementary Table 14. Recording credits for audio files of European species downloaded from Xeno canto (XC) and the Macaulay Library (ML).

| Species | File | Recordist | Species | File | Recordist |
|---|---|---|---|---|---|
| *Acanthis flammea* | XC216985 | Janne Bruun | *Gulosus aristotelis* | XC914590 | Aku Kalliomäki |
| *Acanthis flammea* | XC687649 | Niels Van Doninck | *Gypaetus barbatus* | XC144936 | Fernand Deroussen |
| *Acanthis flammea* | XC710795 | Lars Edenius | *Gypaetus barbatus* | XC569323 | Stanislas Wroza |
| *Acanthis flammea* | XC712649 | Lars Edenius | *Gyps fulvus* | XC423956 | Stanislas Wroza |
| *Accipiter gentilis* | XC756188 | brickegickel | *Gyps fulvus* | XC542056 | Pere Josa |
| *Accipiter gentilis* | XC817762 | Lars Edenius | *Gyps fulvus* | XC653855 | Eduardo Realinho |
| *Accipiter nisus* | XC665236 | Eetu Paljakka | *Gyps fulvus* | XC915617 | Sven Normant |
| *Accipiter nisus* | XC744165 | Romuald Mikusek | *Haematopus ostralegus* | XC831730 | W. Agster |
| *Acrocephalus arundinaceus* | XC811727 | Ulf Elman | *Haematopus ostralegus* | XC942096 | Arjun Dutta |
| *Acrocephalus arundinaceus* | XC813097 | Lukas Thiess | *Haliaeetus albicilla* | XC619160 | Stein Ø. Nilsen |
| *Acrocephalus arundinaceus* | XC813205 | Peter Boesman | *Haliaeetus albicilla* | XC745843 | Lars Edenius |
| *Acrocephalus arundinaceus* | XC815947 | Guy Kirwan | *Hieraaetus pennatus* | XC430828 | Joost van Bruggen |
| *Acrocephalus dumetorum* | XC811764 | Dag Österlund | *Hieraaetus pennatus* | XC668479 | Eduardo Realinho |
| *Acrocephalus dumetorum* | XC813038 | Marcus Fransson | *Himantopus himantopus* | XC828544 | BirdingAlbufera |
| *Acrocephalus dumetorum* | XC814045 | Anders Westman | *Himantopus himantopus* | XC905801 | Albert Lastukhin |
| *Acrocephalus dumetorum* | XC842258 | Ulf Elman | *Hippolais icterina* | XC659733 | Bodo Sonnenburg |
| *Acrocephalus melanopogon* | XC793095 | Stanislas Wroza | *Hippolais icterina* | XC750858 | Ulf Elman |
| *Acrocephalus melanopogon* | XC872907 | Thierry Thomas | *Hippolais icterina* | XC800661 | Stanislas Wroza |
| *Acrocephalus melanopogon* | XC879400 | Manuel Oudard | *Hippolais icterina* | XC815416 | Thomas Bergman |
| *Acrocephalus melanopogon* | XC891244 | Bernard Bousquet | *Hippolais polyglotta* | XC560575 | Jorge Leitão |
| *Acrocephalus paludicola* | XC480838 | Dries Van de Loock | *Hippolais polyglotta* | XC560590 | Jorge Leitão |
| *Acrocephalus paludicola* | XC561499 | Jarek Matusiak | *Hippolais polyglotta* | XC909833 | Jacobo Ramil Millarengo |
| *Acrocephalus paludicola* | XC688954 | Romuald Mikusek | *Hippolais polyglotta* | XC910101 | Olivier Swift, Hélène Cartaud |
| *Acrocephalus paludicola* | XC692253 | Romuald Mikusek | *Hirundo rustica* | XC823733 | Francesco Sottile |
| *Acrocephalus palustris* | XC862700 | Mats Rellmar | *Hirundo rustica* | XC877300 | João Tomás |
| *Acrocephalus palustris* | XC871930 | Mauro Fioretto | *Hirundo rustica* | XC903563 | Stanislas Wroza |
| *Acrocephalus palustris* | XC883779 | Johan Roeland | *Hirundo rustica* | XC910186 | Uku Paal |
| *Acrocephalus palustris* | XC892416 | Alain Malengreau | *Iduna opaca* | XC414658 | Stanislas Wroza |
| *Acrocephalus schoenobaenus* | XC870841 | Alain Malengreau | *Iduna opaca* | XC414663 | Stanislas Wroza |
| *Acrocephalus schoenobaenus* | XC871701 | Alain Malengreau | *Iduna opaca* | XC467771 | Thijs Fijen |
| *Acrocephalus schoenobaenus* | XC884922 | Olivier Swift | *Iduna opaca* | XC798474 | Azens |
| *Acrocephalus schoenobaenus* | XC895063 | Martin Billard | *Ixobrychus minutus* | XC656126 | Lorenzo Maffezzoli |
| *Acrocephalus scirpaceus* | XC862813 | Olivier Swift | *Ixobrychus minutus* | XC836679 | Esperanza Poveda |
| *Acrocephalus scirpaceus* | XC863306 | Chèvremont Fabian | *Jynx torquilla* | XC769997 | Michel Veldt |
| *Acrocephalus scirpaceus* | XC897725 | Lee Alder | *Jynx torquilla* | XC902613 | Ulf Elman |
| *Acrocephalus scirpaceus* | XC901326 | Susanne Kuijpers | *Lagopus lagopus* | XC591861 | Simon Elliott |
| *Actitis hypoleucos* | XC815419 | Thomas Bergman | *Lagopus lagopus* | XC652923 | Thomas Bergman |
| *Actitis hypoleucos* | XC824403 | Romuald Mikusek | *Lagopus lagopus* | XC798642 | Uku Paal |
| *Actitis hypoleucos* | XC834283 | Paolo Zucca | *Lagopus lagopus* | XC900335 | Uku Paal |
| *Actitis hypoleucos* | XC861917 | Peter Stronach | *Lagopus muta* | XC26647 | Patrik Åberg |
| *Aegithalos caudatus* | XC645359 | Regina Eidner | *Lagopus muta* | XC340677 | Tero Linjama |
| *Aegithalos caudatus* | XC707059 | Shamgar Brook | *Lagopus muta* | XC420429 | Stein Ø. Nilsen |
| *Aegolius funereus* | XC708202 | Thomas Bergman | *Lagopus muta* | XC835510 | Petter Westberg |
| *Aegolius funereus* | XC837555 | Romuald Mikusek | *Lanius collurio* | XC560390 | Stanislas Wroza |
| *Aegypius monachus* | XC144935 | Fernand Deroussen | *Lanius collurio* | XC700280 | Dominique Guillerme |
| *Aegypius monachus* | XC915625 | Sven Normant | *Lanius collurio* | XC731753 | Marc Anderson |
| *Aix galericulata* | XC389998 | Jarek Matusiak | *Lanius collurio* | XC803339 | Lars Edenius |
| *Aix galericulata* | XC389999 | Jarek Matusiak | *Lanius excubitor* | XC649962 | Johan Södercrantz |
| *Aix sponsa* | XC509715 | Sunny Tseng | *Lanius excubitor* | XC712619 | Lars Edenius |
| *Aix sponsa* | XC63109 | Jonathon Jongsma | *Lanius excubitor* | XC838997 | Leander Khil |
| *Alauda arvensis* | XC890349 | Olivier Swift | *Lanius excubitor* | XC855983 | Mats Rellmar |
| *Alauda arvensis* | XC893507 | Krzysztof Deoniziak | *Lanius meridionalis* | XC150493 | Paulo Alves |
| *Alauda arvensis* | XC895628 | Agris Celmins | *Lanius meridionalis* | XC727014 | Magnus Wadstein |
| *Alauda arvensis* | XC896982 | Martin Billard | *Lanius meridionalis* | XC767941 | Stanislas Wroza |
| *Alaudala rufescens* | XC730308 | Stanislas Wroza | *Lanius meridionalis* | XC795282 | Étienne Leroy |
| *Alaudala rufescens* | XC730310 | Stanislas Wroza | *Lanius minor* | XC295984 | Marco Dragonetti |
| *Alaudala rufescens* | XC730383 | Stanislas Wroza | *Lanius minor* | XC342809 | Lars Buckx |
| *Alaudala rufescens* | XC756693 | SonoNatura | *Lanius minor* | XC813442 | Peter Boesman |
| *Alcedo atthis* | XC845635 | Johannes Dag Mayer | *Lanius minor* | XC904703 | David Darrell-Lambert |
| *Alcedo atthis* | XC936832 | Johannes Dag Mayer | *Lanius senator* | XC561941 | François Grimal |
| *Alectoris barbara* | ML71168 | Arnoud B. van den Berg | *Lanius senator* | XC722557 | SonoNatura |
| *Alectoris barbara* | XC134392 | Fernand Deroussen | *Lanius senator* | XC754423 | Stein Ø. Nilsen |
| *Alectoris barbara* | XC347457 | Tero Linjama | *Lanius senator* | XC797821 | Xavier Riera |
| *Alectoris barbara* | XC901725 | Andrew Harrop | *Leiothrix lutea* | ML14175 | William V. Ward |
| *Alectoris graeca* | XC182904 | Manuel Schweizer | *Leiothrix lutea* | ML445568141 | Sujith S |
| *Alectoris graeca* | XC247001 | Jerome Fischer | *Leiothrix lutea* | ML580194941 | Benjamin Costello |
| *Alectoris graeca* | XC416855 | Jerome Fischer | *Leiothrix lutea* | ML600224591 | Ray Tsu |
| *Alectoris graeca* | XC730580 | Stanislas Wroza | *Limosa lapponica* | XC654136 | Lars Edenius |
| *Alectoris rufa* | XC265614 | david m | *Limosa lapponica* | XC807724 | Thomas Bergman |
| *Alectoris rufa* | XC334156 | Jordi Calvet | *Limosa lapponica* | XC913545 | Thomas Bergman |
| *Alectoris rufa* | XC372530 | Stanislas Wroza | *Limosa lapponica* | XC915090 | Thomas Bergman |
| *Alectoris rufa* | XC794515 | Stuart Fisher | *Limosa limosa* | XC722732 | Lars Edenius |
| *Alopochen aegyptiaca* | XC546108 | Albert Noorlander | *Limosa limosa* | XC740254 | Uku Paal |
| *Alopochen aegyptiaca* | XC620525 | Francesco Sottile | *Limosa limosa* | XC740270 | Uku Paal |
| *Amandava amandava* | XC545983 | Peter Ericsson | *Limosa limosa* | XC818972 | Azens |
| *Amandava amandava* | XC552807 | Bram Piot | *Linaria cannabina* | XC802274 | Thomas Bergman |
| *Amandava amandava* | XC824847 | He Wenjin | *Linaria cannabina* | XC876858 | João Tomás |
| *Amandava amandava* | XC840610 | Bram Piot | *Linaria cannabina* | XC899180 | Paul Kelly |
| *Anas acuta* | XC712335 | Jack Berteau | *Linaria cannabina* | XC906691 | Olivier Swift |
| *Anas acuta* | XC721976 | Lars Edenius | *Linaria flavirostris* | XC364664 | Elias A. Ryberg |
| *Anas crecca* | XC668192 | Ireneusz Oleksik | *Linaria flavirostris* | XC470797 | Dries Van de Loock |
| *Anas crecca* | XC732343 | Marc Anderson | *Linaria flavirostris* | XC638923 | Lars Edenius |
| *Anas platyrhynchos* | XC755991 | Romuald Mikusek | *Linaria flavirostris* | XC638934 | Lars Edenius |
| *Anas platyrhynchos* | XC774431 | Irish Wildlife Sounds | *Locustella fluviatilis* | XC861724 | András Schmidt |
| *Anser albifrons* | XC644998 | Joost van Bruggen | *Locustella fluviatilis* | XC908882 | Eric Söderman |
| *Anser albifrons* | XC698271 | András Schmidt | *Locustella luscinioides* | XC740299 | Uku Paal |

| | | |
|---|---|---|
| *Anser albifrons* | XC712047 | Joost van Bruggen |
| *Anser albifrons* | XC878146 | Lars Mogensen |
| *Anser anser* | XC792700 | Lars Edenius |
| *Anser anser* | XC803358 | Lars Edenius |
| *Anser anser* | XC896889 | Lars Edenius |
| *Anser anser* | XC914627 | Lennart Jeppsson |
| *Anser brachyrhynchus* | XC521246 | Ireneusz Oleksik |
| *Anser brachyrhynchus* | XC601551 | Will Scott |
| *Anser canagicus* | XC185372 | Andrew Spencer |
| *Anser canagicus* | XC486631 | Jens Kirkeby |
| *Anser cygnoides* | XC635985 | Simon Kiesé |
| *Anser cygnoides* | XC765912 | Stanislas Wroza |
| *Anser fabalis* | XC109681 | Jelmer Poelstra |
| *Anser fabalis* | XC676859 | Thomas Bergman |
| *Anser indicus* | XC526948 | James Lidster |
| *Anser indicus* | XC635939 | David Darrell-Lambert |
| *Anthus berthelotii* | XC134391 | Fernand Deroussen |
| *Anthus berthelotii* | XC374980 | Peter Boesman |
| *Anthus berthelotii* | XC764548 | Stein Ø. Nilsen |
| *Anthus berthelotii* | XC782278 | Rubén Barone |
| *Anthus campestris* | XC801392 | Manceau Lionel |
| *Anthus campestris* | XC813237 | Peter Boesman |
| *Anthus campestris* | XC839904 | Corentin Rivière |
| *Anthus campestris* | XC897500 | Tero Linjama |
| *Anthus cervinus* | XC139099 | Fernand Deroussen |
| *Anthus cervinus* | XC345769 | Tero Linjama |
| *Anthus cervinus* | XC382130 | Jens Kirkeby |
| *Anthus cervinus* | XC424100 | Terje Kolaas |
| *Anthus petrosus* | XC146046 | julien Rochefort |
| *Anthus petrosus* | XC263958 | Terje Kolaas |
| *Anthus petrosus* | XC610047 | Lars Edenius |
| *Anthus petrosus* | XC667502 | Irish Wildlife Sounds |
| *Anthus pratensis* | XC779776 | Erik Normark |
| *Anthus pratensis* | XC831852 | Agris Celmins |
| *Anthus pratensis* | XC831853 | Agris Celmins |
| *Anthus pratensis* | XC857639 | Beatrix Saadi-Varchmin |
| *Anthus spinoletta* | XC716570 | Xavier Riera |
| *Anthus spinoletta* | XC730538 | Stanislas Wroza |
| *Anthus spinoletta* | XC734861 | Magnus Hellström |
| *Anthus spinoletta* | XC839878 | Romuald Mikusek |
| *Anthus trivialis* | XC855476 | Jack Berteau |
| *Anthus trivialis* | XC879350 | Stein Ø. Nilsen |
| *Anthus trivialis* | XC895016 | Christian Kerihuel |
| *Anthus trivialis* | XC897400 | Jacobo Ramil Millarengo |
| *Apus apus* | XC810578 | Susanne Kuijpers |
| *Apus apus* | XC817637 | João Tomás |
| *Apus caffer* | ML5989 | Myles E. W. North |
| *Apus caffer* | XC677525 | Esperanza Poveda |
| *Apus pallidus* | XC557421 | Jorge Leitão |
| *Apus pallidus* | XC763779 | Guillaume Bigayon |
| *Apus unicolor* | XC164524 | Antero Lindholm |
| *Apus unicolor* | XC657346 | Paulo Alves |
| *Aquila adalberti* | XC148645 | Patrik Åberg |
| *Aquila adalberti* | XC420390 | José Carlos Sires |
| *Aquila adalberti* | XC699619 | SonoNatura |
| *Aquila adalberti* | XC864217 | SonoNatura |
| *Aquila chrysaetos* | XC341722 | Tero Linjama |
| *Aquila chrysaetos* | XC453324 | Lars Edenius |
| *Aquila chrysaetos* | XC753803 | Lars Edenius |
| *Aquila chrysaetos* | XC820295 | Hans Norelius |
| *Aquila fasciata* | ML411897521 | Fast As Falcon |
| *Aquila fasciata* | XC781530 | Peter Boesman |
| *Aquila heliaca* | XC582823 | Jarek Matusiak |
| *Aquila heliaca* | XC731082 | brickegickel |
| *Ardea alba* | XC615283 | Stanislas Wroza |
| *Ardea alba* | XC681080 | Uku Paal |
| *Ardea alba* | XC738072 | Marcin Sołowiej |
| *Ardea alba* | XC776170 | Irish Wildlife Sounds |
| *Ardea cinerea* | XC857972 | Martin Billard |
| *Ardea cinerea* | XC881881 | Esperanza Poveda |
| *Ardea cinerea* | XC882189 | Christian Vemmelund Helligsø |
| *Ardea cinerea* | XC897660 | Paul Kelly |
| *Ardea purpurea* | ML568281661 | Anonymous |
| *Ardea purpurea* | XC485377 | Joost van Bruggen |
| *Ardea purpurea* | XC903478 | Stanislas Wroza |
| *Ardea purpurea* | XC903564 | Stanislas Wroza |
| *Ardeola ralloides* | XC577793 | Cedric Mroczko |
| *Ardeola ralloides* | XC836622 | Esperanza Poveda |
| *Arenaria interpres* | XC815489 | Sonothèque ADVL |
| *Arenaria interpres* | XC815490 | Sonothèque ADVL |
| *Arenaria interpres* | XC827973 | Jorge Leitão |
| *Arenaria interpres* | XC895259 | Francesco Sottile |
| *Asio flammeus* | XC718743 | Lars Edenius |
| *Asio flammeus* | XC731000 | Lars Edenius |
| *Asio otus* | XC793292 | Dominique Guillerme |
| *Asio otus* | XC793294 | Dominique Guillerme |
| *Athene noctua* | XC883232 | Anthony Roux |
| *Athene noctua* | XC891838 | András Schmidt |
| *Athene noctua* | XC908597 | Martin Billard |
| *Athene noctua* | XC915445 | Esperanza Poveda |
| *Locustella luscinioides* | XC813023 | Lukas Thiess |
| *Locustella naevia* | XC900840 | Martin Billard |
| *Locustella naevia* | XC900843 | Martin Billard |
| *Lophophanes cristatus* | XC780340 | Lars Edenius |
| *Lophophanes cristatus* | XC790414 | Lars Edenius |
| *Lophophanes cristatus* | XC804854 | Uku Paal |
| *Lophophanes cristatus* | XC896344 | Jacobo Ramil Millarengo |
| *Loxia curvirostra* | XC831945 | Lars Edenius |
| *Loxia curvirostra* | XC837909 | Lars Edenius |
| *Loxia curvirostra* | XC864185 | András Schmidt |
| *Loxia curvirostra* | XC901928 | Lars Edenius |
| *Loxia leucoptera* | XC338457 | Piotr Szczypinski |
| *Loxia leucoptera* | XC484760 | Teet Sirotkin |
| *Loxia leucoptera* | XC602666 | Christopher McPherson |
| *Loxia leucoptera* | XC89151 | Patrik Åberg |
| *Loxia pytyopsittacus* | XC403068 | Hans Matheve |
| *Loxia pytyopsittacus* | XC829642 | Lars Edenius |
| *Loxia pytyopsittacus* | XC893254 | Lars Edenius |
| *Loxia pytyopsittacus* | XC900834 | Johan Forssell |
| *Loxia scotica* | XC544934 | Peter Stronach |
| *Loxia scotica* | XC715938 | Andrew Harrop |
| *Loxia scotica* | XC892394 | Peter Stronach |
| *Loxia scotica* | XC906960 | Peter Stronach |
| *Lullula arborea* | XC712592 | Joost van Bruggen |
| *Lullula arborea* | XC722492 | Lars Edenius |
| *Lullula arborea* | XC723523 | Olivier Swift, Sylvain REYT |
| *Lullula arborea* | XC852698 | Rob van Bemmelen |
| *Luscinia luscinia* | XC654939 | maxander |
| *Luscinia luscinia* | XC727140 | Lennart Jeppsson |
| *Luscinia luscinia* | XC803654 | Johannes Sander |
| *Luscinia luscinia* | XC903880 | Jocce Ekstrom |
| *Luscinia megarhynchos* | XC803656 | Johannes Sander |
| *Luscinia megarhynchos* | XC901015 | Grzegorz Lorek |
| *Luscinia megarhynchos* | XC908425 | Leconte Michel |
| *Luscinia megarhynchos* | XC909501 | Mats Rellmar |
| *Luscinia svecica* | XC707296 | Antonio Xeira |
| *Luscinia svecica* | XC731887 | SonoNatura |
| *Luscinia svecica* | XC884116 | Maarten Sluijter |
| *Luscinia svecica* | XC899983 | Jörgen Pisch |
| *Lymnocryptes minimus* | XC723297 | Lars Edenius |
| *Lymnocryptes minimus* | XC799634 | Lars Edenius |
| *Lyrurus tetrix* | XC711208 | Baltasar Pinheiro |
| *Lyrurus tetrix* | XC758859 | Lars Edenius |
| *Lyrurus tetrix* | XC758861 | Lars Edenius |
| *Lyrurus tetrix* | XC805524 | Lars Edenius |
| *Mareca penelope* | XC767302 | Irish Wildlife Sounds |
| *Mareca penelope* | XC882155 | David Darrell-Lambert |
| *Mareca strepera* | XC679027 | Beatrix Saadi-Varchmin |
| *Mareca strepera* | XC871361 | Paul Kelly |
| *Marmaronetta angustirostris* | XC780493 | Santiago Caballero Carrera |
| *Marmaronetta angustirostris* | XC784480 | Santiago Caballero Carrera |
| *Melanitta fusca* | XC448663 | Patrik Åberg |
| *Melanitta fusca* | XC817948 | Alan Dalton |
| *Melanitta nigra* | XC902773 | david thorns |
| *Melanitta nigra* | XC915469 | Thomas Bergman |
| *Melanocorypha calandra* | XC510598 | Marco Dragonetti |
| *Melanocorypha calandra* | XC730261 | Stanislas Wroza |
| *Melanocorypha calandra* | XC730301 | Stanislas Wroza |
| *Melanocorypha calandra* | XC880726 | Luis Gracia |
| *Mergellus albellus* | XC563997 | Lars Edenius |
| *Mergellus albellus* | XC610927 | Lars Edenius |
| *Mergus merganser* | XC644424 | Lars Edenius |
| *Mergus merganser* | XC759927 | Simon Elliott |
| *Mergus serrator* | XC607514 | Peter Stronach |
| *Mergus serrator* | XC762330 | Simon Elliott |
| *Merops apiaster* | XC824573 | Grzegorz Lorek |
| *Merops apiaster* | XC900856 | João Tomás |
| *Microcarbo pygmaeus* | XC168146 | Andre et Odile Boucher |
| *Microcarbo pygmaeus* | XC296075 | Marco Dragonetti |
| *Milvus migrans* | XC796153 | Beatrix Saadi-Varchmin |
| *Milvus migrans* | XC899786 | Jorge Leitão |
| *Milvus milvus* | XC829695 | Olivier Swift |
| *Milvus milvus* | XC879195 | Olivier Swift |
| *Monticola saxatilis* | XC296005 | Marco Dragonetti |
| *Monticola saxatilis* | XC649771 | Jordi Calvet |
| *Monticola saxatilis* | XC730219 | Stanislas Wroza |
| *Monticola saxatilis* | XC833863 | Martin Billard |
| *Monticola solitarius* | XC265241 | Michele Viganò |
| *Monticola solitarius* | XC417708 | Stanislas Wroza |
| *Monticola solitarius* | XC881595 | Francesco Sottile |
| *Monticola solitarius* | XC99691 | Jordi Calvet |
| *Montifringilla nivalis* | XC141184 | Fernand Deroussen |
| *Montifringilla nivalis* | XC186477 | Michele Peron |
| *Montifringilla nivalis* | XC355444 | maudoc |
| *Montifringilla nivalis* | XC486810 | Audevard Aurélien |
| *Motacilla alba* | XC598489 | Tanguy Loïs |
| *Motacilla alba* | XC656706 | Uku Paal |
| *Motacilla alba* | XC656713 | Uku Paal |
| *Motacilla alba* | XC722531 | Jens Kirkeby |
| *Motacilla cinerea* | XC682324 | Jacobo Ramil MIllarengo |

| | | |
|---|---|---|
| *Aythya ferina* | XC769846 | Dominique Guillerme |
| *Aythya ferina* | XC797060 | Alan Dalton |
| *Aythya fuligula* | XC644316 | Lars Edenius |
| *Aythya fuligula* | XC644396 | Lars Edenius |
| *Aythya marila* | ML137835 | Gerrit Vyn |
| *Aythya marila* | ML444104281 | Reid Hildebrandt |
| *Aythya nyroca* | XC296088 | Marco Dragonetti |
| *Aythya nyroca* | XC528021 | Jan Daniels-Trautner |
| *Bombycilla garrulus* | XC597940 | Lars Edenius |
| *Bombycilla garrulus* | XC861580 | Peter Stronach |
| *Botaurus stellaris* | XC465220 | Gregorio Para |
| *Botaurus stellaris* | XC720112 | Manuel Grosselet |
| *Branta canadensis* | XC646159 | Lars Edenius |
| *Branta canadensis* | XC715941 | Agris Celmins |
| *Branta canadensis* | XC787100 | Susanne Kuijpers |
| *Branta canadensis* | XC909895 | Uku Paal |
| *Branta hutchinsii* | ML135491 | William W. H. Gunn |
| *Branta hutchinsii* | ML203928861 | Josep del Hoyo |
| *Branta hutchinsii* | XC361080 | Paul Marvin |
| *Branta hutchinsii* | XC838299 | Seth Beaudreault (Toolik Field Station) |
| *Branta leucopsis* | XC549292 | Lars Edenius |
| *Branta leucopsis* | XC724651 | Uku Paal |
| *Branta leucopsis* | XC763873 | Uku Paal |
| *Branta leucopsis* | XC903305 | Lars Edenius |
| *Bubo bubo* | XC836565 | Thierry Thomas |
| *Bubo bubo* | XC896167 | Bram Vogels |
| *Bubo scandiacus* | XC343144 | Tero Linjama |
| *Bubo scandiacus* | XC343145 | Tero Linjama |
| *Bubulcus ibis* | XC356329 | Marco Dragonetti |
| *Bubulcus ibis* | XC495869 | Joost van Bruggen |
| *Bubulcus ibis* | XC76666 | Marco Dragonetti |
| *Bubulcus ibis* | XC831486 | Jordi Calvet |
| *Bucanetes githagineus* | XC164199 | Tero Linjama |
| *Bucanetes githagineus* | XC855968 | Mats Rellmar |
| *Bucephala clangula* | XC234677 | dmitry yakubovich |
| *Bucephala clangula* | XC641760 | Lars Edenius |
| *Burhinus oedicnemus* | XC896992 | Martin Billard |
| *Burhinus oedicnemus* | XC914574 | Thierry Thomas |
| *Burhinus oedicnemus* | XC923183 | Jordi Calvet |
| *Burhinus oedicnemus* | XC934316 | Peter Ertl |
| *Buteo buteo* | XC878780 | João Tomás |
| *Buteo buteo* | XC895030 | Martin Billard |
| *Buteo lagopus* | XC665960 | Lars Edenius |
| *Buteo lagopus* | XC747268 | Lars Edenius |
| *Cairina moschata* | ML92424851 | Paul Marvin |
| *Cairina moschata* | ML92475411 | Paul Marvin |
| *Calandrella brachydactyla* | XC806185 | Pere Josa |
| *Calandrella brachydactyla* | XC813260 | Peter Boesman |
| *Calandrella brachydactyla* | XC813265 | Peter Boesman |
| *Calandrella brachydactyla* | XC839296 | Corentin Rivière |
| *Calcarius lapponicus* | XC567566 | Lars Edenius |
| *Calcarius lapponicus* | XC580134 | Niels Van Doninck |
| *Calcarius lapponicus* | XC639055 | Peter Stronach |
| *Calcarius lapponicus* | XC688953 | Lars Edenius |
| *Calidris alpina* | XC739138 | Uku Paal |
| *Calidris alpina* | XC833015 | Michaël Bridoux |
| *Calidris alpina* | XC852842 | Rob van Bemmelen |
| *Calidris alpina* | XC908418 | Lars Edenius |
| *Calidris falcinellus* | XC654231 | Lars Edenius |
| *Calidris falcinellus* | XC911575 | Niclas Backstrom |
| *Calidris maritima* | XC186400 | Stein Ø. Nilsen |
| *Calidris maritima* | XC321244 | Tim de Boer |
| *Calidris maritima* | XC322089 | Rob van Bemmelen |
| *Calidris maritima* | XC423507 | Johan Råghall |
| *Calidris minuta* | XC383563 | Terje Kolaas |
| *Calidris minuta* | XC383564 | Terje Kolaas |
| *Calidris pugnax* | XC765283 | Stanislas Wroza |
| *Calidris pugnax* | XC852294 | Jordi Calvet |
| *Calidris temminckii* | XC765454 | Stanislas Wroza |
| *Calidris temminckii* | XC809521 | Frode Falkenberg |
| *Caprimulgus europaeus* | XC809979 | Olivier Swift |
| *Caprimulgus europaeus* | XC957127 | Jarek Matusiak |
| *Caprimulgus ruficollis* | XC724307 | Antonio Xeira |
| *Caprimulgus ruficollis* | XC905834 | SonoNatura |
| *Carduelis carduelis* | XC854554 | David Pennington |
| *Carduelis carduelis* | XC883327 | João Tomás |
| *Carduelis carduelis* | XC884098 | Christian Kerihuel |
| *Carduelis carduelis* | XC894563 | Jorge Leitão |
| *Carduelis citrinella* | XC246416 | Jordi Calvet |
| *Carduelis citrinella* | XC315443 | Jerome Fischer |
| *Carduelis citrinella* | XC372554 | Jerome Fischer |
| *Carduelis citrinella* | XC730229 | Stanislas Wroza |
| *Carduelis corsicana* | XC349876 | Tero Linjama |
| *Carduelis corsicana* | XC624722 | Charlie Bodin |
| *Carduelis corsicana* | XC662052 | Marta Celej |
| *Carduelis corsicana* | XC662053 | Marta Celej |
| *Carpodacus erythrinus* | XC743414 | Romuald Mikusek |
| *Carpodacus erythrinus* | XC863138 | Mats Rellmar |
| *Carpodacus erythrinus* | XC904579 | Lars Edenius |
| *Carpodacus erythrinus* | XC912797 | Hans Matheve |
| *Cecropis daurica* | XC485355 | Lars Lachmann |
| *Cecropis daurica* | XC723730 | Geoff Carey |
| *Motacilla cinerea* | XC757193 | Lars Lachmann |
| *Motacilla cinerea* | XC797916 | brickegickel |
| *Motacilla cinerea* | XC904381 | Cedric Mroczko |
| *Motacilla flava* | XC745680 | Lars Edenius |
| *Motacilla flava* | XC793956 | Jacobo Ramil Millarengo |
| *Motacilla flava* | XC838987 | Regina Eidner |
| *Motacilla flava* | XC881967 | Olivier Swift |
| *Muscicapa striata* | XC598245 | Niels Van Doninck |
| *Muscicapa striata* | XC727741 | Paolo Zucca |
| *Muscicapa striata* | XC780925 | Bas |
| *Muscicapa striata* | XC877767 | Maarten Sluijter |
| *Myiopsitta monachus* | XC622769 | jesus carrion |
| *Myiopsitta monachus* | XC669961 | Manuel Grosselet |
| *Myiopsitta monachus* | XC678007 | Guilherme Renzo Rocha Brito |
| *Myiopsitta monachus* | XC682586 | Manuel Grosselet |
| *Neophron percnopterus* | XC650774 | Eduardo Realinho |
| *Neophron percnopterus* | XC654205 | Eduardo Realinho |
| *Netta rufina* | XC404318 | Pascal Christe |
| *Netta rufina* | XC407509 | Francesco Sottile |
| *Netta rufina* | XC644652 | Audevard Aurélien |
| *Netta rufina* | XC769720 | Michel Veldt |
| *Nucifraga caryocatactes* | XC736823 | Lars Edenius |
| *Nucifraga caryocatactes* | XC818171 | Lars Edenius |
| *Nucifraga caryocatactes* | XC843797 | Richard Brahmstaedt |
| *Nucifraga caryocatactes* | XC906816 | Frode Falkenberg |
| *Numenius arquata* | XC830500 | Lars Edenius |
| *Numenius arquata* | XC832849 | Agris Celmins |
| *Numenius phaeopus* | XC735243 | Lars Edenius |
| *Numenius phaeopus* | XC741684 | Lars Edenius |
| *Numenius phaeopus* | XC799611 | Niels Van Doninck |
| *Numenius phaeopus* | XC801501 | Olivier Swift |
| *Numida meleagris* | XC280506 | Peter Boesman |
| *Numida meleagris* | XC507927 | Tim Cockcroft |
| *Nycticorax nycticorax* | XC903475 | Stanislas Wroza |
| *Nycticorax nycticorax* | XC921344 | Sébastien Arriubergé |
| *Oenanthe hispanica* | XC343294 | Agostinho Tomás |
| *Oenanthe hispanica* | XC416420 | Jordi Calvet |
| *Oenanthe hispanica* | XC730459 | Stanislas Wroza |
| *Oenanthe hispanica* | XC794843 | João Tomás |
| *Oenanthe leucura* | XC538884 | Xavier Riera |
| *Oenanthe leucura* | XC692725 | Santiago Caballero Carrera |
| *Oenanthe leucura* | XC730407 | Stanislas Wroza |
| *Oenanthe leucura* | XC840125 | Pere Josa |
| *Oenanthe oenanthe* | XC759707 | Ulf Elman |
| *Oenanthe oenanthe* | XC769226 | Michel Veldt |
| *Oenanthe oenanthe* | XC769227 | Michel Veldt |
| *Oenanthe oenanthe* | XC898202 | Jochem verweij |
| *Oriolus oriolus* | XC708637 | Agris Celmins |
| *Oriolus oriolus* | XC739166 | Grzegorz Lorek |
| *Oriolus oriolus* | XC743901 | Romuald Mikusek |
| *Oriolus oriolus* | XC809026 | Bodo Sonnenburg |
| *Otis tarda* | XC721833 | Lars Lachmann |
| *Otis tarda* | XC721834 | Lars Lachmann |
| *Otus scops* | XC905326 | Christian Kerihuel |
| *Otus scops* | XC910434 | Antoine Salmon |
| *Oxyura jamaicensis* | XC591765 | Simon Elliott |
| *Oxyura jamaicensis* | XC603727 | Peter Ward and Ken Hall |
| *Oxyura leucocephala* | XC132436 | Fernand Deroussen |
| *Oxyura leucocephala* | XC462959 | Geoffrey Monchaux |
| *Pandion haliaetus* | XC826075 | Olli Juhnke |
| *Pandion haliaetus* | XC924318 | Lars Edenius |
| *Panurus biarmicus* | XC955702 | Jarek Matusiak |
| *Panurus biarmicus* | XC955703 | Jarek Matusiak |
| *Parus major* | XC881427 | Michel Veldt |
| *Parus major* | XC899545 | Beatrix Saadi-Varchmin |
| *Parus major* | XC899637 | Lennart Jeppsson |
| *Parus major* | XC902756 | Olivier Swift |
| *Passer domesticus* | XC879927 | Paul Kelly |
| *Passer domesticus* | XC881740 | Olivier Swift |
| *Passer domesticus* | XC896493 | Martin Billard |
| *Passer domesticus* | XC899351 | Jack Berteau |
| *Passer hispaniolensis* | XC309787 | Cedric Mroczko |
| *Passer hispaniolensis* | XC334279 | Yoann Blanchon |
| *Passer hispaniolensis* | XC905912 | João Tomás |
| *Passer hispaniolensis* | XC913059 | Stephan Risch |
| *Passer italiae* | XC727597 | Francesco Sottile |
| *Passer italiae* | XC906612 | Frederic Lionel |
| *Passer italiae* | XC906614 | Frederic Lionel |
| *Passer italiae* | XC907069 | Mats Rellmar |
| *Passer montanus* | XC800953 | Daniele Baroni |
| *Passer montanus* | XC813881 | Susanne Kuijpers |
| *Passer montanus* | XC814643 | Uku Paal |
| *Passer montanus* | XC815513 | Susanne Kuijpers |
| *Pavo cristatus* | XC812476 | Francesco Barberini |
| *Pavo cristatus* | XC902565 | Kalle Nibbenhagen |
| *Perdix perdix* | XC651353 | Uku Paal |
| *Perdix perdix* | XC897003 | Martin Billard |
| *Periparus ater* | XC891510 | Frederic Lionel |
| *Periparus ater* | XC902395 | Thomas Bergman |
| *Periparus ater* | XC904730 | Jorge Leitão |
| *Periparus ater* | XC910374 | Johan Södercrantz |
| *Perisoreus infaustus* | XC611732 | Lars Edenius |

| | | | | | |
|---|---|---|---|---|---|
| *Cercotrichas galactotes* | XC670255 | Jonathan Bryant | *Perisoreus infaustus* | XC652996 | Thomas Bergman |
| *Cercotrichas galactotes* | XC692573 | Santiago Caballero Carrera | *Perisoreus infaustus* | XC680829 | Lars Edenius |
| *Cercotrichas galactotes* | XC831603 | Esperanza Poveda | *Perisoreus infaustus* | XC751019 | Lars Edenius |
| *Cercotrichas galactotes* | XC831605 | Esperanza Poveda | *Pernis apivorus* | XC329631 | Kaspars Funts |
| *Certhia brachydactyla* | XC899288 | David Pennington | *Pernis apivorus* | XC666557 | P. Venema |
| *Certhia brachydactyla* | XC901386 | Jorge Leitão | *Pernis apivorus* | XC668502 | Sophie Neill |
| *Certhia brachydactyla* | XC901387 | Jorge Leitão | *Pernis apivorus* | XC826297 | Lars Edenius |
| *Certhia brachydactyla* | XC905013 | Olivier Swift | *Petronia petronia* | XC651365 | Marta Celej |
| *Certhia familiaris* | XC808139 | Lars Edenius | *Petronia petronia* | XC692717 | Santiago Caballero Carrera |
| *Certhia familiaris* | XC878886 | Baltasar Pinheiro | *Petronia petronia* | XC783481 | Jacobo Ramil Millarengo |
| *Certhia familiaris* | XC893809 | Azens | *Petronia petronia* | XC783482 | Jacobo Ramil Millarengo |
| *Certhia familiaris* | XC904077 | Corentin Rivière | *Phalacrocorax carbo* | XC646784 | Agris Celmins |
| *Cettia cetti* | XC907063 | Mats Rellmar | *Phalacrocorax carbo* | XC737064 | Irish Wildlife Sounds |
| *Cettia cetti* | XC963920 | Jack Berteau | *Phalacrocorax carbo* | XC776010 | Paul Kelly |
| *Cettia cetti* | XC964596 | Friedrich Richard | *Phalacrocorax carbo* | XC841971 | Jorge Leitão |
| *Cettia cetti* | XC965043 | Jack Berteau | *Phalaropus lobatus* | XC342920 | Tero Linjama |
| *Charadrius alexandrinus* | XC338266 | José Carlos Sires | *Phalaropus lobatus* | XC420149 | Stein Ø. Nilsen |
| *Charadrius alexandrinus* | XC817932 | Sonothèque ADVL | *Phalaropus lobatus* | XC487653 | Stanislas Wroza |
| *Charadrius dubius* | XC813186 | Lars Edenius | *Phalaropus lobatus* | XC514049 | Timo Janhonen |
| *Charadrius dubius* | XC815926 | Albert Lastukhin | *Phasianus colchicus* | XC640749 | Tanguy Loïs |
| *Charadrius hiaticula* | XC653537 | Elias A. Ryberg | *Phasianus colchicus* | XC929867 | Bram Piot |
| *Charadrius hiaticula* | XC653595 | Elias A. Ryberg | *Phoenicopterus roseus* | XC267542 | Andrew Spencer |
| *Charadrius morinellus* | XC440167 | Stein Ø. Nilsen | *Phoenicopterus roseus* | XC899996 | Paul Coiffard |
| *Charadrius morinellus* | XC570939 | Lars Edenius | *Phoenicurus ochruros* | XC832033 | Francisco Molinero |
| *Chersophilus duponti* | XC216990 | Cristian Pérez-Granados | *Phoenicurus ochruros* | XC844168 | Frederic Lionel |
| *Chersophilus duponti* | XC235394 | Luis Gracia | *Phoenicurus ochruros* | XC857226 | Grzegorz Lorek |
| *Chersophilus duponti* | XC315238 | Jarmo Pirhonen | *Phoenicurus ochruros* | XC898801 | Nicolas Martinez |
| *Chersophilus duponti* | XC536806 | Sean Ronayne | *Phoenicurus phoenicurus* | XC899515 | Lars Mogensen |
| *Chloris chloris* | XC814746 | Martin Billard | *Phoenicurus phoenicurus* | XC911881 | Antonio Xeira |
| *Chloris chloris* | XC834605 | Esperanza Poveda | *Phoenicurus phoenicurus* | XC913428 | Martin Billard |
| *Chloris chloris* | XC879027 | João Tomás | *Phoenicurus phoenicurus* | XC913449 | Thomas Bergman |
| *Chloris chloris* | XC898912 | Nicolas Martinez | *Phylloscopus bonelli* | XC769307 | Michel Veldt |
| *Chrysolophus amherstiae* | XC473878 | Albert Lastukhin | *Phylloscopus bonelli* | XC799404 | João Tomás |
| *Chrysolophus amherstiae* | XC916157 | Peter Boesman | *Phylloscopus bonelli* | XC819672 | Jacobo Ramil Millarengo |
| *Chrysolophus pictus* | XC37540 | David Farrow | *Phylloscopus bonelli* | XC892813 | Frederic Lionel |
| *Chrysolophus pictus* | XC908427 | Xingyu Li | *Phylloscopus borealis* | XC401786 | Karl-Birger Strann |
| *Ciconia ciconia* | XC878180 | Мітла Андрій Володимирович | *Phylloscopus borealis* | XC652032 | Lukas Pelikan |
| *Ciconia ciconia* | XC878788 | João Tomás | *Phylloscopus borealis* | XC785236 | Teet Sirotkin |
| *Ciconia nigra* | XC795530 | Sven Normant | *Phylloscopus borealis* | XC896729 | Zbigniew Kajzer |
| *Ciconia nigra* | XC96963 | Edmunds Racinskis | *Phylloscopus canariensis* | XC626845 | Olivier Swift, Ludivine Delamare |
| *Cinclus cinclus* | XC687370 | Lars Edenius | *Phylloscopus canariensis* | XC934197 | Jose |
| *Cinclus cinclus* | XC700209 | Stanislas Wroza | *Phylloscopus collybita* | XC879588 | Susanne Kuijpers |
| *Cinclus cinclus* | XC700213 | Stanislas Wroza | *Phylloscopus collybita* | XC900042 | Thomas Bergman |
| *Cinclus cinclus* | XC883370 | Teet Sirotkin | *Phylloscopus collybita* | XC904320 | Martin Billard |
| *Circaetus gallicus* | XC40049 | Doug Knapp | *Phylloscopus collybita* | XC908343 | Olivier Swift, Hélène Cartaud |
| *Circaetus gallicus* | XC587198 | Eduardo Realinho | *Phylloscopus ibericus* | XC794962 | Jacobo Ramil Millarengo |
| *Circus aeruginosus* | XC859056 | João Tomás | *Phylloscopus ibericus* | XC894453 | Pere Josa |
| *Circus aeruginosus* | XC884134 | João Tomás | *Phylloscopus ibericus* | XC896346 | Jacobo Ramil Millarengo |
| *Circus cyaneus* | XC341267 | Tero Linjama | *Phylloscopus ibericus* | XC909268 | Pere Josa |
| *Circus cyaneus* | XC670733 | Stanislas Wroza | *Phylloscopus sibilatrix* | XC729301 | Bastian Forkel |
| *Circus macrourus* | XC471461 | Stanislas Wroza | *Phylloscopus sibilatrix* | XC769998 | Michel Veldt |
| *Circus macrourus* | XC643655 | Hans Matheve | *Phylloscopus sibilatrix* | XC840441 | Rob van Bemmelen |
| *Circus pygargus* | XC433734 | José Carlos Sires | *Phylloscopus sibilatrix* | XC911177 | W. Agster |
| *Circus pygargus* | XC573274 | Jarek Matusiak | *Phylloscopus trochiloides* | XC802971 | Jens Kirkeby |
| *Cisticola juncidis* | XC930481 | Esperanza Poveda | *Phylloscopus trochiloides* | XC810536 | Romuald Mikusek |
| *Cisticola juncidis* | XC962299 | Hans Matheve | *Phylloscopus trochiloides* | XC813612 | Peter Boesman |
| *Clamator glandarius* | XC708536 | Sergi Carreras | *Phylloscopus trochiloides* | XC913921 | Lars Edenius |
| *Clamator glandarius* | XC724315 | Alexander Schlatmann | *Phylloscopus trochilus* | XC801762 | Lars Edenius |
| *Clamator glandarius* | XC877435 | João Tomás | *Phylloscopus trochilus* | XC855110 | David Pennington |
| *Clamator glandarius* | XC877437 | João Tomás | *Phylloscopus trochilus* | XC905213 | Azens |
| *Clanga pomarina* | XC329332 | Łukasz Bednarz | *Phylloscopus trochilus* | XC913109 | Thomas Bergman |
| *Clanga pomarina* | XC563265 | Zbigniew Kajzer | *Pica pica* | XC862133 | Olivier Swift, Manuel Grosselet |
| *Clangula hyemalis* | XC106033 | Ryan P. O'Donnell | *Pica pica* | XC872510 | Paul Kelly |
| *Clangula hyemalis* | XC203477 | Andrew Spencer | *Pica pica* | XC879895 | Susanne Kuijpers |
| *Clangula hyemalis* | XC514046 | Timo Janhonen | *Pica pica* | XC897040 | Sławomir Karpicki |
| *Clangula hyemalis* | XC820344 | Johan Willner | *Picoides tridactylus* | XC715987 | Thomas Bergman |
| *Coccothraustes coccothraustes* | XC626792 | brickegickel | *Picoides tridactylus* | XC738886 | Thomas Bergman |
| *Coccothraustes coccothraustes* | XC707043 | Lars Edenius | *Picus canus* | XC878809 | András Schmidt |
| *Coccothraustes coccothraustes* | XC710720 | Török Tamás | *Picus canus* | XC882147 | Lars Edenius |
| *Coccothraustes coccothraustes* | XC712663 | Alan Dalton | *Picus viridis* | XC882026 | Jean Pierre Sciolla |
| *Colinus virginianus* | XC138625 | Paul Marvin | *Picus viridis* | XC927450 | Martin Billard |
| *Colinus virginianus* | XC567479 | Dare Šere | *Pinicola enucleator* | XC684582 | Lars Edenius |
| *Coloeus monedula* | XC856321 | Thierry Thomas | *Pinicola enucleator* | XC778268 | Lars Edenius |
| *Coloeus monedula* | XC904574 | Martin Billard | *Pinicola enucleator* | XC895559 | Maarten Sluijter |
| *Columba bollii* | XC270048 | Peter Boesman | *Pinicola enucleator* | XC895942 | Maarten Sluijter |
| *Columba bollii* | XC530191 | Mats Rellmar | *Platalea leucorodia* | XC596491 | Tristan Guillebot de Nerville |
| *Columba livia* | XC270097 | Timo Schnabel | *Platalea leucorodia* | XC768150 | Stanislas Wroza |
| *Columba livia* | XC92264 | Herman van der Meer | *Plectrophenax nivalis* | XC350489 | Tero Linjama |
| *Columba oenas* | XC738668 | Simon Elliott | *Plectrophenax nivalis* | XC379430 | Stein Ø. Nilsen |
| *Columba oenas* | XC747109 | Simon Elliott | *Plectrophenax nivalis* | XC425901 | Terje Kolaas |
| *Columba palumbus* | XC915274 | Susanne Kuijpers | *Plectrophenax nivalis* | XC425902 | Terje Kolaas |
| *Columba palumbus* | XC920599 | Lars Edenius | *Plegadis falcinellus* | XC530553 | Jordi Calvet |
| *Coracias garrulus* | XC661132 | Camille Monnet | *Plegadis falcinellus* | XC890702 | Bernard Bousquet |
| *Coracias garrulus* | XC770236 | Michel Veldt | *Pluvialis apricaria* | XC745635 | Lars Edenius |
| *Corvus corax* | XC791383 | Marcin Urbański | *Pluvialis apricaria* | XC789865 | Simon Elliott |
| *Corvus corax* | XC800465 | Nicolas Martinez | *Pluvialis apricaria* | XC818974 | Azens |
| *Corvus corax* | XC880900 | Lars Edenius | *Pluvialis apricaria* | XC831715 | W. Agster |
| *Corvus corax* | XC880911 | Lars Edenius | *Podiceps auritus* | XC375396 | Stein Ø. Nilsen |
| *Corvus corone* | XC815510 | Susanne Kuijpers | *Podiceps auritus* | XC715920 | Alan Dalton |

| | | |
|---|---|---|
| *Corvus corone* | XC856556 | Martin Billard |
| *Corvus corone* | XC878485 | Anthony Roux |
| *Corvus corone* | XC893406 | Pere Josa |
| *Corvus frugilegus* | XC770891 | Michel Veldt |
| *Corvus frugilegus* | XC780915 | Jorge Leitão |
| *Corvus frugilegus* | XC783382 | Stuart Fisher |
| *Corvus frugilegus* | XC883859 | Alain Malengreau |
| *Corvus splendens* | XC254340 | az dipu |
| *Corvus splendens* | XC322403 | Thomas Lüthi |
| *Corvus splendens* | XC357593 | Oscar Campbell |
| *Corvus splendens* | XC446886 | Audevard Aurélien |
| *Coturnix coturnix* | XC915429 | Lars Edenius |
| *Coturnix coturnix* | XC915737 | Joost van Bruggen |
| *Crex crex* | XC803640 | Frode Falkenberg |
| *Crex crex* | XC918715 | Dag Österlund |
| *Cuculus canorus* | XC710111 | Alain Malengreau |
| *Cuculus canorus* | XC917958 | Arjun Dutta |
| *Curruca balearica* | XC410968 | Jerome Fischer |
| *Curruca balearica* | XC411039 | Jerome Fischer |
| *Curruca balearica* | XC411067 | Jerome Fischer |
| *Curruca balearica* | XC411330 | Jerome Fischer |
| *Curruca cantillans* | XC807728 | Dare Šere |
| *Curruca cantillans* | XC907144 | Peter Carr |
| *Curruca communis* | XC898903 | Krzysztof Deoniziak |
| *Curruca communis* | XC898982 | Olivier Swift |
| *Curruca communis* | XC900942 | Martin Billard |
| *Curruca communis* | XC904596 | Niels Krabbe |
| *Curruca conspicillata* | XC470756 | Geoffrey Monchaux |
| *Curruca conspicillata* | XC537227 | Charlie Bodin |
| *Curruca conspicillata* | XC793044 | Stanislas Wroza |
| *Curruca conspicillata* | XC793047 | Stanislas Wroza |
| *Curruca conspicillata* | XC819009 | Paul Coiffard |
| *Curruca curruca* | XC800988 | Daniele Baroni |
| *Curruca curruca* | XC809662 | Dominique Guillerme |
| *Curruca curruca* | XC826101 | Esperanza Poveda |
| *Curruca curruca* | XC899224 | Bodo Sonnenburg |
| *Curruca hortensis* | XC801094 | João Tomás |
| *Curruca hortensis* | XC802317 | SonoNatura |
| *Curruca hortensis* | XC845162 | julien Birard |
| *Curruca iberiae* | XC722937 | Xavier Riera |
| *Curruca iberiae* | XC791461 | Maxence Fouillade |
| *Curruca melanocephala* | XC792298 | SonoNatura |
| *Curruca melanocephala* | XC900159 | David Pennington |
| *Curruca melanocephala* | XC917380 | Alain Verneau |
| *Curruca nisoria* | XC801193 | Bodo Sonnenburg |
| *Curruca nisoria* | XC807274 | Frank DD4WH |
| *Curruca nisoria* | XC816689 | Grzegorz Lorek |
| *Curruca nisoria* | XC845030 | Lukas Thiess |
| *Curruca nisoria* | XC905521 | Délfin Gonzalez |
| *Curruca sarda* | XC246980 | Jerome Fischer |
| *Curruca sarda* | XC370650 | Jerome Fischer |
| *Curruca sarda* | XC471853 | Stanislas Wroza |
| *Curruca sarda* | XC666367 | Marta Celej |
| *Curruca undata* | XC483891 | David Bissett |
| *Curruca undata* | XC530015 | Stanislas Wroza |
| *Curruca undata* | XC794772 | Jacobo Ramil Millarengo |
| *Curruca undata* | XC844566 | Martin Billard |
| *Cursorius cursor* | XC149578 | Frank Lambert |
| *Cursorius cursor* | XC164193 | Tero Linjama |
| *Cyanistes caeruleus* | XC893737 | Guillaume Wetzel |
| *Cyanistes caeruleus* | XC894018 | Tanguy Loïs |
| *Cyanistes caeruleus* | XC898570 | Jorge Leitão |
| *Cyanistes caeruleus* | XC898938 | Paul Kelly |
| *Cyanistes teneriffae* | XC367032 | Thomas Lüthi |
| *Cyanistes teneriffae* | XC458948 | Twan Mols |
| *Cyanistes teneriffae* | XC530210 | Mats Rellmar |
| *Cyanistes teneriffae* | XC796196 | Sidnei Dantas |
| *Cyanopica cooki* | XC341189 | José Carlos Sires |
| *Cyanopica cooki* | XC414693 | Stanislas Wroza |
| *Cyanopica cooki* | XC747540 | Jorge Leitão |
| *Cyanopica cooki* | XC747830 | Jorge Leitão |
| *Cygnus atratus* | XC297979 | Timo Schnabel |
| *Cygnus atratus* | XC674006 | Marc Anderson |
| *Cygnus cygnus* | XC791765 | Lars Edenius |
| *Cygnus cygnus* | XC795335 | Lars Edenius |
| *Cygnus olor* | XC764112 | Bodo Sonnenburg |
| *Cygnus olor* | XC861052 | Paul Kelly |
| *Cygnus olor* | XC864373 | Paul Kelly |
| *Cygnus olor* | XC879427 | Olivier Swift |
| *Delichon urbicum* | XC598648 | Simon Elliott |
| *Delichon urbicum* | XC726339 | Jorge Leitão |
| *Delichon urbicum* | XC792837 | João Tomás |
| *Delichon urbicum* | XC792855 | João Tomás |
| *Dendrocopos leucotos* | XC680685 | Jerome Fischer |
| *Dendrocopos leucotos* | XC689011 | Romuald Mikusek |
| *Dendrocopos leucotos* | XC713501 | Lars Edenius |
| *Dendrocopos leucotos* | XC719600 | Uku Paal |
| *Dendrocopos major* | XC622683 | Lars Edenius |
| *Dendrocopos major* | XC895779 | Lennart Jeppsson |
| *Dendrocopos major* | XC911171 | W. Agster |
| *Dendrocopos major* | XC926178 | Samir Hellal |
| *Dendrocopos syriacus* | XC480290 | Albert Lastukhin |
| *Dendrocopos syriacus* | XC528724 | Jarek Matusiak |
| *Dendrocopos syriacus* | XC667315 | András Schmidt |
| *Dendrocopos syriacus* | XC770414 | Michel Veldt |
| *Podiceps cristatus* | XC130588 | maudoc |
| *Podiceps cristatus* | XC773246 | Bernard Bousquet |
| *Podiceps cristatus* | XC790749 | Stanislas Wroza |
| *Podiceps cristatus* | XC801357 | Chèvremont Fabian |
| *Podiceps grisegena* | XC26934 | Patrik Åberg |
| *Podiceps grisegena* | XC804818 | Uku Paal |
| *Podiceps nigricollis* | XC26932 | Patrik Åberg |
| *Podiceps nigricollis* | XC590654 | Simon Elliott |
| *Poecile cinctus* | XC401785 | Karl-Birger Strann |
| *Poecile cinctus* | XC750044 | Lars Edenius |
| *Poecile cinctus* | XC752844 | Lars Edenius |
| *Poecile cinctus* | XC807539 | Thomas Bergman |
| *Poecile montanus* | XC818372 | Thomas Bergman |
| *Poecile montanus* | XC880573 | Bodo Sonnenburg |
| *Poecile montanus* | XC896685 | Zbigniew Kajzer |
| *Poecile montanus* | XC896794 | Lars Edenius |
| *Poecile palustris* | XC864410 | Bodo Sonnenburg |
| *Poecile palustris* | XC864410 | Bodo Sonnenburg |
| *Poecile palustris* | XC893807 | Azens |
| *Poecile palustris* | XC913905 | Elias A. Ryberg |
| *Poicephalus senegalus* | XC447381 | Étienne Leroy |
| *Poicephalus senegalus* | XC523303 | Joost van Bruggen |
| *Poicephalus senegalus* | XC719437 | Peter Boesman |
| *Poicephalus senegalus* | XC719438 | Peter Boesman |
| *Porphyrio porphyrio* | XC505960 | Jordi Calvet |
| *Porphyrio porphyrio* | XC546992 | Vincent Palomares |
| *Porphyrio porphyrio* | XC702417 | Stanislas Wroza |
| *Porphyrio porphyrio* | XC799681 | Pere Josa |
| *Porzana porzana* | XC900764 | Mats Olsson |
| *Porzana porzana* | XC917901 | Uku Paal |
| *Prunella collaris* | XC569165 | Stanislas Wroza |
| *Prunella collaris* | XC569172 | Stanislas Wroza |
| *Prunella collaris* | XC625477 | Marcel Gil Velasco |
| *Prunella collaris* | XC688398 | Stanislas Wroza |
| *Prunella modularis* | XC581070 | Arjun Dutta |
| *Prunella modularis* | XC881318 | Martin Billard |
| *Prunella modularis* | XC902691 | Paul Kelly |
| *Prunella modularis* | XC912899 | Romuald Mikusek |
| *Psittacara mitratus* | XC707076 | Richard E. Webster |
| *Psittacara mitratus* | XC931278 | Jorge Leitão |
| *Psittacula eupatria* | XC628895 | Stanislas Wroza |
| *Psittacula eupatria* | XC628898 | Stanislas Wroza |
| *Psittacula eupatria* | XC628899 | Stanislas Wroza |
| *Psittacula eupatria* | XC628900 | Stanislas Wroza |
| *Psittacula krameri* | XC694156 | Eduardo Realinho |
| *Psittacula krameri* | XC933414 | Jarek Matusiak |
| *Psittacula krameri* | XC933415 | Jarek Matusiak |
| *Psittacula krameri* | XC933415 | Jarek Matusiak |
| *Pterocles alchata* | XC142059 | Jordi Calvet |
| *Pterocles alchata* | XC924210 | Jordi Calvet |
| *Pterocles orientalis* | XC730394 | Stanislas Wroza |
| *Pterocles orientalis* | XC730396 | Stanislas Wroza |
| *Ptyonoprogne rupestris* | XC657489 | Camille Monnet |
| *Ptyonoprogne rupestris* | XC893409 | Pere Josa |
| *Pyrrhocorax graculus* | XC569238 | Stanislas Wroza |
| *Pyrrhocorax graculus* | XC769199 | Frank Lambert |
| *Pyrrhocorax graculus* | XC820207 | João Tomás |
| *Pyrrhocorax graculus* | XC898787 | Daniele Baroni |
| *Pyrrhocorax pyrrhocorax* | XC288230 | Antonio Xeira |
| *Pyrrhocorax pyrrhocorax* | XC584324 | Stanislas Wroza |
| *Pyrrhocorax pyrrhocorax* | XC596499 | Simon Elliott |
| *Pyrrhocorax pyrrhocorax* | XC735115 | Irish Wildlife Sounds |
| *Pyrrhula pyrrhula* | XC690409 | Lars Edenius |
| *Pyrrhula pyrrhula* | XC696696 | Lars Edenius |
| *Pyrrhula pyrrhula* | XC780464 | Lars Edenius |
| *Pyrrhula pyrrhula* | XC903130 | Corentin Rivière |
| *Rallus aquaticus* | XC689678 | Stanislas Wroza |
| *Rallus aquaticus* | XC789677 | Mats Olsson |
| *Rallus aquaticus* | XC859387 | Pere Josa |
| *Rallus aquaticus* | XC879688 | João Tomás |
| *Recurvirostra avosetta* | XC698221 | Olivier Swift, Hélène Cartaud |
| *Recurvirostra avosetta* | XC883880 | Pere Josa |
| *Regulus ignicapilla* | XC893407 | Pere Josa |
| *Regulus ignicapilla* | XC900900 | Ireneusz Oleksik |
| *Regulus ignicapilla* | XC902782 | Lars Edenius |
| *Regulus ignicapilla* | XC913156 | Romuald Mikusek |
| *Regulus regulus* | XC569378 | Stanislas Wroza |
| *Regulus regulus* | XC570588 | Paul Holt |
| *Regulus regulus* | XC831497 | Mats Rellmar |
| *Regulus regulus* | XC902433 | Elias A. Ryberg |
| *Remiz pendulinus* | XC770244 | Michel Veldt |
| *Remiz pendulinus* | XC828602 | Kuzey Cem Kulaçoğlu |
| *Remiz pendulinus* | XC884995 | João Tomás |
| *Remiz pendulinus* | XC902668 | Stephan Risch |
| *Riparia riparia* | XC659405 | Lars Edenius |
| *Riparia riparia* | XC666843 | Domagoj Tomičić |
| *Saxicola dacotiae* | XC153273 | Herman van Oosten |
| *Saxicola dacotiae* | XC346441 | Tero Linjama |
| *Saxicola rubetra* | XC797269 | Grzegorz Lorek |
| *Saxicola rubetra* | XC803189 | Lars Edenius |
| *Saxicola rubetra* | XC808861 | Lars Edenius |
| *Saxicola rubetra* | XC908625 | Martin Billard |
| *Saxicola rubicola* | XC716408 | Christian Kerihuel |
| *Saxicola rubicola* | XC789119 | João Tomás |
| *Saxicola rubicola* | XC799490 | Regina Eidner |

| | | |
|---|---|---|
| *Dendrocoptes medius* | XC859063 | Maarten Sluijter |
| *Dendrocoptes medius* | XC862406 | Martin Billard |
| *Dendrocoptes medius* | XC877498 | Guilherme Melo |
| *Dendrocoptes medius* | XC913132 | Romuald Mikusek |
| *Dryobates minor* | XC709565 | Alan Dalton |
| *Dryobates minor* | XC712215 | Lars Edenius |
| *Dryobates minor* | XC739164 | Uku Paal |
| *Dryobates minor* | XC742967 | Pontus Wennesjö |
| *Dryocopus martius* | XC839151 | Lars Edenius |
| *Dryocopus martius* | XC839284 | Lars Edenius |
| *Dryocopus martius* | XC902134 | Ulf Elman |
| *Dryocopus martius* | XC911632 | Jelle Scharringa |
| *Egretta garzetta* | XC774430 | Irish Wildlife Sounds |
| *Egretta garzetta* | XC790717 | Stanislas Wroza |
| *Egretta garzetta* | XC841128 | Ray Tsu |
| *Egretta garzetta* | XC856824 | Jorge Leitão |
| *Elanus caeruleus* | XC351398 | José Carlos Sires |
| *Elanus caeruleus* | XC352476 | José Carlos Sires |
| *Emberiza calandra* | XC835054 | Esperanza Poveda |
| *Emberiza calandra* | XC864606 | Frederic Lionel |
| *Emberiza calandra* | XC882055 | João Tomás |
| *Emberiza calandra* | XC899277 | David Pennington |
| *Emberiza cia* | XC467766 | Thijs Fijen |
| *Emberiza cia* | XC722554 | SonoNatura |
| *Emberiza cia* | XC724311 | brickegickel |
| *Emberiza cia* | XC865950 | Azens |
| *Emberiza cirlus* | XC799684 | Jacobo Ramil Millarengo |
| *Emberiza cirlus* | XC831373 | Francesco Sottile |
| *Emberiza cirlus* | XC882008 | Azens |
| *Emberiza cirlus* | XC892785 | Martin Billard |
| *Emberiza citrinella* | XC819013 | Grzegorz Lorek |
| *Emberiza citrinella* | XC826623 | Jack Berteau |
| *Emberiza citrinella* | XC879349 | Jarek Matusiak |
| *Emberiza citrinella* | XC895089 | Martin Billard |
| *Emberiza hortulana* | XC795809 | Stephan Börnecke |
| *Emberiza hortulana* | XC802483 | João Tomás |
| *Emberiza hortulana* | XC902403 | Grzegorz Lorek |
| *Emberiza hortulana* | XC903923 | Lars Edenius |
| *Emberiza melanocephala* | XC729335 | Leconte Michel |
| *Emberiza melanocephala* | XC741120 | Agris Celmins |
| *Emberiza melanocephala* | XC801378 | Stephan Risch |
| *Emberiza melanocephala* | XC813175 | Domagoj Tomičić |
| *Emberiza pusilla* | XC484445 | Fraser Simpson |
| *Emberiza pusilla* | XC571116 | Ulf Hansson |
| *Emberiza pusilla* | XC779437 | Erik Normark |
| *Emberiza pusilla* | XC809482 | Kasper Wieck |
| *Emberiza rustica* | XC655533 | Olavi Hinkkanen |
| *Emberiza rustica* | XC815431 | Lars Edenius |
| *Emberiza rustica* | XC900006 | Thomas Bergman |
| *Emberiza rustica* | XC904018 | Lars Edenius |
| *Emberiza schoeniclus* | XC790740 | Stanislas Wroza |
| *Emberiza schoeniclus* | XC791229 | Marcin Urbański |
| *Emberiza schoeniclus* | XC792702 | Stuart Fisher |
| *Emberiza schoeniclus* | XC796332 | Albert Lastukhin |
| *Eremophila alpestris* | XC485435 | Stanislas Wroza |
| *Eremophila alpestris* | XC571278 | Lars Edenius |
| *Erithacus rubecula* | XC865221 | Jean Pierre Sciolla |
| *Erithacus rubecula* | XC897347 | W. Agster |
| *Erithacus rubecula* | XC897396 | Jacobo Ramil Millarengo |
| *Erithacus rubecula* | XC906740 | Mats Rellmar |
| *Estrilda astrild* | XC640274 | Jacobo Ramil MIllarengo |
| *Estrilda astrild* | XC719273 | Nicolas Martinez |
| *Estrilda astrild* | XC748911 | Jacobo Ramil Millarengo |
| *Estrilda astrild* | XC750258 | Jacobo Ramil Millarengo |
| *Estrilda troglodytes* | XC164462 | Fernand Deroussen |
| *Estrilda troglodytes* | XC719131 | Peter Boesman |
| *Estrilda troglodytes* | XC719132 | Peter Boesman |
| *Estrilda troglodytes* | XC957262 | Julien Birard |
| *Euodice malabarica* | XC471526 | Stanislas Wroza |
| *Euodice malabarica* | XC838993 | Piqué Simon |
| *Falco biarmicus* | ML615715308 | Shane Sumasgutner |
| *Falco biarmicus* | ML89506 | Justin Schuetz |
| *Falco biarmicus* | XC101188 | James and David Bradley |
| *Falco biarmicus* | XC398048 | Peter Boesman |
| *Falco cherrug* | XC412094 | Frank Lambert |
| *Falco cherrug* | XC491256 | Peter Boesman |
| *Falco columbarius* | XC731075 | Lars Edenius |
| *Falco columbarius* | XC896663 | Tero Linjama |
| *Falco eleonorae* | XC854572 | Wim Jacobs |
| *Falco eleonorae* | XC893157 | Alain Malengreau |
| *Falco naumanni* | XC383877 | maudoc |
| *Falco naumanni* | XC579422 | Jordi Calvet |
| *Falco peregrinus* | XC748303 | Irish Wildlife Sounds |
| *Falco peregrinus* | XC896686 | Sven Normant |
| *Falco rusticolus* | XC189210 | Andrew Spencer |
| *Falco rusticolus* | XC709104 | Lars Edenius |
| *Falco rusticolus* | XC710854 | Dag Österlund |
| *Falco rusticolus* | XC713307 | Lars Edenius |
| *Falco subbuteo* | XC586296 | Hugo Wieleman |
| *Saxicola rubicola* | XC909121 | Mercier Christophe |
| *Scolopax rusticola* | XC397413 | Ruud van Beusekom |
| *Scolopax rusticola* | XC922073 | Teet Sirotkin |
| *Serinus canaria* | XC722081 | SonoNatura |
| *Serinus canaria* | XC722083 | SonoNatura |
| *Serinus canaria* | XC722084 | SonoNatura |
| *Serinus canaria* | XC831509 | Mats Rellmar |
| *Serinus serinus* | XC725616 | Uku Paal |
| *Serinus serinus* | XC812475 | Romuald Mikusek |
| *Serinus serinus* | XC875263 | João Tomás |
| *Serinus serinus* | XC900917 | Martin Billard |
| *Sinosuthora webbiana* | XC241076 | Michael Grunwell |
| *Sinosuthora webbiana* | XC464672 | Martin Sutherland |
| *Sinosuthora webbiana* | XC722418 | Common Jeff（麻杰夫） |
| *Sinosuthora webbiana* | XC889840 | Geoff Carey |
| *Sitta europaea* | XC897314 | Lennart Jeppsson |
| *Sitta europaea* | XC897670 | Jacobo Ramil Millarengo |
| *Sitta europaea* | XC927958 | Ulf Elman |
| *Sitta europaea* | XC938013 | Francesco Sottile |
| *Somateria mollissima* | XC383544 | Terje Kolaas |
| *Somateria mollissima* | XC549046 | Lars Edenius |
| *Spatula clypeata* | XC644500 | Jarek Matusiak |
| *Spatula clypeata* | XC863292 | João Tomás |
| *Spatula querquedula* | XC544441 | Hans Matheve |
| *Spatula querquedula* | XC650914 | Lars Edenius |
| *Spinus spinus* | XC594919 | Simon Elliott |
| *Spinus spinus* | XC793450 | Lars Edenius |
| *Spinus spinus* | XC799041 | Romuald Mikusek |
| *Spinus spinus* | XC871416 | Xavier Riera |
| *Streptopelia decaocto* | XC843476 | Joost van Bruggen |
| *Streptopelia decaocto* | XC881441 | Michel Veldt |
| *Streptopelia turtur* | XC862628 | Olivier Swift |
| *Streptopelia turtur* | XC902789 | Olivier Swift |
| *Strix aluco* | XC845803 | SonoNatura |
| *Strix aluco* | XC936220 | Martin Billard |
| *Strix nebulosa* | XC343167 | Tero Linjama |
| *Strix nebulosa* | XC655482 | Lars Edenius |
| *Strix uralensis* | XC705899 | Lars Edenius |
| *Strix uralensis* | XC750071 | Agris Celmins |
| *Strix uralensis* | XC797469 | Ulf Elman |
| *Strix uralensis* | XC884935 | Ulf Elman |
| *Sturnus unicolor* | XC642973 | Marcel Gil Velasco |
| *Sturnus unicolor* | XC665188 | Jorge Leitão |
| *Sturnus unicolor* | XC683425 | Jacobo Ramil MIllarengo |
| *Sturnus unicolor* | XC752912 | David Tattersley |
| *Sturnus vulgaris* | XC543800 | Thomas Bergman |
| *Sturnus vulgaris* | XC604977 | Simon Elliott |
| *Sturnus vulgaris* | XC652012 | Uku Paal |
| *Sturnus vulgaris* | XC712625 | Olivier Swift, Ludivine Delamare |
| *Surnia ulula* | XC793932 | Teet Sirotkin |
| *Surnia ulula* | XC862737 | Stein Ø. Nilsen |
| *Sylvia atricapilla* | XC858846 | Jacobo Ramil Millarengo |
| *Sylvia atricapilla* | XC880829 | Jean Pierre Sciolla |
| *Sylvia atricapilla* | XC904739 | Thomas Bergman |
| *Sylvia atricapilla* | XC905957 | Grzegorz Lorek |
| *Sylvia borin* | XC836965 | Jack Berteau |
| *Sylvia borin* | XC904341 | Martin Billard |
| *Sylvia borin* | XC906128 | Ulf Elman |
| *Sylvia borin* | XC906390 | Azens |
| *Syrmaticus reevesii* | XC360690 | Stanislas Wroza |
| *Syrmaticus reevesii* | XC360691 | Stanislas Wroza |
| *Tachybaptus ruficollis* | XC634413 | Peter Boesman |
| *Tachybaptus ruficollis* | XC720814 | Ulf Elman |
| *Tachymarptis melba* | XC544109 | Jordi Calvet |
| *Tachymarptis melba* | XC569487 | Beatrix Saadi-Varchmin |
| *Tadorna ferruginea* | XC683992 | Geoff Carey |
| *Tadorna ferruginea* | XC778623 | Valentin Monnoy |
| *Tadorna ferruginea* | XC835668 | Martin Billard |
| *Tadorna ferruginea* | XC872210 | Johannes Dag Mayer |
| *Tadorna tadorna* | XC735778 | Irish Wildlife Sounds |
| *Tadorna tadorna* | XC882474 | Jean Courtin |
| *Tadorna tadorna* | XC909997 | Uku Paal |
| *Tadorna tadorna* | XC936128 | Arjun Dutta |
| *Tarsiger cyanurus* | XC606234 | Niels Van Doninck |
| *Tarsiger cyanurus* | XC644039 | Agris Celmins |
| *Tarsiger cyanurus* | XC661672 | Lukas Thiess |
| *Tarsiger cyanurus* | XC912128 | Michel Watelet |
| *Tetrao urogallus* | XC473720 | Simon Ernst |
| *Tetrao urogallus* | XC611477 | Stanislas Wroza |
| *Tetrao urogallus* | XC639001 | Stanislas Wroza |
| *Tetrao urogallus* | XC641981 | Stanislas Wroza |
| *Tetrastes bonasia* | XC779196 | Lars Edenius |
| *Tetrastes bonasia* | XC814145 | Cedric Mroczko |
| *Tetrastes bonasia* | XC881064 | Lars Edenius |
| *Tetrastes bonasia* | XC904071 | Corentin Rivière |
| *Tetrax tetrax* | XC363911 | Stanislas Wroza |
| *Tetrax tetrax* | XC722329 | SonoNatura |
| *Thectocercus acuticaudatus* | XC189895 | Andrew Spencer |
| *Thectocercus acuticaudatus* | XC189896 | Andrew Spencer |

| | | | | | |
|---|---|---|---|---|---|
| *Falco subbuteo* | XC796706 | Lars Edenius | *Thectocercus acuticaudatus* | XC635097 | Dante Buzzetti |
| *Falco tinnunculus* | XC822020 | Dominique Guillerme | *Thectocercus acuticaudatus* | XC879279 | Dante Buzzetti |
| *Falco tinnunculus* | XC876577 | João Tomás | *Threskiornis aethiopicus* | XC278223 | Andrew Spencer |
| *Falco vespertinus* | XC770207 | Michel Veldt | *Threskiornis aethiopicus* | XC429511 | Frank Lambert |
| *Falco vespertinus* | XC813402 | Peter Boesman | *Threskiornis aethiopicus* | XC505935 | Lynette Rudman |
| *Ficedula albicollis* | XC653221 | brickegickel | *Threskiornis aethiopicus* | XC651807 | Nature sounds by Simply Birding |
| *Ficedula albicollis* | XC653222 | brickegickel | *Tichodroma muraria* | XC512330 | Thomas Armand |
| *Ficedula albicollis* | XC655679 | W. Agster | *Tichodroma muraria* | XC569274 | Stanislas Wroza |
| *Ficedula albicollis* | XC723932 | Ireneusz Oleksik | *Tichodroma muraria* | XC573620 | Sean Ronayne |
| *Ficedula hypoleuca* | XC802629 | bert de bruin | *Tichodroma muraria* | XC698260 | András Schmidt |
| *Ficedula hypoleuca* | XC803215 | Mats Rellmar | *Tringa erythropus* | XC564369 | Lars Edenius |
| *Ficedula hypoleuca* | XC806550 | Alan Dalton | *Tringa erythropus* | XC915323 | Thomas Bergman |
| *Ficedula hypoleuca* | XC908488 | Jacek Mucha | *Tringa glareola* | XC903314 | Lars Edenius |
| *Ficedula parva* | XC736819 | Lars Edenius | *Tringa glareola* | XC906765 | Francesco Sottile |
| *Ficedula parva* | XC743060 | Romuald Mikusek | *Tringa nebularia* | XC900331 | Uku Paal |
| *Ficedula parva* | XC804988 | Mikael Litsgård | *Tringa nebularia* | XC917906 | Esperanza Poveda |
| *Ficedula parva* | XC807830 | Lars Edenius | *Tringa ochropus* | XC803363 | Lars Edenius |
| *Fringilla coelebs* | XC897397 | Jacobo Ramil Millarengo | *Tringa ochropus* | XC808190 | Lars Edenius |
| *Fringilla coelebs* | XC908632 | Martin Billard | *Tringa totanus* | XC884810 | Olivier Swift |
| *Fringilla coelebs* | XC908812 | Olivier Swift | *Tringa totanus* | XC890808 | Christian Bøggild |
| *Fringilla coelebs* | XC909647 | Carlos Pereira | *Tringa totanus* | XC898939 | Paul Kelly |
| *Fringilla montifringilla* | XC653538 | Agris Celmins | *Tringa totanus* | XC906104 | Thomas Bergman |
| *Fringilla montifringilla* | XC675120 | Albert Lastukhin | *Troglodytes troglodytes* | XC916801 | Olivier Swift |
| *Fringilla montifringilla* | XC731505 | Marc Anderson | *Troglodytes troglodytes* | XC928304 | Jorge Leitão |
| *Fringilla montifringilla* | XC861643 | Nicolas Martinez | *Turdus iliacus* | XC808871 | Lars Edenius |
| *Fulica atra* | XC715239 | Susanne Kuijpers | *Turdus iliacus* | XC881694 | Eric Roualet |
| *Fulica atra* | XC881746 | Paul Kelly | *Turdus iliacus* | XC901039 | Kai Rantala |
| *Fulica atra* | XC893840 | Susanne Kuijpers | *Turdus iliacus* | XC903924 | Lars Edenius |
| *Fulica atra* | XC902418 | Jack Berteau | *Turdus merula* | XC815531 | Agris Celmins |
| *Fulica cristata* | XC306654 | Hans Matheve | *Turdus merula* | XC900559 | Lennart Jeppsson |
| *Fulica cristata* | XC431996 | Joost van Bruggen | *Turdus merula* | XC906491 | Olivier Swift |
| *Fulica cristata* | XC860421 | Dries Van de Loock | *Turdus merula* | XC907818 | Olivier Swift |
| *Fulica cristata* | XC908546 | Nelson Conceição | *Turdus philomelos* | XC831495 | Agris Celmins |
| *Galerida cristata* | XC832503 | Stein Ø. Nilsen | *Turdus philomelos* | XC879352 | Alain Malengreau |
| *Galerida cristata* | XC839414 | Corentin Rivière | *Turdus philomelos* | XC897346 | W. Agster |
| *Galerida cristata* | XC864276 | SonoNatura | *Turdus philomelos* | XC899778 | Jorge Leitão |
| *Galerida cristata* | XC864607 | Frederic Lionel | *Turdus pilaris* | XC635213 | Stanislas Wroza |
| *Galerida theklae* | XC463869 | Jordi Calvet | *Turdus pilaris* | XC686589 | Niels Van Doninck |
| *Galerida theklae* | XC730352 | Stanislas Wroza | *Turdus pilaris* | XC727604 | Lars Edenius |
| *Galerida theklae* | XC779770 | Luis Gracia | *Turdus pilaris* | XC834617 | Grégoire Chauvot |
| *Galerida theklae* | XC900368 | Christian Kerihuel | *Turdus torquatus* | XC593279 | Simon Elliott |
| *Gallinago gallinago* | XC633300 | Krzysztof Deoniziak | *Turdus torquatus* | XC721589 | Loan Delpit |
| *Gallinago gallinago* | XC729310 | Lars Edenius | *Turdus torquatus* | XC724232 | Romuald Mikusek |
| *Gallinago media* | XC201361 | Stein Ø. Nilsen | *Turdus torquatus* | XC837014 | Petter Westberg |
| *Gallinago media* | XC325087 | Terje Kolaas | *Turdus viscivorus* | XC718033 | Manuel Grosselet |
| *Gallinula chloropus* | XC894656 | Lars Mogensen | *Turdus viscivorus* | XC788562 | Jacobo Ramil Millarengo |
| *Gallinula chloropus* | XC895059 | Martin Billard | *Turdus viscivorus* | XC807826 | Lars Edenius |
| *Gallinula chloropus* | XC902620 | Lars Mogensen | *Turdus viscivorus* | XC879652 | Christian Vemmelund Helligsø |
| *Gallinula chloropus* | XC905071 | Vandousselaere Patrick | *Tyto alba* | XC948997 | SonoNatura |
| *Garrulus glandarius* | XC856324 | Thierry Thomas | *Tyto alba* | XC952015 | SonoNatura |
| *Garrulus glandarius* | XC881809 | Esperanza Poveda | *Tyto alba* | XC952969 | Friedrich Richard |
| *Garrulus glandarius* | XC884360 | Thomas Bergman | *Tyto alba* | XC962216 | Hans Matheve |
| *Garrulus glandarius* | XC902468 | Jack Berteau | *Upupa epops* | XC897074 | SonoNatura |
| *Geronticus eremita* | XC361510 | José Carlos Sires y Eloisa Matheu | *Upupa epops* | XC917761 | Regina Eidner |
| *Geronticus eremita* | XC361523 | José Carlos Sires y Eloisa Matheu | *Vanellus vanellus* | XC638538 | Thomas Bergman |
| *Glareola pratincola* | XC379455 | Stanislas Wroza | *Vanellus vanellus* | XC880717 | Olivier Swift |
| *Glareola pratincola* | XC758522 | Dries Van de Loock | *Vanellus vanellus* | XC881982 | Olivier Swift |
| *Glaucidium passerinum* | XC883372 | Teet Sirotkin | *Vanellus vanellus* | XC898816 | Thomas Bergman |
| *Glaucidium passerinum* | XC902777 | David Darrell-Lambert | *Zapornia parva* | XC732543 | Tero Linjama |
| *Grus grus* | XC723192 | Lars Edenius | *Zapornia parva* | XC900252 | Krzysztof Deoniziak |
| *Grus grus* | XC836029 | Hannu Varkki | *Zapornia pusilla* | XC656129 | Yannick Jacob |
| *Gulosus aristotelis* | XC588002 | Simon Elliott | *Zapornia pusilla* | XC745918 | Marcin Sołowiej |

Supplementary Table 15. Acoustic parameters used in a principal component analysis in order to calculate vocalisation dissimilarity, and their transformations.

| Acoustic Parameters | Name | Transformation |
|---|---|---|
| Duration | duration | log(x + 0.001) |
| Mean frequency | meanfreq | Untransformed |
| Standard deviation of frequency | sd | log(x + 0.001) |
| Median frequency | freq.median | Untransformed |
| First quartile frequency | freq.Q25 | Untransformed |
| Third quartile frequency | freq.Q75 | sqrt |
| Frequency interquartile range | freq.IQR | sqrt |
| Median time (time at which the signal is divided in two time intervals of equal energy) | time.median | log(x + 0.001) |
| First quartile time | time.Q25 | log(x + 0.001) |
| Third quartile time | time.Q75 | log(x + 0.001) |
| Time interquartile range | time.IQR | log(x + 0.001) |
| Spectral skewness | skew | log(x + 0.001) |
| Spectral kurtosis (peakedness) | kurt | log(x + 0.001) |
| Spectral entropy (energy distribution of the frequency spectrum) | sp.ent | Untransformed |
| Time entropy (energy distribution on the time envelope) | time.ent | Untransformed |
| Product of time and spectral entropy | entropy | Untransformed |
| Spectral flatness | sfm | log(x + 0.001) |
| Mean peak frequency | meanpeakf | Untransformed |
| Average dominant frequency | meandom | Untransformed |
| Maximum of dominant frequency | maxdom | Untransformed |
| Minimum dominant frequency | mindom | Untransformed |
| Range of dominant frequency measured across the acoustic signal | dfrange | Untransformed |
| Modulation index | modindx | log(x + 0.001) |
| Dominant frequency measurement at the start of the signal | startdom | sqrt |
| Dominant frequency measurement at the end of the signal | enddom | sqrt |
| Slope of the change in dominant frequency through time | dfslope | Untransformed |
| Number of notes | note.count | log(x + 0.001) |
| Note rate | note.rate | log(x + 0.001) |
| Length of longest note | longest.note | log(x + 0.001) |
| Percentage of the song consisting of notes | note.perc | Untransformed |
| Average note length | ave.note.length | log(x + 0.001) |
| Total note duration | total.note.dur | log(x + 0.001) |
| Average pause length | ave.pause.length | log(x + 0.001) |
| Length of longest pause | longest.pause | log(x + 0.001) |

Supplementary Table 16. Studies informing the CLOOTL phylogenetic tree utilised and the proportion of nodes in the phylogeny that are supported by each study.

| References | contribution |
|---|---|
| Jetz, W., G. H. Thomas, J. B. Joy, K. Hartmann, A. O. Mooers. 2012. The global diversity of birds in space and time. Nature 491 (7424): 444-448 | 57.7 |
| Burleigh, J. Gordon, Rebecca T. Kimball, Edward L. Braun. 2015. Building the avian tree of life using a large-scale, sparse supermatrix. Molecular Phylogenetics and Evolution 84: 53-63 | 52.8 |
| Kimball, R. T., Oliveros, C. H., Wang, N., White, N. D., Barker, F. K., Field, D. J., ... Braun, E. L. (2019). A Phylogenomic Supertree of Birds. Diversity, 11(7), 109. doi:10.3390/d11070109 | 11.5 |
| Barker, F. Keith, Kevin J. Burns, John Klicka, Scott M. Lanyon, Irby J. Lovette. 2015. New insights into New World biogeography: An integrated view from the phylogeny of blackbirds, cardinals, sparrows, tanagers, warblers, and allies. The Auk 132 (2): 333-348. | 11.3 |
| Selvatti, Alexandre Pedro, Luiz Pedreira Gonzaga, Claudia Augusta de Moraes Russo. 2015. A Paleogene origin for crown passerines and the diversification of the Oscines in the New World. Molecular Phylogenetics and Evolution 88: 1-15. | 10 |
| Stiller, J., Feng, S., Chowdhury, AA. et al. Complexity of avian evolution revealed by family-level genomes. Nature (2024). https://doi.org/10.1038/s41586-024-07323-1 | 9.2 |
| Hooper, Daniel M., Trevor D. Price. 2017. Chromosomal inversion differences correlate with range overlap in passerine birds. Nature Ecology & Evolution 1 (10): 1526-1534 | 7.4 |
| Price, Trevor D., Daniel M. Hooper, Caitlyn D. Buchanan, Ulf S. Johansson, D. Thomas Tietze, Per Alström, Urban Olsson, Mousumi Ghosh-Harihar, Farah Ishtiaq, Sandeep K. Gupta, Jochen Martens, Bettina Harr, Pratap Singh, Dhananjai Mohan. 2014. Niche filling slows the diversification of Himalayan songbirds. Nature 509: 222-225. | 7.2 |
| Päckert, M., Favre, A., Schnitzler, J., Martens, J., Sun, Y., Tietze, D. T., Hailer, F., Michalak, I., & Strutzenberger, P. (2020). “Into and Out of” the Qinghai-Tibet Plateau and the Himalayas: Centers of origin and diversification across five clades of Eurasian montane and alpine passerine birds. Ecology and Evolution, 10(17), 9283–9300. Portico. https://doi.org/10.1002/ece3.6615 | 6.3 |
| Černý, D., & Natale, R. (2022). Comprehensive taxon sampling and vetted fossils help clarify the time tree of shorebirds (Aves, Charadriiformes). Molecular Phylogenetics and Evolution, 177, 107620. https://doi.org/10.1016/j.ympev.2022.107620 | 6 |
| Zhao, M., Oswald, J. A., Allen, J. M., Owens, H. L., Hosner, P. A., Guralnick, R. P., Braun, E. L., & Kimball, R. T. (2025). A phylogenomic tree of wood-warblers (Aves: Parulidae): Dealing with good, bad, and ugly samples. Molecular Phylogenetics and Evolution, 202, 108235. https://doi.org/10.1016/j.ympev.2024.108235 | 5.7 |
| McCullough, J. M., Oliveros, C. H., Benz, B. W., Zenil-Ferguson, R., Cracraft, J., Moyle, R. G., & Andersen, M. J. (2022). Wallacean and Melanesian Islands Promote Higher Rates of Diversification within the Global Passerine Radiation Corvides. Systematic Biology. https://doi.org/10.1093/sysbio/syac044 | 5.5 |
| Catanach, T. A., Halley, M. R., & Pirro, S. (2024). Enigmas no longer: using ultraconserved elements to place several unusual hawk taxa and address the non-monophyly of the genusAccipiter(Accipitriformes: Accipitridae). Biological Journal of the Linnean Society. https://doi.org/10.1093/biolinnean/blae028 | 5.1 |
| Claramunt, Santiago, Joel Cracraft. 2015. A new time tree reveals Earth historys imprint on the evolution of modern birds. Science Advances 1 (11): e1501005-e1501005 | 4.9 |
| Prum, Richard O., Jacob S. Berv, Alex Dornburg, Daniel J. Field, Jeffrey P. Townsend, Emily Moriarty Lemmon, Alan R. Lemmon. 2015. A comprehensive phylogeny of birds (Aves) using targeted next-generation DNA sequencing. Nature 526, (7574): 569-573 | 4.8 |
| Reddy, Sushma, Rebecca T. Kimball, Akanksha Pandey, Peter A. Hosner, Michael J. Braun, Shannon J. Hackett, Kin-Lan Han, John Harshman, Christopher J. Huddleston, Sarah Kingston, Ben D. Marks, Kathleen J. Miglia, William S. Moore, Frederick H. Sheldon, Christopher C. Witt, Tamaki Yuri, Edward L. Braun. 2017. Why do phylogenomic data sets yield conflicting trees? Data type influences the avian tree of life more than taxon sampling. Systematic Biology | 4.6 |
| Oliveros, C. H., Field, D. J., Ksepka, D. T., Barker, F. K., Aleixo, A., Andersen, M. J., ... Faircloth, B. C. (2019). Earth history and the passerine superradiation. Proceedings of the National Academy of Sciences, 116(16), 7916–7925. doi:10.1073/pnas.1813206116 | 4.6 |
| Gonzalez, J., H. Düttmann, M. Wink. 2009. Phylogenetic relationships based on two mitochondrial genes and hybridization patterns in Anatidae. Journal of Zoology 279 (3): 310-318. | 4.4 |
| Jønsson, Knud Andreas, Pierre-Henri Fabre, Jonathan D. Kennedy, Ben G. Holt, Michael K. Borregaard, Carsten Rahbek, Jon Fjeldså. 2016. A supermatrix phylogeny of corvoid passerine birds (Aves: Corvides). Molecular Phylogenetics and Evolution 94: 87-94. | 4.3 |
| Feng, S., Stiller, J., Deng, Y., Armstrong, J., Fang, Q., Reeve, A. H., Xie, D., Chen, G., Guo, C., Faircloth, B. C., Petersen, B., Wang, Z., Zhou, Q., Diekhans, M., Chen, W., Andreu-Sánchez, S., Margaryan, A., Howard, J. T., Parent, C., ... Zhang, G. (2020). Dense sampling of bird diversity increases power of comparative genomics. Nature, 587(7833), 252–257. https://doi.org/10.1038/s41586-020-2873-9 | 4.2 |
| Harvey, M. G., Bravo, G. A., Claramunt, S., Cuervo, A. M., Derryberry, G. E., Battilana, J., ... Derryberry, E. P. (2020). The evolution of a tropical biodiversity hotspot. Science, 370(6522), 1343–1348. doi:10.1126/science.aaz6970 | 4.1 |

Fulton, T. L., B. Letts, B. Shapiro. 2012. Multiple losses of flight and recent speciation in steamer ducks. Proceedings of the Royal Society B: Biological Sciences 279 (1737): 2339-2346. 3.8

Shakya, S. B., Fuchs, J., Pons, J.-M., & Sheldon, F. H. (2017). Tapping the woodpecker tree for evolutionary insight. Molecular Phylogenetics and Evolution, 116, 182–191. doi:10.1016/j.ympev.2017.09.005 3.6

Nagy, J., & Tökölyi, J. (2014). Phylogeny, Historical Biogeography and the Evolution of Migration in Accipitrid Birds of Prey (Aves: Accipitriformes). Ornis Hungarica, 22(1), 15–35. https://doi.org/10.2478/orhu-2014-0008 3.5

Stein, R. Will, Joseph W. Brown, Arne Ø. Mooers. 2015. A molecular genetic time scale demonstrates Cretaceous origins and multiple diversification rate shifts within the order Galliformes (Aves). Molecular Phylogenetics and Evolution 92: 155-164 3.2

Lovette, Irby J., Jorge L. Pérez-Emán, John P. Sullivan, Richard C. Banks, Isabella Fiorentino, Sergio Córdoba-Córdoba, María Echeverry-Galvis, F. Keith Barker, Kevin J. Burns, John Klicka, Scott M. Lanyon, Eldredge Bermingham. 2010. A comprehensive multilocus phylogeny for the wood-warblers and a revised classification of the Parulidae (Aves). Molecular Phylogenetics and Evolution 57 (2): 753-770. 3.2

Wu, S., Rheindt, F. E., Zhang, J., Wang, J., Zhang, L., Quan, C., Li, Z., Wang, M., Wu, F., Qu, Y., Edwards, S. V., Zhou, Z., & Liu, L. (2024). Genomes, fossils, and the concurrent rise of modern birds and flowering plants in the Late Cretaceous. Proceedings of the National Academy of Sciences, 121(8). https://doi.org/10.1073/pnas.2319696121 3.1

Klicka, John, F. Keith Barker, Kevin J. Burns, Scott M. Lanyon, Irby J. Lovette, Jaime A. Chaves, Robert W. Bryson. 2014. A comprehensive multilocus assessment of sparrow (Aves: Passerellidae) relationships. Molecular Phylogenetics and Evolution 77: 177–182. 3.1

Chen, D., Hosner, P. A., Dittmann, D. L., O'Neill, J. P., Birks, S. M., Braun, E. L., & Kimball, R. T. (2021). Divergence time estimation of Galliformes based on the best gene shopping scheme of ultraconserved elements. BMC Ecology and Evolution, 21(1). https://doi.org/10.1186/s12862-021-01935-1 3

Dufort, Matthew J. 2015. An augmented supermatrix phylogeny of the avian family Picidae reveals uncertainty deep in the family tree. Molecular Phylogenetics and Evolution 3

Alstrom, P., D. M. Hooper, Y. Liu, U. Olsson, D. Mohan, M. Gelang, H. Le Manh, J. Zhao, F. Lei, T. D. Price. 2014. Discovery of a relict lineage and monotypic family of passerine birds. Biology Letters 10 (3): 20131067-20131067. 2.8

Zuccon, Dario, Robert Prŷs-Jones, Pamela C. Rasmussen, and Per G.P. Ericson. 2012. The phylogenetic relationships and generic limits of finches (Fringillidae). Molecular Phylogenetics and Evolution 62 (2): 581-596. 2.8

Barker, F. K., K. J. Burns, J. Klicka, S. M. Lanyon, I. J. Lovette. 2013. Going to extremes: contrasting rates of diversification in a recent radiation of New World passerine birds. Systematic Biology 62 (2): 298-320. 2.6

Kimball, Rebecca T., Colette M. St. Mary, Edward L. Braun. 2011. A macroevolutionary perspective on multiple sexual traits in the Phasianidae (Galliformes). International Journal of Evolutionary Biology 2011: 1-16. 2.6

Gibson, Rosemary, Allan Baker. 2012. Multiple gene sequences resolve phylogenetic relationships in the shorebird suborder Scolopaci (Aves: Charadriiformes). Molecular Phylogenetics and Evolution 64 (1): 66-72 2.5

Scofield, R. Paul, Kieren J. Mitchell, Jamie R. Wood, Vanesa L. De Pietri, Scott Jarvie, Bastien Llamas, Alan Cooper. 2016. The Origin and Phylogenetic Relationships of the New Zealand Ravens. Molecular Phylogenetics and Evolution 106: 136–143 2.5

Cicero, C., Mason, N. A., Benedict, L., & Rising, J. D. (2020). Behavioral, morphological, and ecological trait evolution in two clades of New World Sparrows (AimophilaandPeucaea, Passerellidae). PeerJ, 8, e9249. Portico. https://doi.org/10.7717/peerj.9249 2.3

Zhao, M., Gordon Burleigh, J., Olsson, U., Alström, P., & Kimball, R. T. (2023). A near-complete and time-calibrated phylogeny of the Old World flycatchers, robins and chats (Aves, Muscicapidae). Molecular Phylogenetics and Evolution, 178, 107646. https://doi.org/10.1016/j.ympev.2022.107646 2.3

Imfeld, T. S., Barker, F. K., Vázquez-Miranda, H., Chaves, J. A., Escalante, P., Spellman, G. M., & Klicka, J. (2024). Diversification and dispersal in the Americas revealed by new phylogenies of the wrens and allies (Passeriformes: Certhioidea). Ornithology, 141(2). https://doi.org/10.1093/ornithology/ukae007 2.2

Powell, Alexis F.L.A., F. Keith Barker, Scott M. Lanyon, Kevin J. Burns, John Klicka, Irby J. Lovette. 2014. A comprehensive species-level molecular phylogeny of the New World blackbirds (Icteridae). Molecular Phylogenetics and Evolution 71: 94-112. 2.2

Barker, F.K. 2014. Mitogenomic data resolve basal relationships among passeriform and passeridan birds. Molecular Phylogenetics and Evolution 79: 313-324. 2.1

Hosner, Peter A. Brant C. Faircloth, Travis C. Glenn, Edward L. Braun, Rebecca T. Kimball. 2015. Avoiding missing data biases in phylogenomic inference: an empirical study in the landfowl (Aves: Galliformes). Molecular Biology and Evolution 33 (4): 1110-1125 2

Hugall, Andrew F., Devi Stuart-Fox. 2012. Accelerated speciation in colour-polymorphic birds. Nature 485 (7400): 631-634. 2

Cai, T., Cibois, A., Alström, P., Moyle, R. G., Kennedy, J. D., Shao, S., Zhang, R., Irestedt, M., Ericson, P. G. P., Gelang, M., Qu, Y., Lei, F., & Fjeldså, J. (2019). Near-complete phylogeny and taxonomic revision of the world's babblers (Aves: Passeriformes). Molecular Phylogenetics and Evolution, 130, 346–356. https://doi.org/10.1016/j.ympev.2018.10.010 2

Wang, Ning, Rebecca T. Kimball, Edward L. Braun, Bin Liang, Zhengwang Zhang. 2016. Ancestral range reconstruction of Galliformes: the effects of topology and taxon sampling. Journal of Biogeography 2

Moyle, Robert G., Carl H. Oliveros, Michael J. Andersen, Peter A. Hosner, Brett W. Benz, Joseph D. Manthey, Scott L. Travers, Rafe M. Brown, Brant C. Faircloth. 2016. Tectonic collision and uplift of Wallacea triggered the global songbird radiation. Nature Communications 7: 12709 2

Simone Treplin, Romy Siegert, Christoph Bleidorn, Hazell Shokellu Thompson, Roger Fotso, Ralph Tiedemann, 2008, 'Molecular phylogeny of songbirds (Aves: Passeriformes) and the relative utility of common nuclear marker loci', Cladistics, vol. 24, no. 3, pp. 328-349 2

Salter, J. F., Oliveros, C. H., Hosner, P. A., Manthey, J. D., Robbins, M. B., Moyle, R. G., Brumfield, R. T., & Faircloth, B. C. (2020). Extensive paraphyly in the typical owl family (Strigidae). The Auk, 137(1). https://doi.org/10.1093/auk/ukz070 1.8

Jarvis, E. D., S. Mirarab, A. J. Aberer, B. Li, P. Houde, C. Li, S. Y. W. Ho, B. C. Faircloth, B. Nabholz, J. T. Howard, A. Suh, C. C. Weber, R. R. da Fonseca, J. Li, F. Zhang, H. Li, L. Zhou, N. Narula, L. Liu, G. Ganapathy, B. Boussau, M. S. Bayzid, V. Zavidovych, S. Subramanian, T. Gabaldon, S. Capella-Gutierrez, J. Huerta-Cepas, B. Rekepalli, K. Munch, M. Schierup, B. Lindow, W. C. Warren, D. Ray, R. E. Green, M. W. Bruford, X. Zhan, A. Dixon, S. Li, N. Li, Y. Huang, E. P. Derryberry, M. F. Bertelsen, F. H. Sheldon, R. T. Brumfield, C. V. Mello, P. V. Lovell, M. Wirthlin, M. P. C. Schneider, F. Prosdocimi, J. A. Samaniego, A. M. V. Velazquez, A. Alfaro-Nunez, P. F. Campos, B. Petersen, T. Sicheritz-Ponten, A. Pas, T. Bailey, P. Scofield, M. Bunce, D. M. Lambert, Q. Zhou, P. Perelman, A. C. Driskell, B. Shapiro, Z. Xiong, Y. Zeng, S. Liu, Z. Li, B. Liu, K. Wu, J. Xiao, X. Yinqi, Q. Zheng, Y. Zhang, H. Yang, J. Wang, L. Smeds, F. E. Rheindt, M. Braun, J. Fjeldsa, L. Orlando, F. K. Barker, K. A. Jonsson, W. Johnson, K.-P. Koepfli, S. O'Brien, D. Haussler, O. A. Ryder, C. Rahbek, E. Willerslev, G. R. Graves, T. C. Glenn, J. McCormack, D. Burt, H. Ellegren, P. Alstrom, S. V. Edwards, A. Stamatakis, D. P. Mindell, J. Cracraft, E. L. Braun, T. Warnow, W. Jun, M. T. P. Gilbert, G. Zhang. 2014. Whole-genome analyses resolve early branches in the tree of life of modern birds. Science 346 (6215): 1320-1331. 1.7

Hruska, J. P., Holmes, J., Oliveros, C., Shakya, S., Lavretsky, P., McCracken, K. G., Sheldon, F. H., & Moyle, R. G. (2023). Ultraconserved elements resolve the phylogeny and corroborate patterns of molecular rate variation in herons (Aves: Ardeidae). Ornithology. https://doi.org/10.1093/ornithology/ukad005 1.7

Wood, Jamie R., Kieren J. Mitchell, R. Paul Scofield, Vanesa L. De Pietri, Nicolas J. Rawlence, Alan Cooper. 2016. Phylogenetic relationships and terrestrial adaptations of the extinct laughing owl, Sceloglaux albifacies (Aves: Strigidae). Zoological Journal of the Linnean Society 1.7

McGuire, Jimmy A., Christopher C. Witt, J.V. Remsen, Ammon Corl, Daniel L. Rabosky, Douglas L. Altshuler, Robert Dudley. 2014. Molecular phylogenetics and the diversification of hummingbirds. Current Biology 24 (8): 910-916. 1.6

Fuchs, J., Johnson, J. A., & Mindell, D. P. (2015). Rapid diversification of falcons (Aves: Falconidae) due to expansion of open habitats in the Late Miocene. Molecular Phylogenetics and Evolution, 82, 166–182. https://doi.org/10.1016/j.ympev.2014.08.010 1.5

Smith, B. T., Merwin, J., Provost, K. L., Thom, G., Brumfield, R. T., Ferreira, M., Mauck, W. M., Moyle, R. G., Wright, T. F., & Joseph, L. (2022). Phylogenomic Analysis of the Parrots of the World Distinguishes Artifactual from Biological Sources of Gene Tree Discordance. Systematic Biology. https://doi.org/10.1093/sysbio/syac055 1.5

Claramunt, S., Derryberry E.P., Brumfield R., and Remsen J. 2012. Ecological opportunity and diversification in a continental radiation of birds: climbing adaptations and cladogenesis in the Furnariidae. American Naturalist 179 (5): 649-666. 1.5

Barker, F. Keith, Alice Cibois, Peter Schikler, Julie Feinstein, Joel Cracraft. 2004. Phylogeny and diversification of the largest avian radiation. Proceedings of the National Academy of Sciences of the United States of America 101 (30): 11040-11045. 1.5

Lerner, H., M. Klaver, D. Mindell. 2008. Molecular phylogenetics of the Buteonine birds of prey (Accipitridae). The Auk 125 (2): 304-315. 1.4

Cai, T., Wu, G., Sun, L., Zhang, Y., Peng, Z., Guo, Y., Liu, X., Pan, T., Chang, J., Sun, Z., & Zhang, B. (2021). Biogeography and diversification of Old World buntings (Aves: Emberizidae): radiation in open habitats. Journal of Avian Biology, 52(6). Portico. https://doi.org/10.1111/jav.02672 1.2

Bryson, R. W., Faircloth, B. C., Tsai, W. L. E., McCormack, J. E., & Klicka, J. (2016). Target enrichment of thousands of ultraconserved elements sheds new light on early relationships within New World sparrows (Aves: Passerellidae). The Auk, 133(3), 451–458. https://doi.org/10.1642/auk-16-26.1 1.2

Kirchman, J. J., McInerney, N. R., Giarla, T. C., Olson, S. L., Slikas, E., & Fleischer, R. C. (2021). Corrigendum to: Phylogeny based on ultra-conserved elements clarifies the evolution of rails and allies (Ralloidea) and is the basis for a revised classification. Ornithology, 139(1). https://doi.org/10.1093/ornithology/ukab065 1.2

Schield, D. R., Brown, C. E., Shakya, S. B., Calabrese, G. M., Safran, R. J., & Sheldon, F. H. (2024). Phylogeny and historical biogeography of the swallow family (Hirundinidae) inferred from comparisons of thousands of UCE loci. Molecular Phylogenetics and Evolution, 197, 108111. https://doi.org/10.1016/j.ympev.2024.108111 1.2

Odeen, A., Hastad O., & Alstrom P. 2011. Evolution of ultraviolet vision in the largest avian radiation - the passerines. BMC Evolutionary Biology 11 (313): 1-8. 1.2

Johansson, Ulf S., Jan Ekman, Rauri C.K. Bowie, Peter Halvarsson, Jan I. Ohlson, Trevor D. Price, Per G.P. Ericson. 2013. A complete multilocus species phylogeny of the tits and chickadees (Aves: Paridae). Molecular Phylogenetics and Evolution 69 (3): 852-860. 1.2

Miller et al. 1.2

Martim Stervander, Peter G. Ryan, Martim Melo, Bengt Hansson, 2019, 'The origin of the world's smallest flightless bird, the Inaccessible Island Rail Atlantisia rogersi (Aves: Rallidae)', Molecular Phylogenetics and Evolution, vol. 130, pp. 92-98 1.1

Oswald, J. A., Boyd, B. M., Szewczak, A. R., LeFebvre, M. J., Stucky, B. J., Guralnick, R. P., Johnson, K. P., Allen, J. M., & Steadman, D. W. (2025). Genomic data reveal that the Cuban blue-headed quail-dove (
Starnoenas cyanocephala
) is a biogeographic relict. Biology Letters, 21(1). https://doi.org/10.1098/rsbl.2024.0464 1.1

Ulf S. Johansson, Jon Fjeldså, Rauri C.K. Bowie, 2008, 'Phylogenetic relationships within Passerida (Aves: Passeriformes): A review and a new molecular phylogeny based on three nuclear intron markers', Molecular Phylogenetics and Evolution, vol. 48, no. 3, pp. 858-876 1.1

Donne-Goussé, Carole, Vincent Laudet, Catherine Hänni. 2002. A molecular phylogeny of anseriformes based on mitochondrial DNA analysis. Molecular Phylogenetics and Evolution 23 (3): 339-356 1

Sheldon, Frederick H., Linda A. Whittingham, Robert G. Moyle, Beth Slikas, David W. Winkler. 2005. Phylogeny of swallows (Aves: Hirundinidae) estimated from nuclear and mitochondrial DNA sequences. Molecular Phylogenetics and Evolution 35 (1): 254-270. 1

Lovette, Irby J., Brian S. Arbogast, Robert L. Curry, Robert M. Zink, Carlos A. Botero, John P. Sullivan, Amanda L. Talaba, Rebecca B. Harris, Dustin R. Rubenstein, Robert E. Ricklefs, Eldredge Bermingham. 2012. Phylogenetic relationships of the mockingbirds and thrashers (Aves: Mimidae). Molecular Phylogenetics and Evolution 63 (2): 219-229. 1

Lovette, Irby J., Dustin R. Rubenstein. 2007. A comprehensive molecular phylogeny of the starlings (Aves: Sturnidae) and mockingbirds (Aves: Mimidae): congruent mtDNA and nuclear trees for a cosmopolitan avian radiation. Molecular Phylogenetics and Evolution 44 (3): 1031-1056. 1

Alström, P., Mohammadi, Z., Enbody, E., Irestedt, M., Engelbrecht, D., Crochet, P.-A., Guillaumet, A., Rancilhac, L., Tieleman, B. I., Olsson, U., Donald, P. F., & Stervander, M. (2023). Systematics of the avian family Alaudidae using multilocus and genomic data. Avian Research, 100095. https://doi.org/10.1016/j.avrs.2023.100095 1

Gibb, Gillian C., Ryan England, Gerrit Hartig, P.A. (Trish) McLenachan, Briar L. Taylor Smith, Bennet J. McComish, Alan Cooper, David Penny. 2015. New Zealand passerines help clarify the diversification of major songbird lineages during the Oligocene. Genome Biology and Evolution 7 (11): 2983-2995. 1

Ottenburghs, Jente, Hendrik-Jan Megens, Robert H.S. Kraus, Ole Madsen, Pim van Hooft, Sipke E. van Wieren, Richard P.M.A. Crooijmans, Ronald C. Ydenberg, Martien A.M. Groenen, Herbert H.T. Prins. 2016. A Tree of Geese: A Phylogenomic Perspective on the Evolutionary History of True Geese. Molecular Phylogenetics and Evolution 101: 303-313. 1

Ericson, Per GP, Seraina Klopfstein, Martin Irestedt, Jacqueline MT Nguyen and Johan AA Nylander. 2014. Dating the diversification of the major lineages of Passeriformes (Aves). BMC Evolutionary Biology, 14(8). 1

Beresford, P., F.K. Barker, P.G. Ryan, T.M. Crowe. 2005. African endemics span the tree of songbirds (Passeri): molecular systematics of several evolutionary 'enigmas'. Proceedings of the Royal Society B: Biological Sciences 272 (1565): 849-858 1

Fuchs, Jérôme, Jan I. Ohlson, Per G. P. Ericson, Eric Pasquet. 2007. Synchronous intercontinental splits between assemblages of woodpeckers suggested by molecular data. Zoologica Scripta 36 (1): 11-25 1

Zuccon, Dario, Alice Cibois, Eric Pasquet, Per G.P. Ericson. 2006. Nuclear and mitochondrial sequence data reveal the major lineages of starlings, mynas and related taxa. Molecular Phylogenetics and Evolution 41 (2): 333-344 1

Provost, K. L., Joseph, L., & Smith, B. T. (2018). Resolving a phylogenetic hypothesis for parrots: implications from systematics to conservation. Emu - Austral Ornithology, 118(1), 7–21. doi:10.1080/01584197.2017.1387030 0.9

eBird Taxonomy draft, 2021, 2023 0.9

Oliver, P. M., Heiniger, H., Hugall, A. F., Joseph, L., & Mitchell, K. J. (2020). Oligocene divergence of frogmouth birds (Podargidae) across Wallace's Line. Biology Letters, 16(5), 20200040. https://doi.org/10.1098/rsbl.2020.0040 0.9

Pietersen, D. W., McKechnie, A. E., Jansen, R., Little, I. T., & Bastos, A. D. S. (2018). Multi-locus phylogeny of African pipits and longclaws (Aves: Motacillidae) highlights taxonomic inconsistencies. Ibis, 161(4), 781–792. Portico. https://doi.org/10.1111/ibi.12683 0.9

Lapiedra, O., Sayol, F., Garcia-Porta, J., & Sol, D. (2021). Niche shifts after island colonization spurred adaptive diversification and speciation in a cosmopolitan bird clade. Proceedings of the Royal Society B: Biological Sciences, 288(1958), 20211022. https://doi.org/10.1098/rspb.2021.1022 0.9

Dos Remedios, Natalie, Patricia L.M. Lee, Terry Burke, Tamás Székely, Clemens Küpper. 2015. North or south? Phylogenetic and biogeographic origins of a globally distributed avian clade. Molecular Phylogenetics and Evolution 89:151-159. 0.9

Slager, David L., C.J. Battey, Robert W. Bryson, Gary Voelker, John Klicka. 2014. A multilocus phylogeny of a major New World avian radiation: The Vireonidae. Molecular Phylogenetics and Evolution 80: 95-104 0.9

Alström, Per, Keith N. Barnes, Urban Olsson, F. Keith Barker, Paulette Bloomer, Aleem Ahmed Khan, Masood Ahmed Qureshi, Alban Guillaumet, Pierre-André Crochet, Peter G. Ryan. 2013. Multilocus phylogeny of the avian family Alaudidae (larks) reveals complex morphological evolution, non-monophyletic genera and hidden species diversity. Molecular Phylogenetics and Evolution 69 (3): 1043-1056. 0.9

Arbabi, Tayebeh, Javier Gonzalez, Michael Wink. 2014. A re-evaluation of phylogenetic relationships within reed warblers (Aves: Acrocephalidae) based on eight molecular loci and ISSR profiles. Molecular Phylogenetics and Evolution 78: 304-313. 0.7

Kennedy, J. D., Marki, P. Z., Reeve, A. H., Blom, M. P. K., Prawiradilaga, D. M., Haryoko, T., Koane, B., Kamminga, P., Irestedt, M., Jønsson, K. A., & Sheard, C. (2022). Diversification and community assembly of the world's largest tropical island. Global Ecology and Biogeography. Portico. https://doi.org/10.1111/geb.13484 0.7

Hooper, Daniel M., Urban Olsson, Per Alström. 2016. The Rusty-tailed Flycatcher (Muscicapa ruficauda; Aves: Muscicapidae) is a member of the genus Ficedula. Molecular Phylogenetics and Evolution 102: 56-61 0.7

Voelker, G., & Light J.E. 2011. Palaeoclimatic events, dispersal and migratory losses along the Afro-European axis as drivers of biogeographic distribution in Sylvia warblers. BMC Evolutionary Biology 11: 163. 0.7

Han, Kin-Lan, Mark B. Robbins, Michael J. Braun. 2010. A multi-gene estimate of phylogeny in the nightjars and nighthawks (Caprimulgidae). Molecular Phylogenetics and Evolution 55 (2): 443-453. 0.7

Peter A. Hosner, Edward L. Braun, Rebecca T. Kimball, 2015, 'Land connectivity changes and global cooling shaped the colonization history and diversification of New World quail (Aves: Galliformes: Odontophoridae)', Journal of Biogeography, vol. 42, no. 10, pp. 1883-1895 0.6

Per Alström, Frank E. Rheindt, Ruiying Zhang, Min Zhao, Jing Wang, Xiaojia Zhu, Chyi Yin Gwee, Yan Hao, Jan Ohlson, Chenxi Jia, Dewi M. Prawiradilaga, Per G.P. Ericson, Fumin Lei, Urban Olsson, 2018, 'Complete species-level phylogeny of the leaf warbler (Aves: Phylloscopidae) radiation', Molecular Phylogenetics and Evolution, vol. 126, pp. 141-152 0.6

Benz, Brett W., Mark B. Robbins, A. Townsend Peterson. 2006. Evolutionary history of woodpeckers and allies (Aves: Picidae): placing key taxa on the phylogenetic tree. Molecular Phylogenetics and Evolution 40 (2): 389-399. 0.6

Batista, R., Olsson, U., Andermann, T., Aleixo, A., Ribas, C. C., & Antonelli, A. (2020). Phylogenomics and biogeography of the world’s thrushes (Aves, Turdus): new evidence for a more parsimonious evolutionary history. Proceedings of the Royal Society B: Biological Sciences, 287(1919), 20192400. https://doi.org/10.1098/rspb.2019.2400 0.6

Garcia-R, J. C., Lemmon, E. M., Lemmon, A. R., & French, N. (2020). Phylogenomic Reconstruction Sheds Light on New Relationships and Timescale of Rails (Aves: Rallidae) Evolution. Diversity, 12(2), 70. https://doi.org/10.3390/d12020070 0.6

Salter, J. F., Hosner, P. A., Tsai, W. L. E., McCormack, J. E., Braun, E. L., Kimball, R. T., Brumfield, R. T., & Faircloth, B. C. (2022). Historical specimens and the limits of subspecies phylogenomics in the New World quails (Odontophoridae). Molecular Phylogenetics and Evolution, 175, 107559. https://doi.org/10.1016/j.ympev.2022.107559 0.6

Zhou, Xiaoping, Chengte Yao, Qingxian Lin, Wenzhen Fang, Xiaolin Chen. 2015. Complete mitochondrial genomes render the Night Heron genus Gorsachius non-monophyletic. Journal of Ornithology 157 (2): 505-513 0.6

Jérôme Fuchs, Jean-Marc Pons, Steven M Goodman, Vincent Bretagnolle, Martim Melo, Rauri CK Bowie, David Currie, Roger Safford, Munir Z Virani, Simon Thomsett, Alawi Hija, Corinne Cruaud, Eric Pasquet, 2008, 'Tracing the colonization history of the Indian Ocean scops-owls (Strigiformes: Otus) with further insight into the spatio-temporal origin of the Malagasy avifauna', BMC Evolutionary Biology, vol. 8, no. 1, p. 197 0.6

Zink, Robert M., Donna L. Dittmann, John Klicka, Rachelle C. Blackwell-Rago. 1999. Evolutionary patterns of morphometrics, allozymes, and mitochondrial DNA in thrashers (genus Toxostoma). The Auk 116 (4): 1021-1038 0.6

Päckert, Martin, Jochen Martens, Michael Wink, Anna Feigl, Dieter Thomas Tietze. 2012. Molecular phylogeny of Old World swifts (Aves: Apodiformes, Apodidae, Apus and Tachymarptis) based on mitochondrial and nuclear markers. Molecular Phylogenetics and Evolution 63 (3): 606-616. 0.5

Payne, Robert B. The cuckoos. Vol. 15. Oxford University Press, 2005.
ISBN-13: 978-0198502135 ISBN-10: 0198502133 0.5

Alström, P., Jønsson, K. A., Fjeldså, J., Ödeen, A., Ericson, P. G. P., & Irestedt, M. (2015). Dramatic niche shifts and morphological change in two insular bird species. Royal Society Open Science, 2(3), 140364. https://doi.org/10.1098/rsos.140364 0.5

García–R, Juan C., Gillian C. Gibb, Steve A. Trewick. 2014. Eocene diversification of crown group rails (Aves: Gruiformes: Rallidae). PLoS ONE 9 (10): e109635 0.5

Mann, Nigel I., F. Keith Barker, Jeff A. Graves, Kimberly A. Dingess-Mann, Peter J.B. Slater. 2006. Molecular data delineate four genera of “Thryothorus” wrens. Molecular Phylogenetics and Evolution 40 (3): 750-759 0.5

Wright, T. F., E. E. Schirtzinger, T. Matsumoto, J. R. Eberhard, G. R. Graves, J. J. Sanchez, S. Capelli, H. Muller, J. Scharpegge, G. K. Chambers, R. C. Fleischer. 2008. A multilocus molecular phylogeny of the parrots (Psittaciformes): support for a Gondwanan origin during the Cretaceous. Molecular Biology and Evolution 25 (10): 2141-2156. 0.5

Ramirez, J.L., C.Y. Miyaki, S.N. Del Lama. 2013. Molecular phylogeny of Threskiornithidae (Aves: Pelecaniformes) based on nuclear and mitochondrial DNA. Genetics and Molecular Research 12 (3): 2740-2750. 0.4

Harris, R.B., M.D. Carling, I.J. Lovette. 2014. The influence of sampling design on species tree inference: a new relationship for the New World chickadees (Aves: Poecile). Evolution 68 (2): 501-513. 0.4

Stervander, M., Hansson, B., Olsson, U., Hulme, M. F., Ottosson, U., & Alström, P. (2020). Molecular Species Delimitation of Larks (Aves: Alaudidae), and Integrative Taxonomy of the Genus Calandrella, with the Description of a Range-Restricted African Relic Taxon. Diversity, 12(11), 428. https://doi.org/10.3390/d12110428 0.4

Gibb, G. C., & Shepherd, L. D. (2022). Recent evolution of extreme sexual dimorphism in the huia (Heteralocha acutirostris; Callaeidae). Molecular Phylogenetics and Evolution, 175, 107575. https://doi.org/10.1016/j.ympev.2022.107575 0.4

Olsson, U., & Alström, P. (2020). A comprehensive phylogeny and taxonomic evaluation of the waxbills (Aves: Estrildidae). Molecular Phylogenetics and Evolution, 146, 106757. https://doi.org/10.1016/j.ympev.2020.106757 0.4

Oatley, Graeme, Robert E. Simmons, Jérôme Fuchs. 2015. A molecular phylogeny of the harriers (Circus, Accipitridae) indicate the role of long distance dispersal and migration in diversification. Molecular Phylogenetics and Evolution 85:150-160. 0.4

| | |
|---|---|
| Pasquet, Eric, F. Keith Barker, Jochen Martens, Annie Tillier, Corinne Cruaud, Alice Cibois. 2014. Evolution within the nuthatches (Sittidae: Aves, Passeriformes): molecular phylogeny, biogeography, and ecological perspectives. Journal of Ornithology 155 (3):755-765. | 0.4 |
| Nylander, Johan A. A., Urban Olsson, Per Alström, Isabel Sanmartín. 2008. Accounting for phylogenetic uncertainty in biogeography: a Bayesian approach to dispersal-vicariance analysis of the thrushes (Aves: Turdus). Systematic Biology 57 (2): 257-268. | 0.4 |
| Soares, André E. R. , Ben J. Novak, James Haile, Tim H. Heupink, Jon Fjeldså, M. Thomas P. Gilbert, Hendrik Poinar, George M. Church, Beth Shapiro. 2016. Complete mitochondrial genomes of living and extinct pigeons revise the timing of the columbiform radiation. BMC Evolutionary Biology 16: 230 | 0.2 |
| Andersen, Michael J., Jenna M. McCullough, William M. Mauck, Brian Tilston Smith, Robert G. Moyle. 2017. A phylogeny of kingfishers reveals an Indomalayan origin and elevated rates of diversification on oceanic islands. Journal of Biogeography | 0.2 |
| Brian Tilston Smith, Robert W. Bryson, William M. Mauck, Jaime Chaves, Mark B. Robbins, Alexandre Aleixo, John Klicka, 2018, 'Species delimitation and biogeography of the gnatcatchers and gnatwrens (Aves: Polioptilidae)', Molecular Phylogenetics and Evolution, vol. 126, pp. 45-57 | 0.2 |
| Robert W. Bryson, Jaime Chaves, Brian Tilston Smith, Matthew J. Miller, Kevin Winker, Jorge L. Pérez-Emán, John Klicka, 2014, 'Diversification across the New World within the 'blue' cardinalids (Aves: Cardinalidae)', Journal of Biogeography, vol. 41, no. 3, pp. 587-599 | 0.2 |
| Päckert, Martin , Jochen Martens, Lucia Liu Severinghaus. 2009. The Taiwan Firecrest (Regulus goodfellowi) belongs to the Goldcrest assemblage (Regulus regulus s. l.): evidence from mitochondrial DNA and the territorial song of the Regulidae. Journal of Ornithology 150 (1): 205-220. | 0.2 |
| Yonezawa, T., Segawa, T., Mori, H., Campos, P. F., Hongoh, Y., Endo, H., Akiyoshi, A., Kohno, N., Nishida, S., Wu, J., Jin, H., Adachi, J., Kishino, H., Kurokawa, K., Nogi, Y., Tanabe, H., Mukoyama, H., Yoshida, K., Rasoamiaramanana, A., … Hasegawa, M. (2017). Phylogenomics and Morphology of Extinct Paleognaths Reveal the Origin and Evolution of the Ratites. Current Biology, 27(1), 68–77. https://doi.org/10.1016/j.cub.2016.10.029 | 0.2 |
| White, N. D., & Braun, M. J. (2019). Extracting phylogenetic signal from phylogenomic data: Higher-level relationships of the nightbirds (Strisores). Molecular Phylogenetics and Evolution, 141, 106611. https://doi.org/10.1016/j.ympev.2019.106611 | 0.2 |
| McCullough, J. M., Moyle, R. G., Smith, B. T., & Andersen, M. J. (2019). A Laurasian origin for a pantropical bird radiation is supported by genomic and fossil data (Aves: Coraciiformes). Proceedings of the Royal Society B: Biological Sciences, 286(1910), 20190122. https://doi.org/10.1098/rspb.2019.0122 | 0.2 |
| Hosner, P. A., Braun, E. L., & Kimball, R. T. (2016). Rapid and recent diversification of curassows, guans, and chachalacas (Galliformes: Cracidae) out of Mesoamerica: Phylogeny inferred from mitochondrial, intron, and ultraconserved element sequences. Molecular Phylogenetics and Evolution, 102, 320–330. https://doi.org/10.1016/j.ympev.2016.06.006 | 0.2 |
| Alström, P., Cibois, A., Irestedt, M., Zuccon, D., Gelang, M., Fjeldså, J., Andersen, M. J., Moyle, R. G., Pasquet, E., & Olsson, U. (2018). Comprehensive molecular phylogeny of the grassbirds and allies (Locustellidae) reveals extensive non-monophyly of traditional genera, and a proposal for a new classification. Molecular Phylogenetics and Evolution, 127, 367–375. https://doi.org/10.1016/j.ympev.2018.03.029 | 0.2 |
| Dantas, Sidnei M., Jason D. Weckstein, John M. Bates, Niels K. Krabbe, Carlos Daniel Cadena, Mark B. Robbins, Eugenio Valderrama, Alexandre Aleixo. 2016. Molecular systematics of the new world screech-owls (Megascops: Aves, Strigidae): biogeographic and taxonomic implications. Molecular Phylogenetics and Evolution 94: 626-634 | 0.2 |
| Besnard, Guillaume, Joris A. M. Bertrand, Boris Delahaie, Yann X. C. Bourgeois, Emeline Lhuillier, Christophe Thébaud. 2015. Valuing museum specimens: high-throughput DNA sequencing on historical collections of New Guinea crowned pigeons (Goura). Biological Journal of the Linnean Society 117 (1): 71-82 | 0.2 |
| Robert G. Moyle, 2006, 'A MOLECULAR PHYLOGENY OF KINGFISHERS (ALCEDINIDAE) WITH INSIGHTS INTO EARLY BIOGEOGRAPHIC HISTORY', The Auk, vol. 123, no. 2, p. 487 | 0.2 |
| Rheindt, Frank E., Janette A. Norman, Les Christidis. 2008. Phylogenetic relationships of tyrant-flycatchers (Aves: Tyrannidae), with an emphasis on the elaeniine assemblage. Molecular Phylogenetics and Evolution 46 (1): 88-101 | 0.2 |
| Fain, M., Krajewski C., & Houde P. 2007. Phylogeny of "core Gruiformes" (Aves: Grues) and resolution of the Limpkin-Sungrebe problem. Molecular Phylogenetics and Evolution 43 (2): 515-529. | 0.2 |
| Joseph, Leo, Alicia Toon, Erin E. Schirtzinger, Timothy F. Wright. 2011. Molecular systematics of two enigmatic genera Psittacella and Pezoporus illuminate the ecological radiation of Australo-Papuan parrots (Aves: Psittaciformes). Molecular Phylogenetics and Evolution 59 (3): 675-684. | 0.2 |
| Shakya, Subir B., Frederick H. Sheldon. 2017. The phylogeny of the world's bulbuls (Pycnonotidae) inferred using a supermatrix approach. Ibis 159 (3): 498-509 | 0.1 |
| Pitra, Christian, Dietmar Lieckfeldt, Sylke Frahnert, Joerns Fickel. 2002. Phylogenetic relationships and ancestral areas of the bustards (Gruiformes: Otididae), inferred from mitochondrial DNA and nuclear intron sequences. Molecular Phylogenetics and Evolution 23 (1): 63-74. | 0.1 |
| Krajewski, Carey, Justin T. Sipiorski, Frank E. Anderson. 2010. Complete mitochondrial genome sequences and the phylogeny of cranes (Gruiformes: Gruidae). The Auk 127 (2): 440-452. | 0.1 |
| Slikas, Beth. 1997. Phylogeny of the avian family Ciconiidae (storks) based on cytochrome b sequences and DNA–DNA hybridization distances. Molecular Phylogenetics and Evolution 8 (3): 275-300. | 0.1 |

| | |
|---|---|
| Kennedy, Martyn, Hamish G. Spencer. 2014. Classification of the cormorants of the world. Molecular Phylogenetics and Evolution 79: 249-257. | 0.1 |
| Russello, Michael A, George Amato. 2004. A molecular phylogeny of Amazona: implications for Neotropical parrot biogeography, taxonomy, and conservation. Molecular Phylogenetics and Evolution 30 (2): 421-437. | 0.1 |
| Ribas, Camila C., Cristina Y. Miyaki, Joel Cracraft. 2009. Phylogenetic relationships, diversification and biogeography in Neotropical Brotogeris parakeets. Journal of Biogeography 36 (9): 1712-1729. | 0.1 |
| Päckert, Martin, Jochen Martens, Yue-Hua Sun, Lucia Liu Severinghaus, Alexander A. Nazarenko, Ji Ting, Till Töpfer, Dieter Thomas Tietze. 2011. Horizontal and elevational phylogeographic patterns of Himalayan and Southeast Asian forest passerines (Aves: Passeriformes). Journal of Biogeography 39 (3): 556-573. | 0.1 |
| Imfeld, Barker, Brumfield, Mitochondrial genomes and thousands of ultraconserved elements resolve the taxonomy and historical biogeography of the Euphonia and Chlorophonia finches (Passeriformes: Fringillidae), The Auk, Volume 137, Issue 3, 1 July 2020, ukaa016, | 0.1 |
| Nováková, N., & Robovský, J. (2021). Behaviour of cranes (family Gruidae) mirrors their phylogenetic relationships. Avian Research, 12(1). https://doi.org/10.1186/s40657-021-00275-4 | 0.1 |
| Nagy, J., Végvári, Z., & Varga, Z. (2019). Phylogeny, migration and life history: filling the gaps in the origin and biogeography of the Turdus thrushes. Journal of Ornithology, 160(2), 529–543. https://doi.org/10.1007/s10336-019-01632-3 | 0.1 |
| Jha, A., Seneviratne, S., Prayag, H. S., & Vasudevan, K. (2021). Phylogeny identifies multiple colonisation events and Miocene aridification as drivers of South Asian bulbul (Passeriformes: Pycnonotidae) diversification. Organisms Diversity & Evolution, 21(4), 783–794. https://doi.org/10.1007/s13127-021-00506-y | 0.1 |
| Vázquez-López, M., Ramírez-Barrera, S. M., Terrones-Ramírez, A. K., Robles-Bello, S. M., Nieto-Montes de Oca, A., Ruegg, K., & Hernández-Baños, B. E. (2024). Biogeographic factors contributing to the diversification of Euphoniinae (Aves, Passeriformes, Fringillidae): a phylogenetic and ancestral areas analysis. ZooKeys, 1188, 169–195. https://doi.org/10.3897/zookeys.1188.107047 | 0.1 |
| Sweet, Andrew D., Kevin P. Johnson. 2015. Patterns of diversification in small New World ground doves are consistent with major geologic events. The Auk 132 (1): 300-312. | 0.1 |
| Johnson, Jeff A., Joseph W. Brown, Jérôme Fuchs, David P. Mindell, 2016, 'Multi-locus phylogenetic inference among New World Vultures (Aves: Cathartidae)', Molecular Phylogenetics and Evolution, vol. 105, pp. 193-199 | 0.1 |
| Paxinos, E. E. , H. F. James, S. L. Olson, M. D. Sorenson, J. Jackson, R. C. Fleischer. 2002. mtDNA from fossils reveals a radiation of Hawaiian geese recently derived from the Canada goose (Branta canadensis). Proceedings of the National Academy of Sciences 99 (3): 1399-1404 | 0.1 |
| Pasquet, Eric. 2008. Phylogeny of the nuthatches of the Sitta canadensis group and its evolutionary and biogeographic implications. Ibis 140 (1): 150-156 | 0.1 |
| Robert G. Moyle, Joel Cracraft, Maklarin Lakim, Jamili Nais, Frederick H. Sheldon, 2006, 'Reconsideration of the phylogenetic relationships of the enigmatic Bornean Bristlehead (Pityriasis gymnocephala)', Molecular Phylogenetics and Evolution, vol. 39, no. 3, pp. 893-898 | 0.1 |
| Irby J. Lovette, Brynn V. McCleery, Amanda L. Talaba, Dustin R. Rubenstein, 2008, 'A complete species-level molecular phylogeny for the “Eurasian” starlings (Sturnidae: Sturnus, Acridotheres, and allies): Recent diversification in a highly social and dispersive avian group', Molecular Phylogenetics and Evolution, vol. 47, no. 1, pp. 251-260 | 0.1 |
| Juan Carlos Illera, David S. Richardson, Barbara Helm, Juan Carlos Atienza, Brent C. Emerson, 2008, 'Phylogenetic relationships, biogeography and speciation in the avian genus Saxicola', Molecular Phylogenetics and Evolution, vol. 48, no. 3, pp. 1145-1154 | 0.1 |
| Zink, Robert M., Rachelle C. Blackwell. 1998. Molecular systematics and biogeography of aridland gnatcatchers (genus Polioptila) and evidence supporting species status of the California gnatcatcher (Polioptila californica). Molecular Phylogenetics and Evolution 9 (1): 26-32 | 0.1 |
| Aliabadian, Mansour, Mohammad Kaboli, Roger Prodon, Vincent Nijman, Miguel Vences. 2007. Phylogeny of Palaearctic wheatears (genus Oenanthe)—Congruence between morphometric and molecular data. Molecular Phylogenetics and Evolution 42 (3): 665-675 | 0.1 |
| Moyle, R.G., Andersen M.J., Oliveros C., Steinheimer F.D., and Reddy S. 2012. Phylogeny and biogeography of the core babblers (Aves: Timaliidae). Systematic Biology 61 (4): 631-651. | 0.1 |
| Kennedy, Martyn, Carlos A. Valle, Hamish G. Spencer. 2009. The phylogenetic position of the Galapagos Cormorant. Molecular Phylogenetics and Evolution 53 (1): 94-98. | 0.1 |
| Pereira, S., Johnson K., Clayton D., and Baker A. 2007. Mitochondrial and nuclear DNA sequences support a Cretaceous origin of Columbiformes and a dispersal-driven radiation in the Paleogene. Systematic Biology 56 (4): 656-672. | 0.1 |
| Fuchs, Jérôme, Steven Chen, Jeff A. Johnson, David P. Mindell. 2011. Pliocene diversification within the South American forest falcons (Falconidae: Micrastur). Molecular Phylogenetics and Evolution 60 (3): 398-407. | 0.1 |
| Fuchs, Jérôme, Jeff A. Johnson, David P. Mindell. 2012. Molecular systematics of the caracaras and allies (Falconidae: Polyborinae) inferred from mitochondrial and nuclear sequence data. Ibis 154 (3): 520-532. | 0.1 |

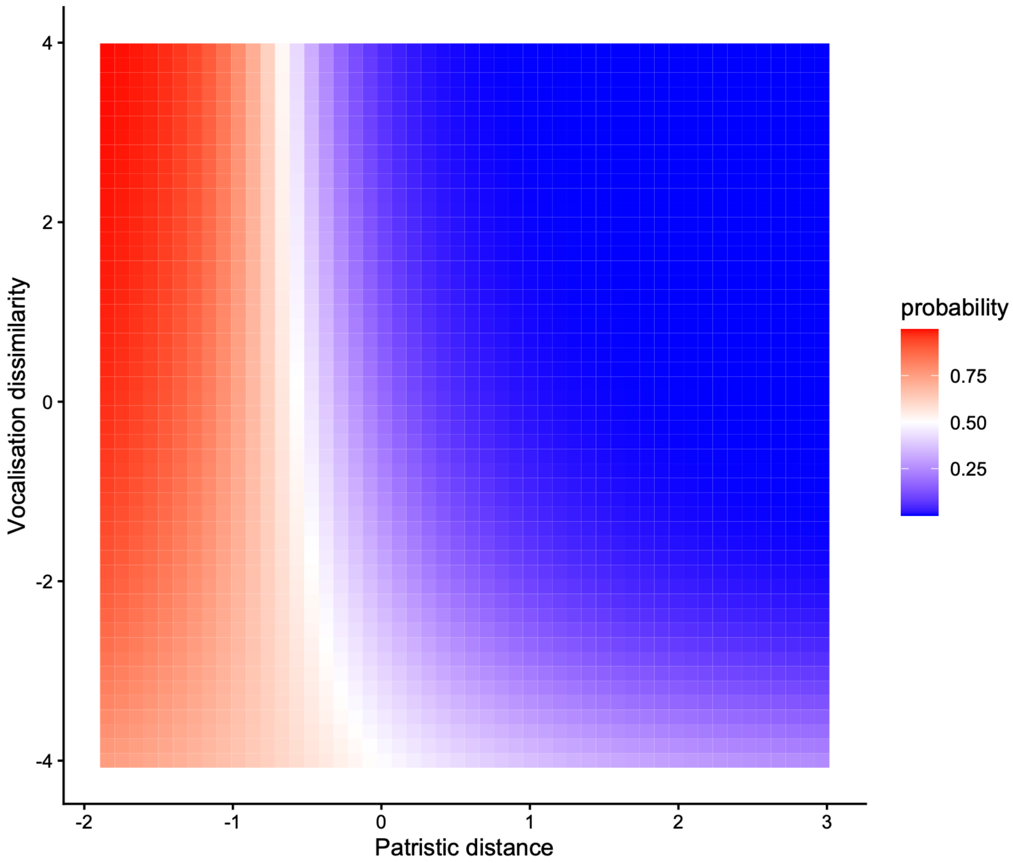

Supplementary Figure 1. Heatmap depicting the predicted probability of hybridisation as a function of the interaction between patristic distance and vocalisation dissimilarity from the non-passerine interaction model presented in Supplementary Table 8. NB: axes represent scaled values.

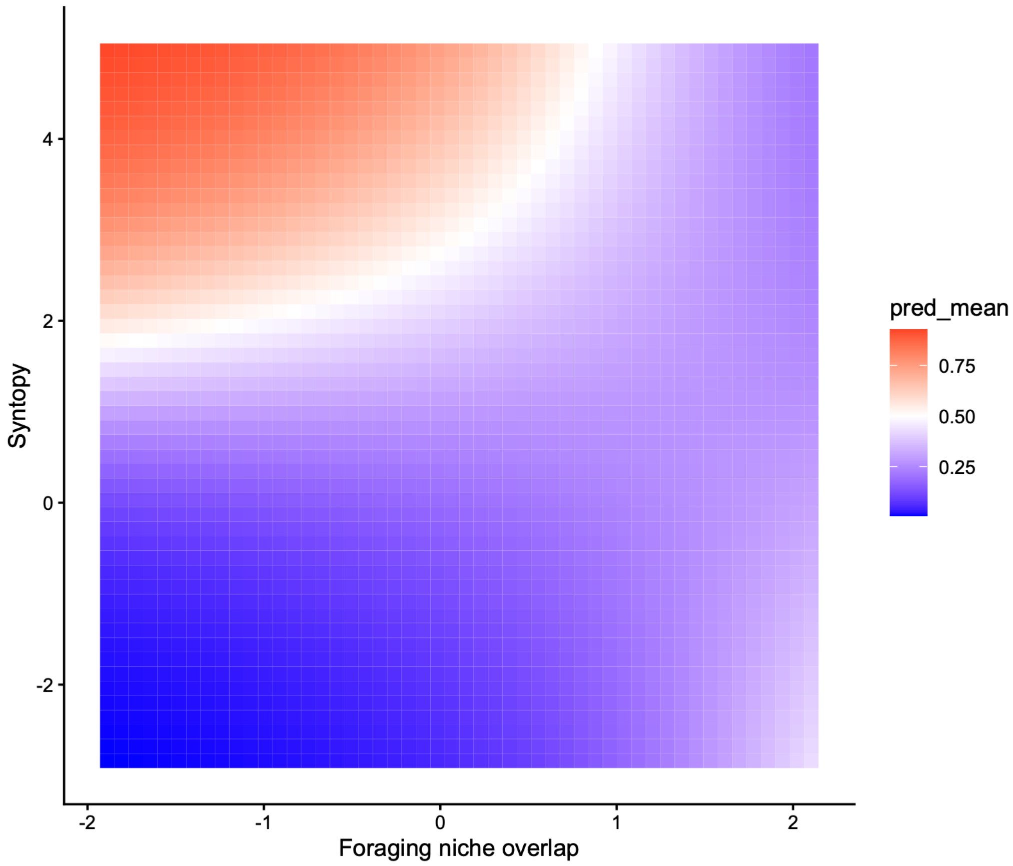

Supplementary Figure 2. Heatmap depicting the predicted probability of hybridisation as a function of the interaction between foraging niche overlap and syntopy from the non-passerine interaction model presented in Supplementary Table 8. NB: axes represent scaled values.

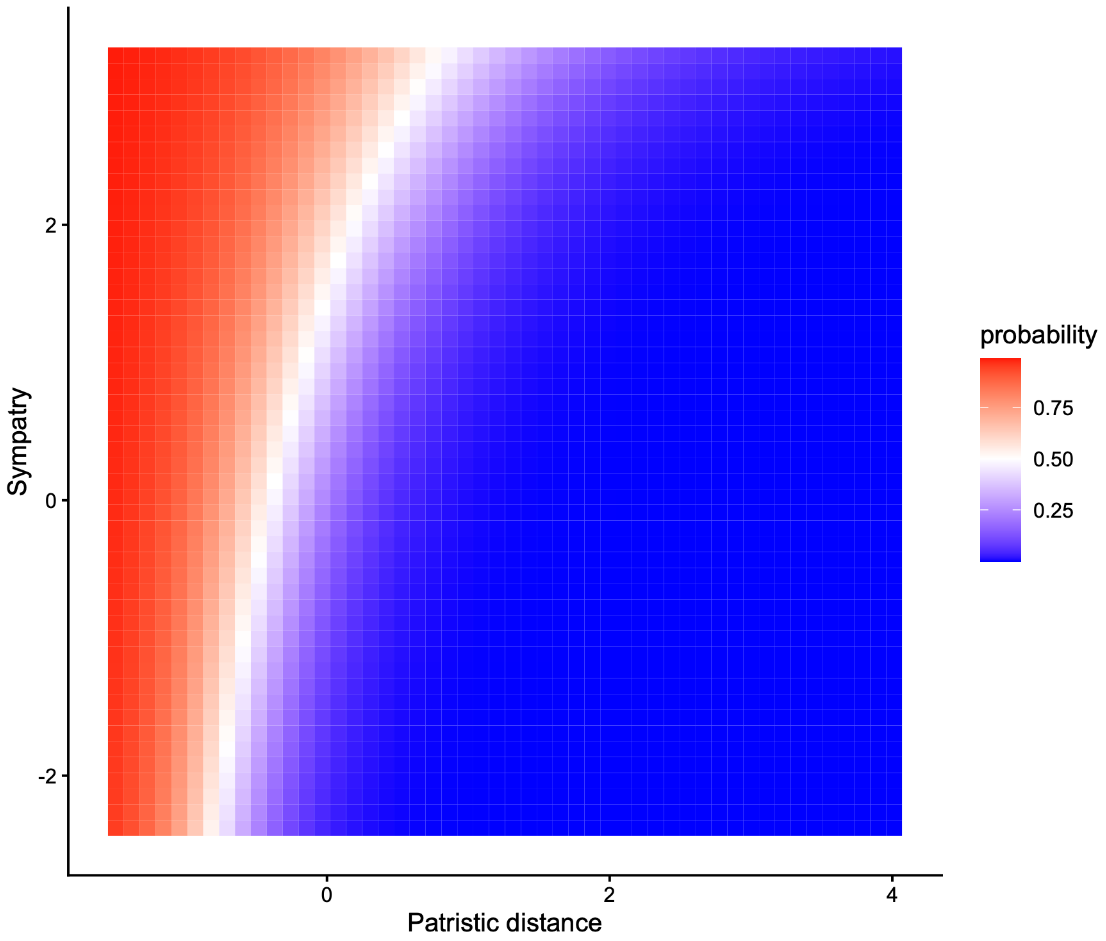

Supplementary Figure 3. Heatmap depicting the predicted probability of hybridisation as a function of the interaction between patristic distance and sympatry from the full interaction model presented in Supplementary Table 6. NB: axes represent scaled values.

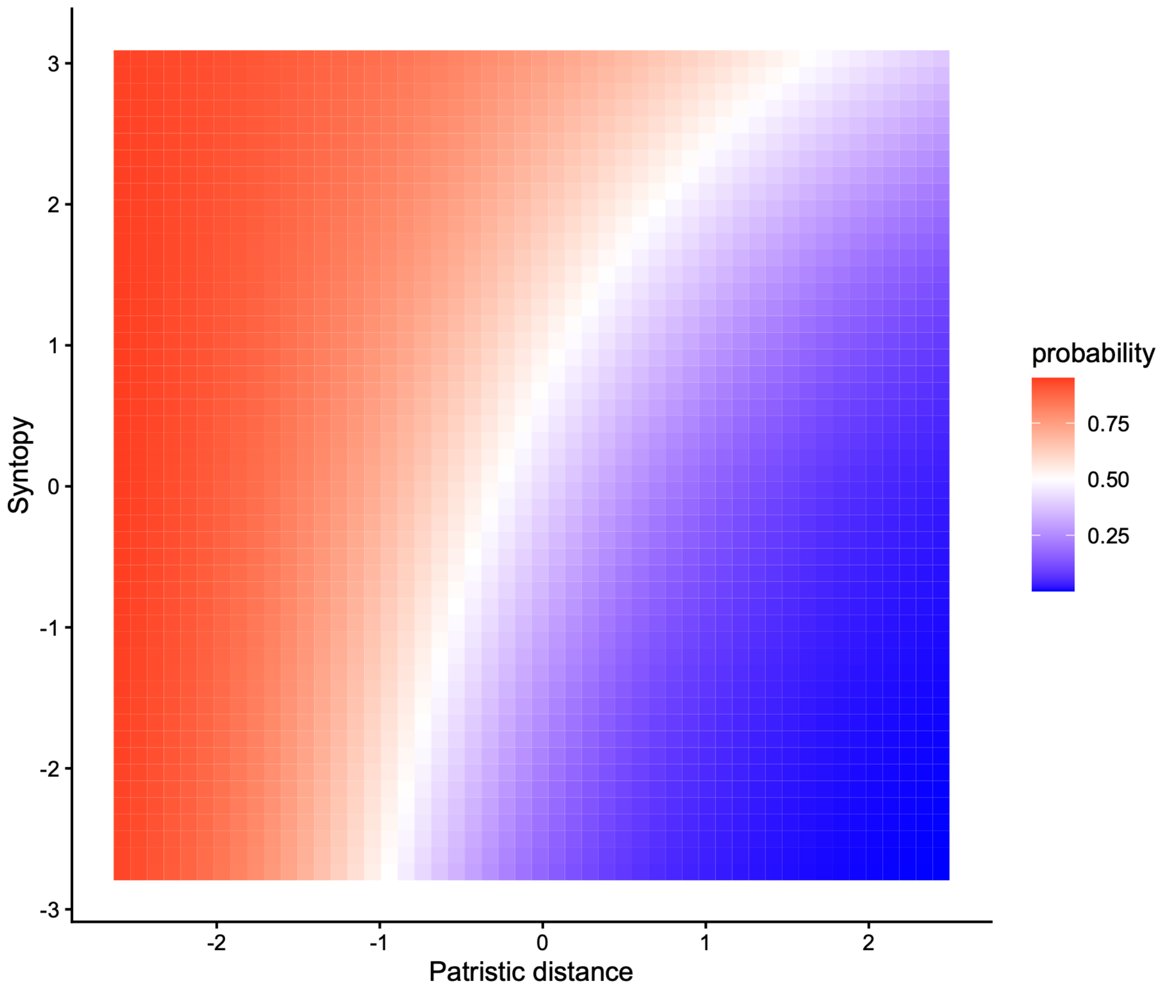

Supplementary Figure 4. Heatmap depicting the predicted probability of hybridisation as a function of the interaction between foraging niche overlap and syntopy from the passerine interaction model presented in Supplementary Table 7. NB: axes represent scaled values.

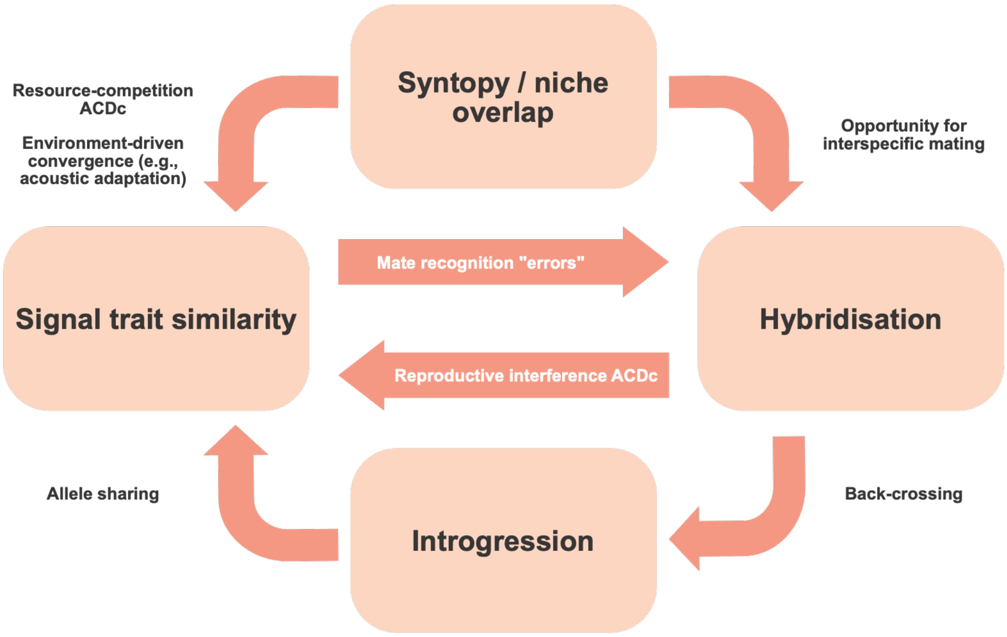

Supplementary Figure 5. Potential causal pathways linking trait similarity in signal traits and hybridisation in birds.
